# ARF6 controls VSMC cell phenotypic switching upon lipid stimulation to promote inflammatory signaling contributing to the progression of atherosclerosis

**DOI:** 10.1101/2025.09.16.676422

**Authors:** Fiola-Masson Émilie, Shirley Campbell, Véronique Laplante, Rania Lejri, Yasunori Kanaho, Jean-François Gauchat, Marc J. Servant, Audrey Claing

## Abstract

Vascular smooth muscle cells (VSMCs) play a pivotal role in the development and progression of atherosclerosis. Traditionally viewed as contractile cells that maintain vascular tone and structure, VSMCs undergo phenotypic switching in response to atherogenic stimuli, such as high circulating levels of LDL, thus adopting synthetic, osteogenic, or macrophage-like phenotypes. This plasticity contributes to plaque formation, extracellular matrix remodeling, and inflammatory signaling. We have previously shown that ADP-ribosylation factor 1 (ARF1), a small GTP-binding protein, regulates the expression and function of actin, which is important for maintaining the contractile phenotype of VSMCs. However, the role of ARF6 in phenotypic switching remains to be elucidated. Here, we demonstrate that ARF6 knockdown in human aortic smooth muscle cells (HASMCs) reduced lipid uptake through alterations in the expression of scavenger receptors (LOX-1, MSR1), cytokine production (IL-6) as well as modulation of inflammatory markers and pathways (adhesion molecules, PI3K, NFkB, p38). To confirm our findings in an *in vivo* setting, we engineered a novel conditional smooth muscle cell-specific ARF6 KO mouse in an atheroprone background (*Acta2-Cre-ERT2^+/−^/ApoE^−/−^/Arf6^f/f^* mice). Mice were fed a high-fat diet to accelerate plaque formation. ARF6 knockout resulted in a significant reduction of atherosclerotic lesions in the aortic arch, which was associated with a reduction of collagen and foam cells. Furthermore, we observed that ARF6 regulates the expression of inflammatory markers. These findings highlight the importance of ARF6 expression in VSMCs and its role in the pathogenesis of atherosclerosis.

## Introduction

Vascular smooth muscle cells (VSMCs) play a crucial role in maintaining the structural integrity and the function of blood vessels, and thus directly impact blood pressure and blood flow. These cells respond to a variety of stimuli ranging from circulating hormones and lipids, to neurotransmitters and mechanical stress. In a normal physiological state, they exhibit a contractile phenotype and are referred to as differentiated VSMCs. However, they possess the ability to adapt and change their phenotype in response to stimulation. This plasticity between a differentiated (or contractile) phenotype and a dedifferentiated (or synthetic) phenotype is critical for healing, but can also contribute to vascular diseases, namely atherosclerosis, if unregulated.

Atherosclerosis is characterized by the accumulation of cholesterol coupled to chronic inflammation and excessive production of extracellular matrix (ECM) within the inner wall of arteries, leading to plaque formation and stabilization. When contractile VSMCs undergo phenotypic switching, they can proliferate, migrate and produce ECM proteins. They may also develop characteristics reminiscent of many cell types, including mesenchymal cells, fibroblasts, osteoblasts, macrophages and foam cells (1). Recent studies have shown that macrophage-like VSMCs acquire macrophagic markers, making them difficult to distinguish from myeloid-derived macrophages (2). Interestingly, it was suggested that in mouse models of atherosclerosis, almost 70% of the foam cells were of VSMC, rather than macrophage, origin (3). During atherosclerosis, low-density lipoprotein (LDL) particles infiltrate the subendothelial space of blood vessels and can undergo chemical modification, such as oxidation. When oxidized lipids (OxLDL) are present, VSMCs express cholesterol receptors to mediate lipid uptake, as well as pro-inflammatory markers and cytokines (4). Intracellular OxLDL accumulation drives foam cell formation through numerous mechanisms, including activation of transcription factors such as the Kruppel-like factor 4 (KLF4), which lead to altered gene expression and intracellular signaling cascades (5). Foam cells contribute to the aggravation of the pro-inflammatory environment surrounding atherosclerotic lesions by continually secreting cytokines, thereby further promoting disease progression. Eventually, this inflammatory imbalance causes VSMC apoptosis and necrosis, which weakens the stability of the lesions and leads to plaque rupture (6, 7).

Because ADP-ribosylation factors (ARF) are key regulators of vascular smooth muscle cell function (8–10) and monomeric G proteins in general have emerged as key players in regulating a diverse set of biological activities, we speculated that ARF proteins could act as molecular switches to regulate the dedifferentiation of VSMCs when in the presence of LDL, an essential step for atherosclerosis progression. ARF proteins belong to the superfamily of Ras-like GTPases. Six ARF isoforms are found in mammals. ARF1 and ARF6 are the best-characterized isoforms and act as key regulators of vesicle formation and trafficking, membrane lipid transformation, actin remodeling, cell migration and invasion (11). Strategies targeting the inhibition of ARF1 activation in VSMCs prevent phenotypic switching by regulating the expression of contractile markers (12). Although the role of ARF6 in this context remains elusive, this ARF isoform acts as a molecular switch to promote angiotensin II type 1 receptor (AT_1_R) internalization and extracellular signal-regulated kinase ½ (Erk1/2) activation. Furthermore, ARF6 increases VSMC proliferation, migration and reactive oxygen species (ROS) production, which are often enhanced in disease state (8, 9). Our latest findings demonstrate that ARF6 also plays a key role in regulating angiotensin II (Ang II) and platelet-derived growth factor (PDGF)-dependent invasion of human aortic smooth muscle cells (HASMCs) (10).

Although cultured VSMCs can help define key proteins and signaling axes important for phenotypic switching, whether the findings from these cells can be replicated in pre-clinical models remains a concern. Elucidation of the role of ARF6 *in vivo* has been particularly challenging due to embryonic mice lethality (13). To confirm our findings in a pathological model, we have thus generated a novel conditional knockout (CKO) mouse that lacked *Arf6* in smooth muscle cells (SMCs) using a Cre-LoxP system, in an atherosclerosis-prone background (ApoE^−/−^) to examine whether ARF6 controls the function of VSMCs and development of the plaque *in vivo*.

Altogether, our findings demonstrate that the depletion of ARF6 hinders the phenotypic transition of VSMCs induced by lipid stimulation. Therefore, the expression and activation of this GTPase constitute a prerequisite for the development of atherosclerosis in a pre-clinical model. Future therapeutic approaches will need to consider VSMC plasticity, as current drugs used to prevent atherosclerosis are primarily limited to lowering cholesterol levels and blood pressure. Identification of the molecular mechanisms regulating phenotypic switching will provide excellent opportunities for the discovery of new pharmacological targets and the design of novel, targeted drugs.

## Results

### ARF6 regulates LDL uptake in HASMCs by modulating the expression of scavenger receptors and micropinocytosis

Migration of VSMCs is a complex process influenced by various factors. As illustrated in Figure 1A, the addition of OxLDL for 4h significantly enhanced the migration of HASMC. Notably, depletion of ARF6 inhibited both basal and OxLDL-promoted effects. Prolonged exposure of VSMCs to OxLDL can upregulate the expression of scavenger receptors to facilitate lipid uptake and foam cell formation. To investigate this, we stimulated control and ARF6-depleted HASMCs with OxLDL for 72h and assessed protein levels of LOX-1, MSR1, LDLR, CD36 as well as TLR4 by Western blotting (Fig 1B). First, we observed that OxLDL increased LOX-1 expression, an effect that was completely abolished by the knockdown of ARF6. Although the expression of MSR1 was not significantly enhanced by lipid stimulation, ARF6 depletion effectively prevented expression of this scavenger receptor in our cell model. Conversely, levels of LDLR, CD36, and TLR4 remained unchanged following the depletion of this GTPase.

**Figure 1.**
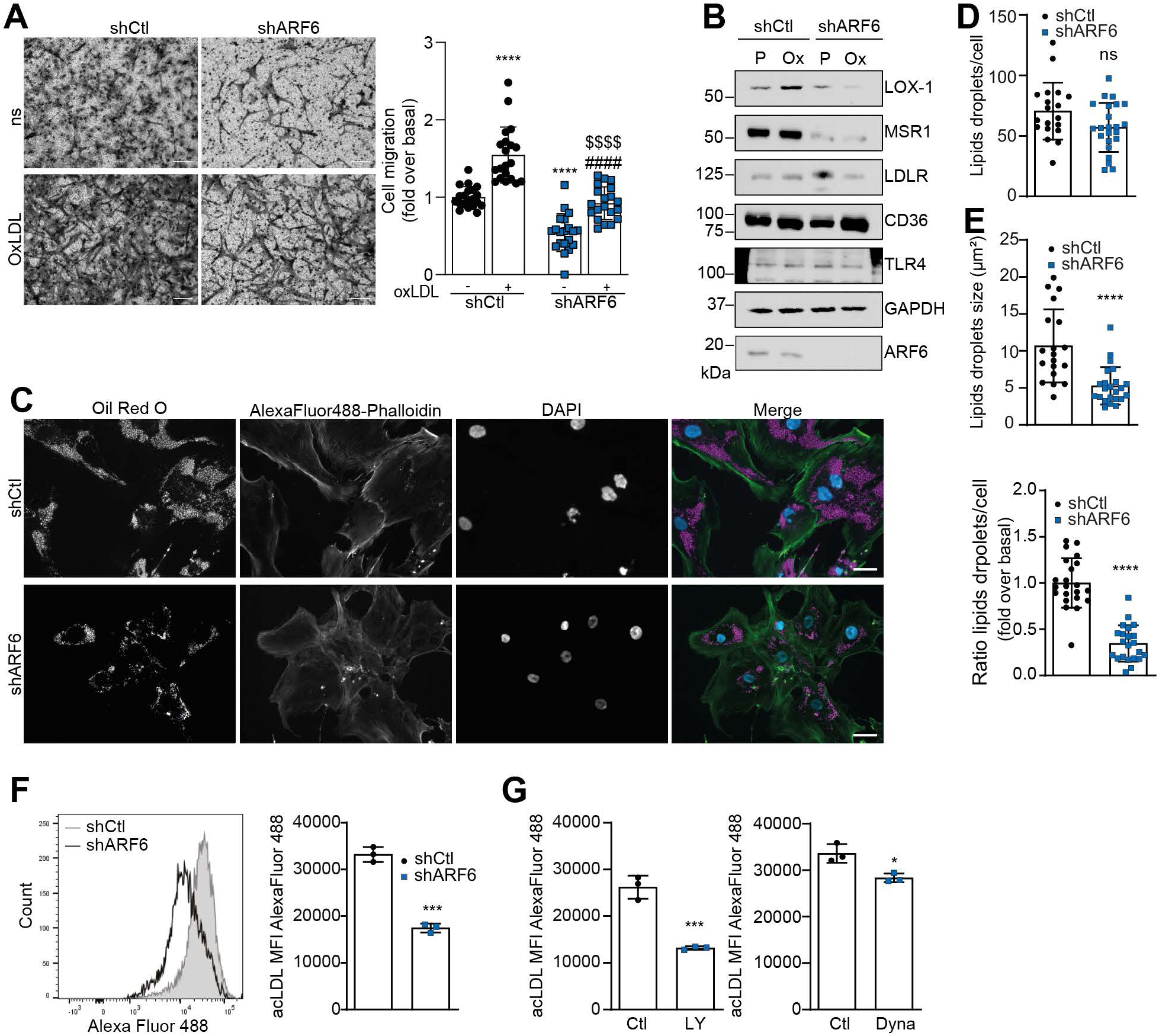
ARF6 modulates phenotypic switching in HASMCs. **A**) Cells were infected with untargeting shRNA Ctl (shCtl) or ARF6 shRNA (shARF6) lentivirus. Infected HASMCs were seeded into Collagen type 1-coated Boyden chambers and stimulated with 80 µg/mL OxLDL for 4 h. Images are from the lower part of the membrane and are representative of five images taken per condition. Quantifications are the mean ± SD of 20 images from four independent experiments. ****P < 0.0001 are values compared to the basal condition, ####P < 0.0001 are values compared to shCtl OxLDL conditions, and $$$$ P < 0.0001 are values compared to shARF6 ns (Two-Way ANOVA with Tukey’s multiple comparison test) (bar = 100 µm). **B**) Infected HASMC were stimulated with OxLDL (80 µg/ml) or PBS and lysed after 72 h. Expression levels of indicated proteins were assessed by Western blotting using specific antibodies. Images are representative of four independent experiments. (P: PBS, Ox: OxLDL). **C, D, E**) Cells were infected as in **A** and were incubated with high-fat mice serum for 72 h. After fixation, cells were stained with Oil Red O and actin stained with Alexa Fluor 488-phalloidin. Images are representative of three independent experiments. Quantifications are the mean ± SD of 22 images representative of 100 cells per condition from three independent experiments. ****P<0.0001, ns not significant (Unpaired Student’s *t*-test). Bar scale: 50 µm. **F)** Infected cells were incubated with Alexa Fluor 488-labelled AcLDL (1 µg/ml) for 18 h and then analyzed by flow cytometry. Quantifications are mean ± SD of mean fluorescence intensity (MFI) of triplicates from one experiment, representative of three independent experiments. ***P < 0.001 (Unpaired Student’s *t*-test). **G**) Cells were incubated with DMSO (Ctl), LY294002 50 µM (LY), or Dynasore 100 µM (Dyna) 1 h before adding Alexa Fluor 488-labelled AcLDL (1 µg/ml) for 18 h. Cells were analyzed by flow cytometry. Quantifications are the mean ± SD of triplicates from one experiment, representative of three independent experiments. ***P < 0.001, *P<0.05 (Unpaired Student’s *t*-test).

Furthermore, cells were treated for 72h with serum derived from high-fat diet-fed mice. As illustrated in Figure 1C, Oil Red O staining revealed lipid droplet accumulation within HASMCs. Depletion of ARF6 significantly reduced lipid accumulation. Additional analysis demonstrated that while the droplet number was comparable in control and ARF6-depleted conditions, their size was markedly decreased when the GTPase was knocked down (average diameter: 10 µm in control vs. 5 µm in ARF6 shRNA cells; Fig. 1D, E). Next, we examined HASMC uptake of acetylated (AcLDL), an artificially modified LDL form, labelled with Alexa Fluor 488. Flow cytometry analysis showed a significant reduction in AcLDL accumulation in ARF6-depleted cells when compared to control (Fig. 1F). To further investigate the mechanism by which this ARF isoform controls intracellular lipid accumulation, we pretreated cells with either LY294002 (50µM), a micropinocytosis inhibitor targeting PI3K (14), or Dynasore (100 µM), which inhibits dynamin-dependent internalization processes such as clathrin-coated vesicles and caveolae-mediated pathways. As shown in Figure 1G, both inhibitors significantly reduced AcLDL uptake. However, the micropinocytosis pathway appeared to be the predominant mechanism utilized for LDL internalization.

Altogether, these findings suggest that expression of ARF6 is essential for the ability of VSMCs to accumulate lipids and switch to a foam-cell-like phenotype primarily through regulation of scavenger receptor expression and micropinocytosis.

### Cytokine secretion, inflammatory marker expression, and collagen production in OxLDL-stimulated HASMC depend on the activation of ARF6

Given that VSMC-derived foam cells and plaque-infiltrated immune cells release numerous pro-inflammatory cytokines which contribute to local inflammation in atherosclerotic lesions (2), we aimed to determine whether ARF6 could act to regulate IL-6 secretion. It is widely recognized that when foam cells are stimulated by pro-inflammatory mediators such as IL-1ß and TNFα, they increase IL-6 production, as this plays a pivotal role in amplifying the inflammatory cycle in atherosclerosis. As illustrated in Figure 2A, the addition of OxLDL increased IL-6 secretion from HASMCs, an effect significantly attenuated by the depletion of ARF6. qPCR analysis confirmed that ARF6 knockdown reduced *IL6* mRNA levels by over 50%. Furthermore, stimulation of the cells with the pro-inflammatory cytokines IL-1β and TNFα also increased IL-6 secretion. These stimuli also require the expression of ARF6 (Fig. 2B), as all cytokines, in addition to OxLDL, potently activate this ARF isoform, as evidenced by increased GTP loading (Fig. 2C).

**Figure 2.**
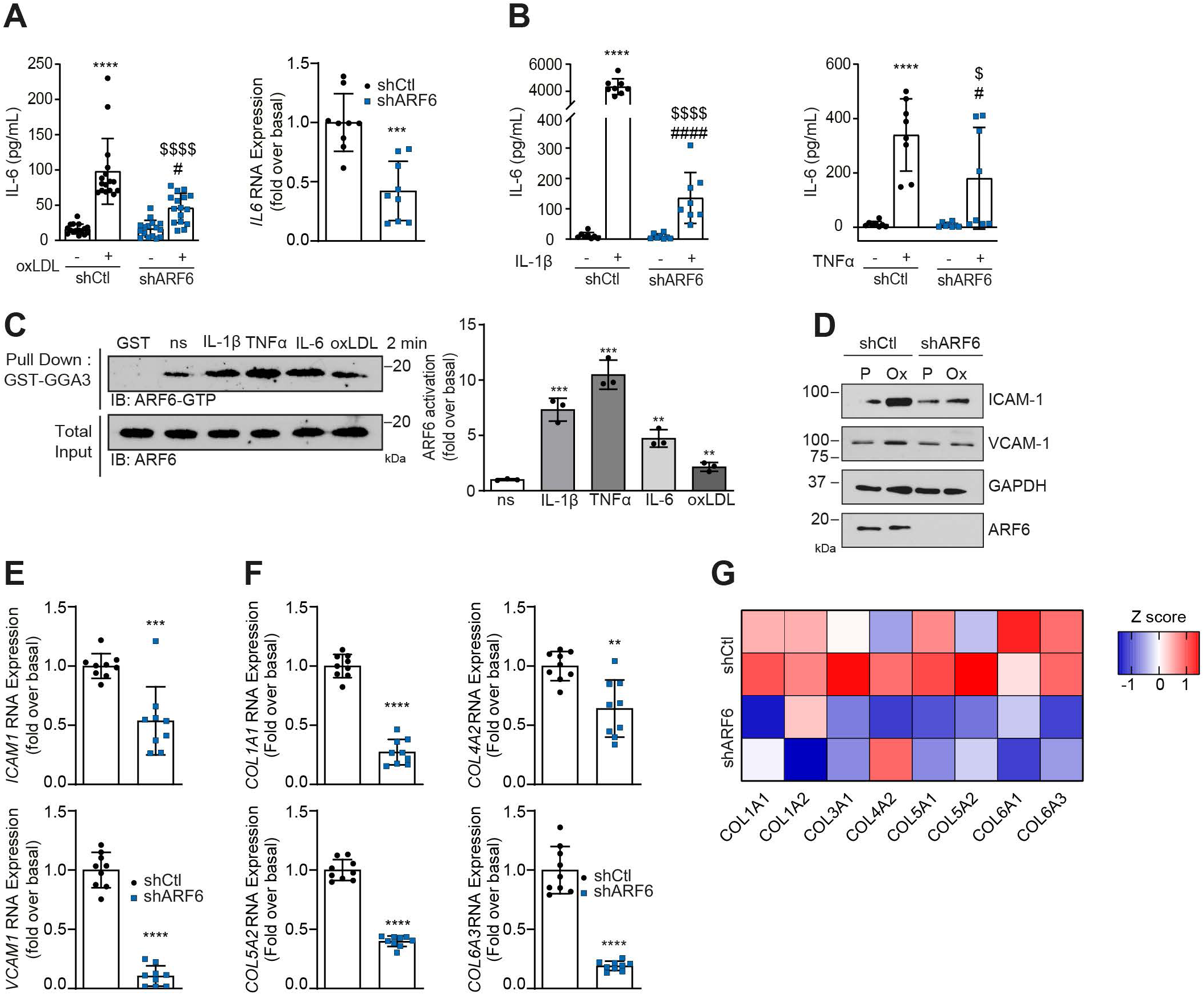
ARF6 modulates IL-6 and collagen secretion. **A)** Infected cells were stimulated with OxLDL (80 µg/ml) for 24 h. Supernatants were collected and processed to perform an IL-6 ELISA according to the manufacturer’s instructions. Quantifications are the mean ± SD of technical duplicates from eight independent experiments. ****P<0.0001 compared to the basal condition, $$$$P<0.0001 compared to Ctl stimulated, #P<0.05 compared to shARF6 ns (Two-way ANOVA with Tukey’s multiple comparison test). Total RNA of HASMCs was extracted with the Qiagen RNeasy kit according to the manufacturer’s instructions. Quantifications are the mean ± SD of triplicates from three independent experiments. ***P<0.001 (Unpaired Student’s *t*-test). **B**) Infected HASMCs were incubated with IL-1β (10 ng/ml) or TNFα (25 ng/ml) for 24 h. Supernatants were collected and processed to perform an IL-6 ELISA according to the manufacturer’s instructions. Quantifications are the mean ± SD of duplicates from four independent experiments. ****P<0.0001 compared to the basal condition, $$$$P<0.0001; $P<0.05 compared to Ctl stimulated; ####P<0.000, # P<0.05 compared to shARF6 ns (Two-way ANOVA with Tukey’s multiple comparison test). **C**) Cells were serum-starved for 16 h and stimulated with IL-1β (10 ng/ml), TNFα (25 ng/ml), IL-6 (10 ng/ml), or OxLDL (80 µg/ml) for 2 min. Cells were lysed and ARF6 activity was assessed in a GTPase activation assay with GST-GGA3. ARF6-GTP levels were evaluated by Western blotting using specific antibodies against ARF6. Quantifications are the mean ± SD of three independent experiments. ***P < 0.001, **P<0.01 are values compared to the basal condition (One-Way ANOVA with Dunnett’s multiple comparisons test). **D**) Infected HASMCs were stimulated with OxLDL (80 µg/ml) or PBS and lysed after 72 h. Expression levels were assessed by Western blotting using specific antibodies. Images are representative of three independent experiments. (P: PBS, Ox: OxLDL). **E, F**) Infected cells were stimulated with OxLDL (80 µg/ml) for 72 h. Total RNA of HASMCs was extracted with the Qiagen RNeasy kit according to the manufacturer’s instructions. Quantifications are the mean ± SD of triplicates from three independent experiments. ****P<0.0001,***P < 0.001, **P<0.01 are values compared to the basal condition (Unpaired Student’s *t*-test). **G**) Infected cells were starved for 16 h. Supernatants were collected and concentrated for mass spectrometry analysis. Heat map of Z scores of spectral counts of collagen fiber proteins. n=2.

We next examined the expression of inflammation markers when cells were exposed to OxLDL. Intracellular Adhesion Molecule-1 (ICAM-1) and Vascular Cell Adhesion Molecule-1 (VCAM-1) are upregulated during the development of atherosclerosis. These are usually considered markers of endothelial cell dysfunction. However, when VSMCs undergo phenotype switching phenotype and are exposed to a pro-inflammatory environment, they overexpress ICAM-1 and VCAM-1, enabling VSMCs to interact with monocytes and lymphocytes, thus contributing to inflammation and lesion progression (15). Figure 2D shows that OxLDL caused an upregulation of ICAM-1 and VCAM-1 protein levels in HASMCs. However, knockdown of ARF6 was effective in reducing the expression of both markers. This effect was also observed at the transcriptional level as mRNA expression of *ICAM1* and *VCAM1* was reduced in ARF6-depleted HASMCs, further suggesting that ARF6 contributes to the pro-inflammatory phenotype of VSMCs (Fig. 2E).

Because synthetic VSMCs secrete large amounts of collagen, leading to vascular wall remodeling, we next examined whether ARF6 could modulate this key feature. The absence of ARF6 reduced the mRNA expression of several types of collagen fibers (*COL1A1, COL4A2, COL5A2, COL6A3*) when cells were exposed to OxLDL for 72 hours (Fig. 2F). To confirm these findings, we measured protein levels of secreted collagen fibers under basal and ARF6-depleted conditions using mass spectrometry. Figure 2G shows that HASMCs secrete types III, IV, V and VI collagens, and that the absence of ARF6 significantly reduces the secretion of all these forms, thus highlighting ARF6’s role in modulating extracellular matrix components during vascular remodeling.

### OxLDL activate ARF6 through TLR4 signaling, promoting IL-6 secretion

Considering ARF6’s role as a key regulator of multiple signaling pathways, we investigated whether this GTPase could modulate OxLDL-induced signaling events. As illustrated in Figure 3A, OxLDL stimulation resulted in the phosphorylation of AKT, p65NF-κB and STAT3, all of which were inhibited in ARF6-depleted cells (Fig. 3A). We also examined the activation of the MAP kinase pathways (p38, ERK1/2 and JNK). In our conditions, phosphorylation of all MAP kinase pathways was induced by OxLDL. However, depletion of the GTPase significantly reduced the activation of only p38 (Fig. 3B). As OxLDL signaling through NF-κB and p38 results in increased transcription of KLF4 to promote lipid uptake, foam cell formation, and inflammatory reprogramming (16), we investigated whether depletion of ARF6 could affect the expression of this transcription factor. Figure 3C shows that depletion of the GTPase significantly inhibited *KLF4* mRNA levels in OxLDL-treated HASMCs.

**Figure 3.**
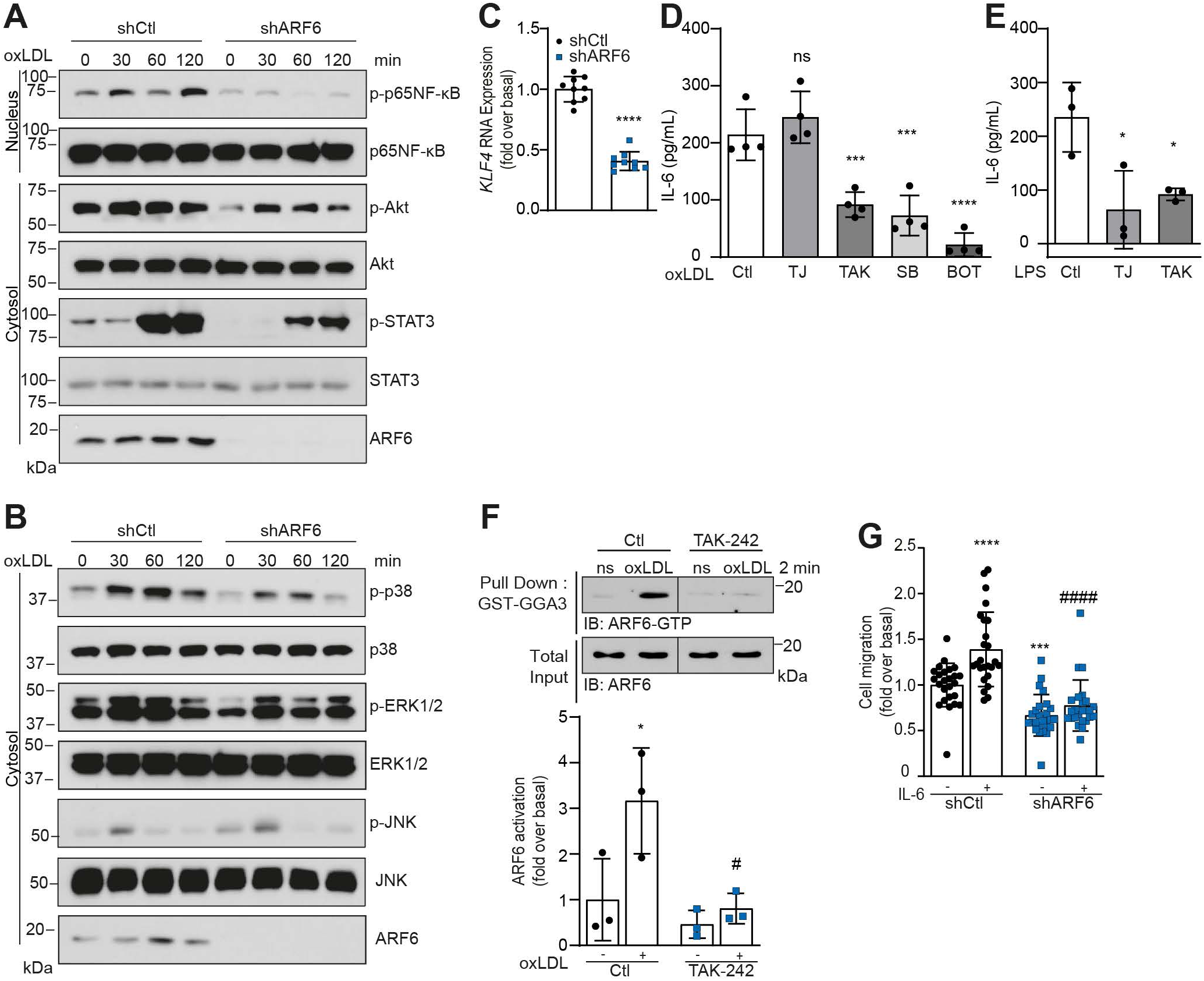
ARF6 controls IL-6 secretion through TLR4. **A, B**) Infected cells were stimulated for 0, 30, 60, and 120 min with OxLDL (80 µg/ml). Cytosolic or nucleus expression levels of indicated proteins were assessed by Western blotting using specific antibodies. Images are representative of three independent experiments. **C**) Infected cells were stimulated with OxLDL (80 µg/ml) for 72 h. Total RNA of HASMCs was extracted with the Qiagen RNeasy kit according to the manufacturer’s instructions. Quantifications are the mean ± SD of triplicates from three independent experiments. ****P<0.0001 (Unpaired Student’s *t*-test). **D, E**) Cells were pretreated with DMSO (Ctl), TJ-m2010-5 (TJ) (40 µM), TAK-242 (TAK) (30 µM), SB203580 (SB) (10 µM) or BOT-4 (BOT) (10 µM) for 1 h. Cells were then stimulated with OxLDL (80 µg/mL) (**D**) or LPS (100 ng/mL) (**E**) for 24 h. Supernatants were collected and processed to perform an IL-6 ELISA according to the manufacturer’s instructions. Quantifications are the mean ± SD from four (**D**) or three (**E**) independent experiments. ****P<0.0001,***P < 0.001, *P<0.05 are values compared to the basal condition, ns not significant (One-way ANOVA with Dunnett’s multiple comparisons test). **F**) Cells were pretreated for 1 h with DMSO (Ctl) or TAK-242 (30 µM) before stimulation with OxLDL (80µg/mL) for 2 min. ARF6 activity was assessed in a GTPase activation assay with GST-GGA3. ARF6-GTP levels were evaluated by Western blotting using specific antibodies against ARF6. Quantifications are the mean ± SD of three independent experiments. *P<0.05 compared to the basal condition, #P<0.05 compared to Ctl OxLDL (Two-Way ANOVA with Tukey’s multiple comparison test). **G**) Infected cells were seeded into collagen type I-coated Boyden chambers for 1 h. Afterwards, cells were left untreated or stimulated with IL-6 (10 ng/ml) for 4 h. Quantifications are the mean ± SD of 25 images per condition from five independent experiments. ***P < 0.001, ****P < 0.0001 are values compared to the basal condition, ####P < 0.0001 are values compared to shCtl IL-6 conditions (Two-Way ANOVA with Tukey’s multiple comparison test).

Next, we further investigated the mechanism by which ARF6 regulates IL-6 secretion. Indeed, we examined the involvement of the Toll-Like Receptor 4 (TLR4), which is known to mediate inflammation by regulating the phenotypic switching of VSMCs to macrophage-like and foam cells (17). Pretreatment of OxLDL-stimulated HASMCs with the TLR4 inhibitor TAK242 decreased IL-6 secretion (47%) (Fig. 3D). TLR4 can activate two pathways, the MyD88-dependent and MyD88-independent signaling cascades. Here, pretreatment of the cells with the MyD88 inhibitor TJ-m2010-5 did not affect OxLDL-stimulated IL-6 secretion. To confirm the efficacy of this inhibitor, cells were stimulated with LPS which can activate both MyD88-dependent and independent pathways (18). In these conditions, both TAK-242 and TJ-m2010-5 decreased IL-6 secretion (Fig. 3E). To better define the molecular mechanism and confirm the involvement of p38 and NF-κB, we pretreated OxLDL-stimulated HASMCs with SB203580 and BOT-4, respectively. Figure 3D illustrates that inhibiting either pathway led to a decrease in IL-6 secretion. To confirm the role of ARF6 in regulating this signaling axis, we examined whether TLR4 stimulation could indeed lead to ARF6 activation. As illustrated in Figures 2C and 3F, OxLDL stimulated the cells to promote GTP-loading of this ARF isoform. Pretreatment of the cells with the TLR4 inhibitor, TAK-242, completely blocked ARF6 activation caused by OxLDL (Fig 3F).

Lastly, as IL-6 is known for its ability to act in an autocrine fashion in VSMCs to promote migration (19), we investigated whether ARF6 could be the GTPase that mediates IL-6-promoted migration. Using the Boyden chamber assay, we demonstrated that the addition of IL-6 enhances HASMC migration by 39% under the control condition. However, depletion of ARF6 inhibited both the basal migration capability of the cells and the IL-6-mediated response (Fig. 3G). These results suggest that ARF6 expressed in VSMCs regulates different aspects of inflammation following TLR4 activation and that the GTPase is also a key player in IL-6 autocrine signaling.

### The knockout of ARF6 in SMCs decreases the formation of atherosclerotic plaques in ApoE^−/−^ mice

Building on the insights gained from our findings in cultured cells, we aimed to investigate the inherent function of ARF6 in VSMC function within a pre-clinical atherosclerotic model. To do so, we generated a novel conditional knockout mouse model that lacked *Arf6* in smooth muscle cells by crossing *Acta2-Cre-ERT2/ApoE^−/−^* transgenic mice with *Arf6*^flox/flox^ mice (*Acta2-Cre-ERT2^+/−^/ApoE^−/−^/Arf6^f/f^*) (Fig. 4A). We have previously shown that there is no Cre recombinase leak when *Acta2-Cre-ERT2/ApoE^−/−^*mice were crossed with Cre stain reporter (lacZ)(20). To induce ARF6 KO, 8-week-old male mice were intraperitoneally injected with tamoxifen (TAM) for five consecutive days. As demonstrated in Figure 4B, DNA quantification revealed decreased *Arf6* gene expression in ARF6 KO mice compared to control mice (those injected with peanut oil). The knockout of ARF6 was specific to smooth muscle cell-rich tissues, such as the aorta, and was not found in the kidney or liver (Fig. 4C), as demonstrated by mRNA quantification. To further characterize knockout efficiency, we examined protein levels of the GTPase in tissue extracts. As shown in Figure 4D, ARF6 protein expression was decreased in both the aorta and the intestine, two tissues rich in smooth muscle cells, but not in the liver. Using immunohistochemistry, we also confirmed the knockout of ARF6 in tissues by demonstrating that ARF6 expression was reduced in smooth muscle actin (SMA)-rich cells in the aorta (Fig. 4E). Furthermore, we investigated the long-term effect of TAM exposure on atherosclerotic lesion formation. 8-week-old male mice were injected with vehicle or TAM for five consecutive days. Animals were kept on a standard diet for 6 months post-injection. Lesions in the aorta were observed in both groups, with no significant difference (Fig. S1A and B). Besides, we tested the effect of TAM on lipid internalization and foam cell phenotypic switching in HASMCs. Using the TAM metabolite, (Z)-4-Hydroxytamoxifen (4OH-TAM), we observed no effect of TAM on AcLDL internalization (Fig. S1C) or scavenger receptor expression (Fig S1D).

**Figure 4.**
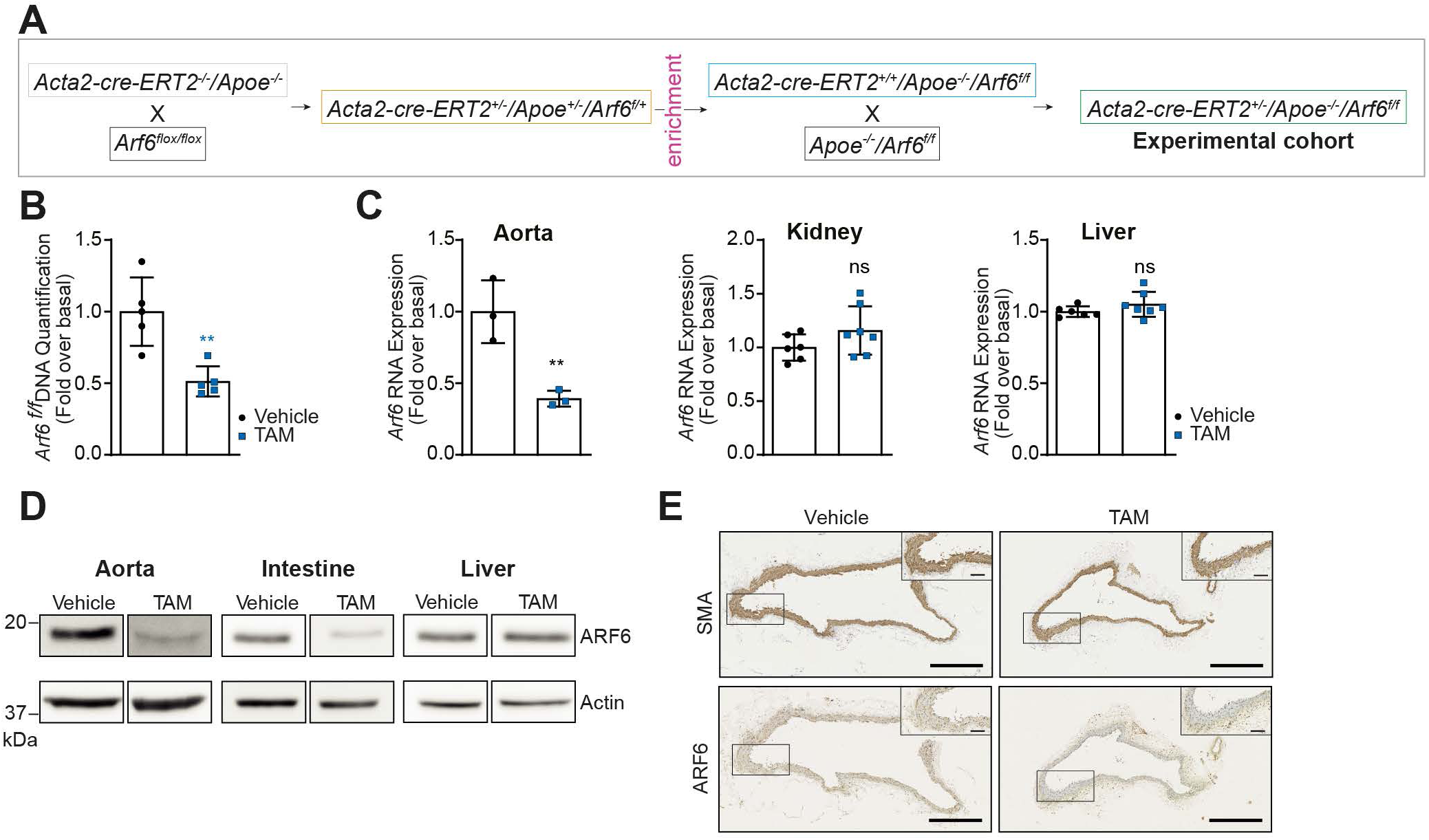
Characterization of *Acta2-Cre-ERT2^+/−^/ApoE^−/−^/Arf6^flox/flox^ mice*. **A**) Schematics of the breeding protocol to generate the desired genotype for the study. **B**) DNA was extracted from the whole aorta with the Qiagen AllPrep DNA/RNA kit according to the manufacturer’s instructions. A qPCR targeting *Arf6^flox^* was performed. Quantifications are the mean ± SD of five animals per group (Vehicle or TAM (Tamoxifen)). **P<0.01 (Unpaired Student’s *t*-test). **C**) Total RNA was extracted from the aorta, kidney or liver with the AllPrep DNA/RNA kit, both according to the manufacturer’s instructions. Quantifications are the mean ± SD of six or seven animals per group (three animals per group for the aorta). **P<0.01, ns not significant (Unpaired Student’s *t*-test). **D**) Total proteins were extracted from whole organs with RIPA buffer. ARF6 expression was assessed by Western blotting using specific antibodies. The blots are representatives of 3 animals per group. **E**) Representative aorta cross-sections stained for immunohistochemistry using specific ARF6 or SMA antibodies. Bar scale: 500 µm or 100 µm for close-ups.

To initiate the formation of atherosclerotic lesions, all animals were fed a high-fat diet (HFD) for 10 weeks after TAM or vehicle injections (Fig. 5A). Both the control animals and ARF6 KO mice exhibited comparable weight gain throughout the HFD treatment (Fig. 5B). The diet induces hypercholesterolemia, as total cholesterol levels were found to be elevated by about 3-fold when compared with standard regular chow diet (21). Notably, we observed no differences in the lipid profiles of control and ARF6 KO mice (Table S1). To determine whether the deletion of the *Arf6* gene specifically in SMCs impacts atherosclerosis development, we first examined the plaque in ARF6 KO mice. Analysis of Oil Red O-stained *en-face*-opened aorta of control and ARF6 KO mice fed a HFD for 10 weeks showed no overall difference in lesion development (Fig. 5C). However, when specifically examining the aortic arch, the section of the aorta where the blood flow is most disturbed, we observed a significant decrease in the percentage of atherosclerotic plaque area of mice treated with TAM (6,7% ± 0,6; n=12) compared to vehicle-treated mice (8,6% ± 0,6; n=14) (Fig. 5D). Heart sections further showed that the size of the lesions found in the aortic sinus was diminished in ARF6 KO animals (12,2% ± 1,4; n=14) compared to vehicle-treated animals (16,4% ± 0,8; n=9) (Fig. 5E). Furthermore, Hematoxylin & Eosin (H&E) coloration revealed that the intimal thickness of the atheroma was reduced in the aortic sinus of ARF6 KO mice (Fig. 5F).

**Figure 5.**
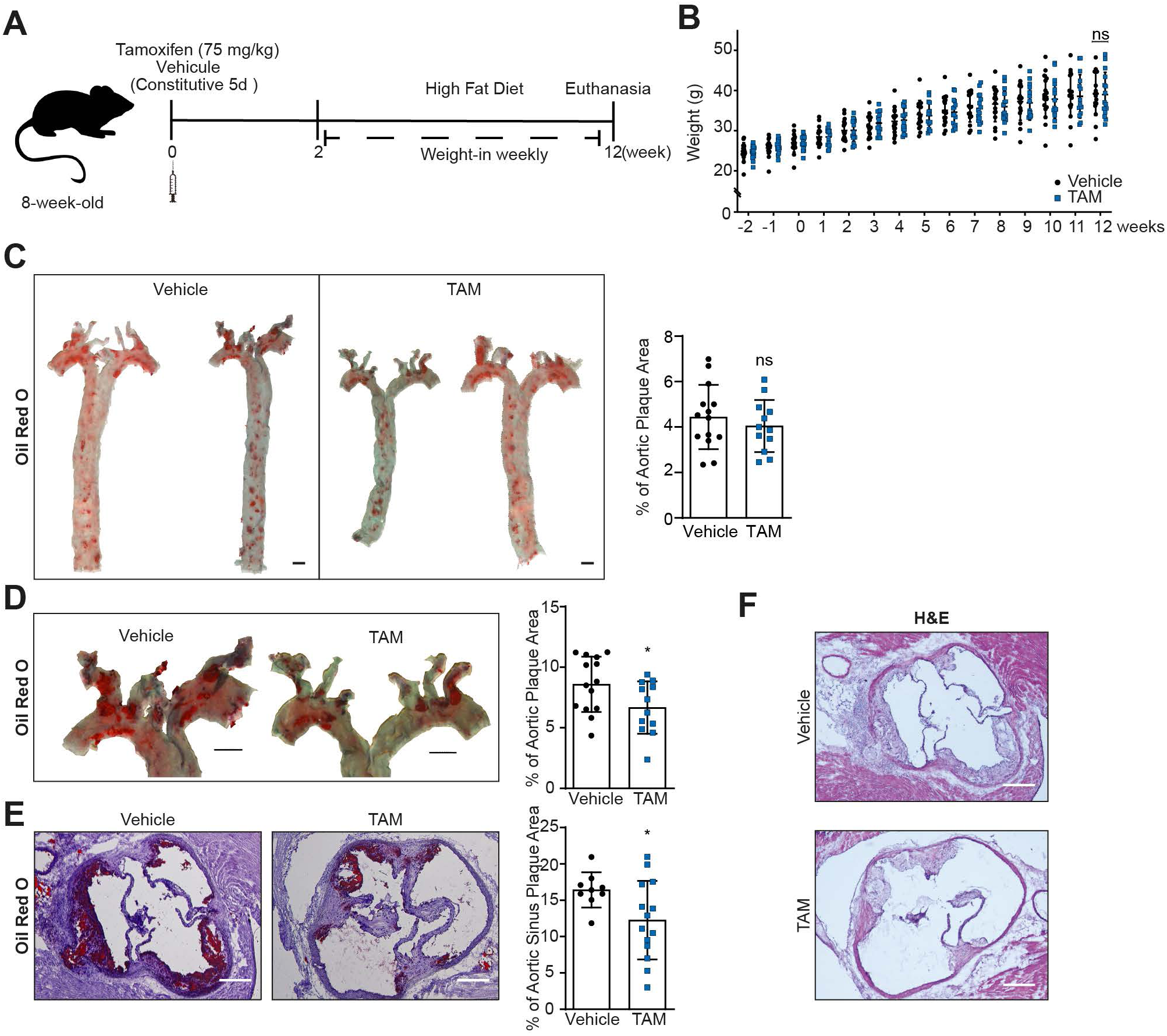
Knockout of ARF6 in smooth muscle cells reduces atherosclerotic lesions. **A**) Schematics of the experimental design. After five daily injections of Vehicle or Tamoxifen (75 mg/kg), mice were fed with a high-fat diet for ten weeks. **B**) Mice were weighed weekly for 14 weeks. Quantifications are the mean ± SD of 15 or 17 animals per group. ns not significant (Two-way ANOVA with Sidak’s multiple comparisons test). **C**) Representative *en-face* aorta (**C**) or aortic arch (**D**) stained with Oil Red O (ORO) lipid stain to visualize atherosclerotic lesions. Quantifications are the mean ± SD of 15 or 17 animals per group. *P<0.05, ns not significant (Unpaired Student’s *t*-test) Bar scale: 1 mm. **E, F**) Image of frozen cross-sections of the aortic sinus stained with Oil Red O (**E**) or Hematoxylin & Eosin stains (H&E) (**F**). Bar scale: 50 µm. Quantifications are the mean ± SD of 15 or 17 animals per group. *P<0.05 (Unpaired Student’s *t*-test).

### The composition of atherosclerotic lesions is altered when ARF6 is knocked out in VSMCs

During the progression of atherosclerosis, dedifferentiated VSMCs migrate throughout the plaque, contributing to the development of the fibrous cap. Using H&E staining, we observed that the fibrous cap of the atheroma was reduced in brachiocephalic arteries of ARF6 KO mice when compared to controls (Fig. 6A). To investigate the importance of VSMCs in this process, we utilized α-SMA as a marker of contractile smooth muscle cells. As shown in Figure 6B, α-SMA staining was present inside and around the plaque. When ARF6 was knocked out, SMC staining was still visible around the plaque, but the area of positive staining was significantly reduced. Next, since VSMCs can secrete components of the extracellular matrix, we investigated the composition of the lesions in ARF6 KO mice. Here, Masson Trichrome staining was performed to visualize connective tissues, particularly the presence of collagen. In vehicle-treated mice, histological analysis showed large collagen-rich plaques in the aortic sinus, while in ARF6 KO mice, plaques were still visible but showed reduced staining intensity (Fig. 6C). Transversal sections of *en-face*-opened brachiocephalic artery showed a similar reduction of collagen staining (Fig. 6D). To confirm these observations, we performed Sirius Red staining experiments. As illustrated in Figure 6E, a significant decrease in collagen staining was observed in atherosclerotic lesions of ARF6 KO mice. Altogether, our findings suggest that ARF6 expression in VSMCs regulates the composition of atherosclerotic lesions, particularly the presence of extracellular matrix components, such as collagen.

**Figure 6.**
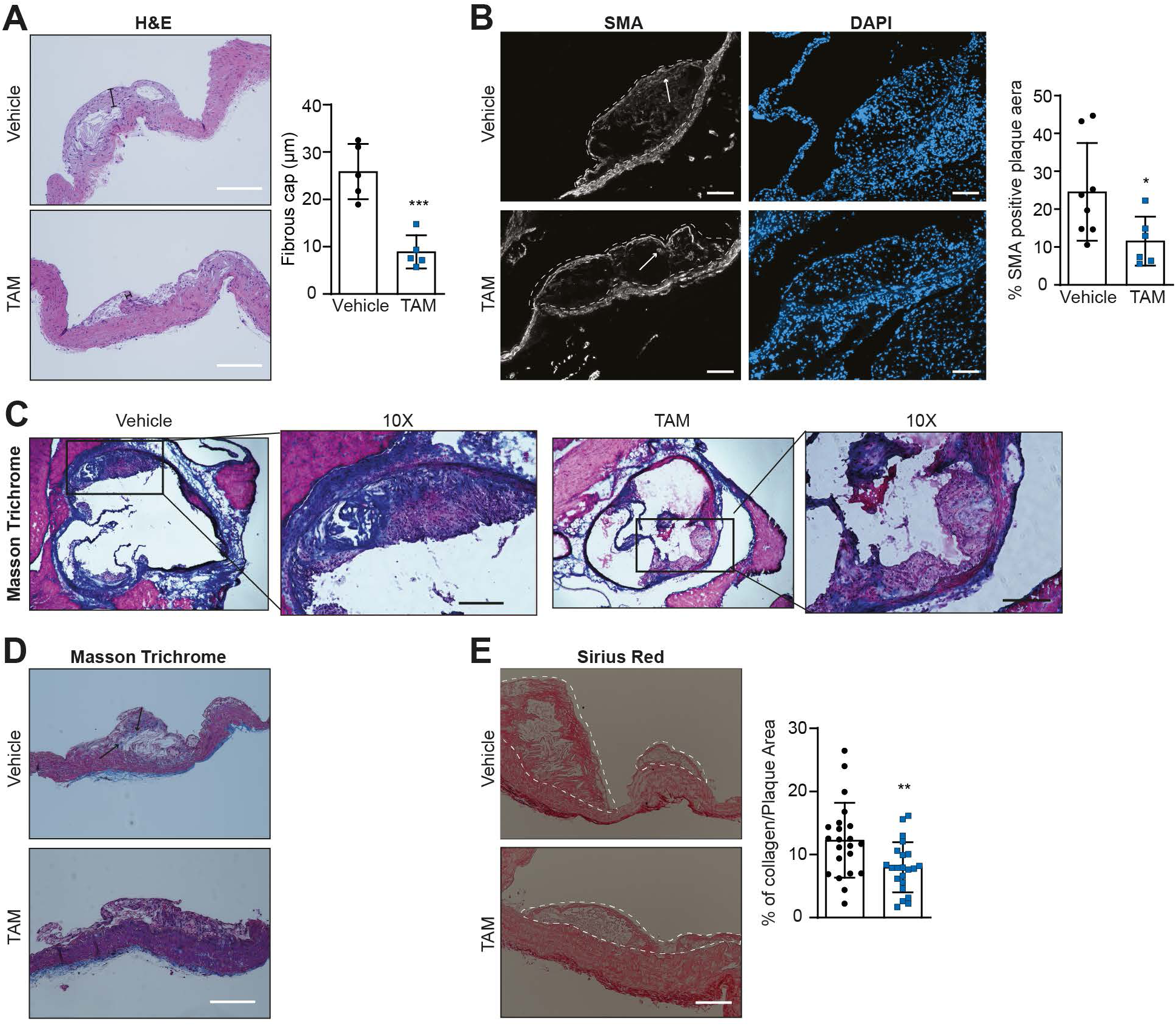
ARF6 knock-out modifies lesion composition. **A**) Images of aortic arch cross-sections stained with H&E staining. Bar identifies the fibrous cap. Three to five atheroma were analyzed per animal. Quantifications are the mean ± SD of five animals per group. ***P<0.001 (Unpaired Student’s *t*-test). Bar scale: 100 µm. **B**) Immunofluorescence of frozen aortic sinus stained with anti-SMA and then with Alexa Fluor 568-labelled secondary antibody. Nuclei were stained with DAPI. Images are representative of three non-constitutive sections per animal, n=5 per group. Quantifications are the mean ± SD 6 or 8 lesions. *P<0.05 (Unpaired Student’s *t*-test). Arrows represent SMA-positive regions. Dotted lines encircle the lesions. Bar scale: 100 µm. **C**) Aortic sinus cross-sections marked with Masson’s trichrome staining. Images are representative of 5 animals per condition. Bar scale: 100 µm. **D**) Images of aortic arch cross-sections marked with Masson’s trichrome staining. n=5 animals per group. Arrows identify the collagen regions. Bar scale: 100 µm. **E)** Aortic arch cross-sections were stained with PicroSirius Red stain, and collagen content was quantified in the plaque aera. Three to five non-consecutive sections of plaque were analyzed. Quantifications are the mean ± SD of 3 to 5 lesions from four animals per group. **P<0.01 (Unpaired Student’s *t*-test). Dotted lines encircle the lesions. Bar scale: 200 µm.

### Ablation of ARF6 expression in aortic SMCs reduces local inflammation in lesions

In addition to a decreased amount of extracellular matrix components, we also observed a reduction in the quantity of foam cells present in the aortic arch lesions of ARF6 KO mice (Fig 7A). Given that VSMC-derived foam cells and plaque-infiltrated immune cells release numerous pro-inflammatory cytokines which contribute to local inflammation in atherosclerotic lesions, we next isolated native murine VSMCs from the abdominal aorta and analyzed various inflammatory markers. As illustrated in Figure 7B, ARF6 knockout decreased mRNA expression of cell adhesion molecules (*Icam1, Vcam1*), cytokine and cytokine receptors (*Il-6, Ccl2, TNF, Il1r1*) in our *in vivo* model. Furthermore, *Rela*, an important gene involved in inflammation pathways, was also reduced in ARF6 KO VSMCs compared to controls. However, this reduction in the expression of inflammation markers was not due to an effect on the infiltration of immune cells to the atherosclerotic lesion. Indeed, no difference in the expression of CD45 or CD68 was observed between the two groups (Fig. 7C, D). We also analyzed the frequencies of the main immune cell populations by flow cytometry in the peripheral blood (Fig. S2A), spleen (Fig. S2B), and bone marrow (Fig. S2C) of control and ARF6 KO mice. In the peripheral blood, there was no modulation of different subpopulations of immune cells. However, in the immune organs, the frequency of total immune cells and most immune cell types was decreased in ARF6 KO mice compared to control animals.

**Figure 7.**
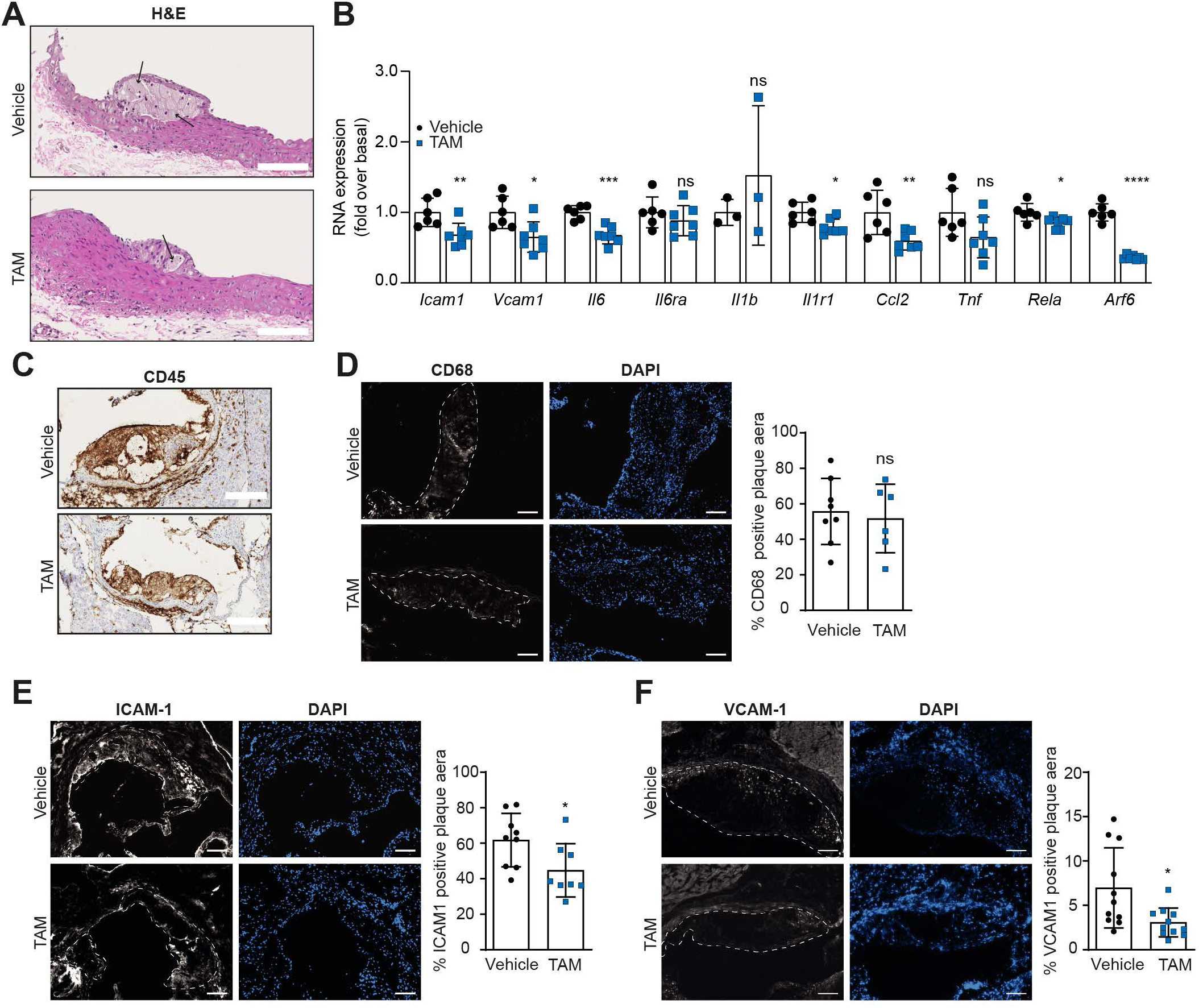
ARF6 knockout affects local inflammation in the plaque. **A**) Images of aortic arch cross-sections stained with H&E. Arrows identify the foam cells. n=5 animals per group. Bar scale: 100 µm. **B**) Total RNA was extracted from mouse aortic VSMCs with the Qiagen RNeasy kit according to the manufacturer’s instructions. Quantifications are the mean ± SD of six or seven animals per group (3 animals per group for *Il1*β). ****P<0.0001, ***P<0.001, **P<0.01,*P<0.05, ns not significant (Unpaired Student’s *t*-test). **C**) Representative aortic sinus cross-sections stained for immunohistochemistry with anti-CD45. n=5 animals per group. Bar scale: 250 µm. **D**) Immunofluorescence of frozen aortic sinus stained with anti-CD68 and Alexa Fluor 568. Nuclei were stained with DAPI. Images are representative of three non-constitutive sections per animal, n=5 per group. Quantifications are the mean ± SD of 6 or 8 lesions per group. ns not significant (Unpaired Student’s *t*-test) **E, F**) Immunofluorescence of frozen aortic sinus stained with anti-ICAM-1(**E**) or anti-VCAM-1(**F**) and Alexa Fluor 568. Nuclei were stained with DAPI. Images are representative of three non-constitutive sections per animal, n=5 per group. Quantifications are the mean ± SD of 8 to 11 lesions per group. *P<0.05 (Unpaired Student’s *t*-test). Dotted lines encircle the lesions. Bar scale: 100 µm.

To confirm our observations, we further examined the protein expression of inflammation markers (ICAM-1, VCAM-1) within the aortic sinus tissues. As illustrated in Figure 7E, ICAM-1 is widely expressed in heart sections of mice treated with vehicle. Conversely, in heart sections of mice treated with TAM, there was a notable decrease in the area percentage of plaques containing this marker. In comparison to ICAM-1, VCAM-1 displayed a generally lower presence within the atherosclerosis plaques and a reduction was also observed in ARF6 KO mice (Fig. 7F).

Altogether, these results suggest that ARF6 expression in SMCs accentuates local inflammation in the aorta during lipid-induced atherosclerosis. These *in vivo* studies thus confirm our *in vitro* findings in isolated human smooth muscle cells.

## Discussion

Understanding the underlying mechanisms responsible for the initiation and progression of atherosclerosis is of paramount significance to identify effective strategies to control the advancement of this disease. Our study highlights the key role of the small GTP-binding protein ARF6 in controlling VSMC transition into a transdifferentiated phenotype, which facilitates LDL uptake, ECM secretion, inflammatory mediator production, and migration. This was demonstrated using cultured cells of human vascular origin, as well as a genetically engineered atherosclerosis mouse model. Our approach has focused on the consequences of knocking down or knocking out ARF6, thereby inhibiting the expression of this protein and its ability to become activated in response to stimuli such as high levels of circulating lipids, a well-documented risk factor for vascular diseases, and cytokines, the key mediators of inflammation.

In recent years, we have established that ARF proteins serve as pivotal regulators of cellular migration and proliferation of, namely, rat aortic VSMCs (8–10). Although cell models are widely used and can be predictive of an *in vivo* role, whether our observations can be translated into human pathophysiology remains a concern. Animal models, with their limitations, represent the best alternative to studying disease development in the intact microenvironment. Although ARF proteins have been shown to regulate processes ranging from vesicle trafficking to complex cellular responses in cells, the physiological functions of this family of GTPases in whole animals was unknown until recently. The first global knockout of ARF6 in mice resulted in nearly complete embryonic lethality, beginning at mid-gestation and persisting through birth. Affected embryos displayed multiple abnormalities, including head edema, with occasional instances of hemorrhage and anemia during the mid-to-late stages of gestation. Additionally, a marked reduction in liver size was evident, attributed to impaired hepatic cord formation (13). Conversely, generation of conditional knockout animals confirmed numerous key roles for ARF6 *in vivo*. For example, deletion in neurons showed impaired oligodendrocyte myelination (22). Using an endothelial cell-specific conditional knockout of ARF6 in mice, it was reported that ARF6 played a crucial role in tumor angiogenesis and growth (23). Because *in situ* hybridization of ARF6 mRNA in E13.5 mouse embryos revealed transcripts in vascular smooth muscle cells of the dorsal aorta (23) and because we have shown a key role of this GTPase in regulating important VSMC functions, we generated a novel ARF6 conditional knockout mouse model in which the deletion of the ARF6 gene can be temporally and specifically achieved in SMCs.

Here, using this new inducible atherosclerosis mouse model, *Acta2-Cre-ERT2^+/−^/ApoE^−/−^/Arf6^f/f^*, we show for the first time that ARF6 controls specific atherosclerosis processes. Namely, Oil Red O (ORO) staining of aortic arches revealed that ARF6 deletion decreases the tissue area exhibiting atherosclerotic plaques. The fact that heart sinus sections further showed an important effect of ARF6 deletion may suggest that the expression of this GTPase in VSMCs affects both cardiac and vascular functions during the development of the pathology. Upon closer examination of the atherosclerotic plaque structure, we observed that the accumulation of collagen is less significant in lesions of TAM-treated mice. This confirmed our findings in isolated cultured HASMCs, where we demonstrated that ARF6 depletion inhibits the ability of these cells to express and secrete various collagen isoforms (I, III, IV, V, and VI). Namely, collagen I is highly secreted by VSMCs during disease development (24) and contributes to maintaining VSMCs in a synthetic phenotype, thereby keeping the cells in a pathological state. Extracellular matrix proteins, in general, have been associated with plaque calcification and rupture (25). After 10 weeks of HFD treatment, we did not observe calcification of atherosclerotic lesions in our model as this process occurs later, nor a difference in necrosis (26).

The intriguing ability of VSMCs to switch phenotypes and acquire properties relevant to different pathological states is complex. VSMC can switch phenotype to become macrophage-like cells and express scavenger receptors, such as LOX-1, and acquire phagocytosis functions (27, 28), but can also develop characteristics reminiscent of foam cells when in the presence of OxLDL, and enhance inflammatory pathways. This leads to an increase in the production of proinflammatory cytokines and the expression of adhesion cell molecules and scavenger receptors, which then fuels the chronic inflammatory environment (29–31). In HASMCs, we demonstrate that the intracellular accumulation of modified lipids, particularly OxLDL, results in a foam cell-like phenotype. Other groups have suggested that ARF6 controls cholesterol uptake (32), further supporting our findings. To better define the molecular mechanisms regulated by the GTPase during this process, we demonstrated that ARF6 depletion alters the expression of scavenger receptors associated with lipid uptake. Thus, we suggest that the decrease in intracellular lipid accumulation is caused in part by the inhibition of LOX-1 and MSR1 expression, as well as a faulty dynamin-dependent internalization process, such as micropinocytosis. LOX-1 has been reported to mediate OxLDL endocytosis via a clathrin-independent internalization pathway (33). We have previously suggested that ARF6 and the ARF GAP GIT1 can regulate clathrin-dependent and independent G protein-coupled receptor internalization (34–36). Another possibility that may impair intracellular lipid accumulation is the inability of ARF6-depleted cells to store, metabolize, or promote efflux through specialized transporters.

Our experiments, aimed at identifying the molecular mechanisms regulated by ARF6, have revealed a key role of this GTPase in the TLR4 signaling axis, a well-established pathway in the progression of atherosclerosis. By upregulating the expression of cytokines and adhesion molecules involved in cell recruitment and proliferation, TLR4 is a key contributor to inflammation and foam cell accumulation (37, 38). Although the TLR4 MyD88-dependent pathway results in the activation of NFκB and the production of inflammatory cytokines, its inhibition using TJ-m2010-5 did not affect OxLDL-stimulated IL-6 secretion. This suggests that, in HASMCs, ARF6 may act through the MyD88-independent pathway, which uses TRIF/TRAM to activate MAPK and NFκB supporting a previous study showing that this ARF isoform was essential for TLR4/TRIF/TRAM signaling in HEK293 cells (39).

As supported by our previous findings in cell cultures, our results suggest that deletion of ARF6 in VSMCs reduces the ability of these cells to migrate and proliferate in an *in vivo* pathological setting. The repartition of SMA-positive cells in atherosclerosis lesions present in the aortic sinus or brachiocephalic artery showed that VSMCs, in TAM-treated mice, remained at the edge of the plaque. Also, the reduction in VSMC-derived foam cells is associated with a decrease in the expression of inflammation markers, such as ICAM-1 and pro-inflammatory cytokines such as TNFα, CCL2 and IL-6. Considering the importance of inflammation in the pathogenicity of atherosclerosis, pharmacological strategies which aim at reducing this response have shown potential for delaying atherosclerosis progression (40). The reduction of inflammation in the atherosclerotic lesions alters the immune cell populations. Indeed, in atherogenic conditions, circulating monocytes are chemoattracted to the intima by the local expression of pro-inflammatory molecules and the overexpression of adhesion molecules in the vicinity of the vessel (41). The infiltrating monocytes will differentiate into macrophages, which will then become foam cells and thus contribute to the development of the plaque (42). Since ARF6 is also expressed in myeloid cells, the inactivation of the *Arf6* gene in these cells may also contribute to reducing atherosclerotic lesions. In addition to being expressed in smooth muscle cells, fibroblasts, and myofibroblasts, the *Acta2* gene is also present in mice circulating neutrophils and monocytes (43, 44). Thus, the effect we observed on plaque formation in the *Acta2-Cre-ERT2^+/−^ /ApoE^−/−^/Arf6^f/f^*mouse model may be partly mediated by immune cells. Furthermore, the reduced inflammatory environment in ARF6 KO mice will inhibit the recruitment of pro-atherosclerotic immune cells in circulation, which may be reflected in the hematopoiesis in the bone marrow (45).

Atherosclerosis is a chronic inflammatory disease where there is continuous crosstalk between lipid metabolism and activation of inflammatory pathways. As such, it is well established that the inflammatory cytokine IL-6 is highly expressed within atherosclerosis lesions and exerts pro-atherosclerotic effects (46). In our *in vitro* model, we noted a decrease in the expression and secretion of IL-6 when ARF6 was knocked down. Given that this cytokine serves as an autocrine regulator of VSMC function and that decreased IL-6 secretion is associated with diminished VSMC mobility, this could explain the reduced VSMC presence within atherosclerotic lesions when ARF6 is absent.

In conclusion, our study has significantly advanced our understanding of the *in vivo* role of ARF6 in the progression of atherosclerosis in a preclinical model. Our findings establish that, in response to OxLDL stimulation, ARF6 facilitates the trans-differentiation of VSMCs into foam cells by upregulating scavenger receptor expression, and the ability of OxLDL to signal through the TLR4/MAPK/NF-κB pathway which leads to inflammation. This enhanced understanding of the pivotal regulators governing VSMC phenotypic transition holds promise for the identification of new therapeutic targets for the treatment of the vascular wall diseases marked by dysfunctional smooth muscle cell behavior. Our work makes a substantial contribution to the broader understanding of the roles played by ARF proteins in cells of the cardiovascular system.

## Experimental procedures

### HASMC Cell culture

Human Aortic Smooth Muscle Cells (HASMC) were obtained from ScienCell Research Laboratories (Carlsbad, CA), cultured in smooth muscle cell medium (ScienCell Research Laboratories) according to the manufacturer’s instructions, and maintained in a humidified incubator with 5% CO_2_ at 37°C. All experiments were performed between passages 4 and 8. HASMC were infected in the presence of polybrene (8 µg/ml) with untargeted (control) or ARF6 shRNA lentiviruses. Stable clones were selected using puromycin (2 µg/ml) and cells were used for experiments 72 hours (h) post-infection.

### Lentiviral constructs and virus production

MISSION ARF6 shRNA plasmids were purchased from Sigma Aldrich (ARF6: TRCN0000381410, TRCN0000286788, TRCN0000380270, TRCN0000294067 and TRCN0000294069; non-targeting sequence SHC016). The ARF6 TRCN0000380270 (AGCTGCACCGCATTATCAATG) target sequence was chosen as it showed the highest level of knockdown in HVSMC cells (10). Lentivirus containing the shRNA were generated using 293T cells transfected with pLKO-ARF6 or non-targeting shRNA constructs and the psPax.2 and pMD2.G packaging plasmids (Addgene, Watertown, MA).

### qPCR

DNA/RNA from mice aorta was extracted using the Qiagen AllPrep DNA/RNA kit (Qiagen, Qc). qPCR was performed with primers specific for *Arf6^flox/flox^* flox sites to verify the cleavage of *Arf6*. Absence of qPCR signal confirmed the cleavage of *Arf6*. RNA from cultured HAMSCs or other mouse tissues was extracted using the Qiagen RNAeasy kit (Qiagen, Qc). All qPCR experiments were conducted at the IRIC genomic platform (Université de Montréal).

### Antibodies

Anti-ICAM-1, anti-CD36 and anti-OLR1 were from Proteintech (Chicago, IL). Anti-ARF6 and anti-LDLR were from Abcam Biochemicals (Cambridge, MA). HRP-conjugated secondary antibodies and anti-VCAM-1 were from R&D Systems (Minneapolis, MN). All other antibodies were obtained from Cell Signaling Technology (Danvers, MA).

### ARF6 activation assay

HASMCs were serum starved for 16 h and stimulated for 2 minutes (min) with IL-1β (10 ng/ml), TNFα (25 ng/ml), IL-6 (10 ng/ml), or OxLDL (80 µg/ml). Cells were harvested in lysis buffer E (pH 7.4, 50 mM Tris–HCl, 1% Nonidet P-40, 137 mM NaCl, 10% glycerol, 5 mM MgCl_2_, 20 mM NaF, 1 mM NaPPi, 1 mM Na_3_VO_4_, and protease inhibitors). Samples were spun for 10 min at 10,000 × g at 4 °C. GST-GGA3 fusion proteins coupled to glutathione-Sepharose 4B beads were added to each tube, and samples were rotated at 4°C for 1 h (47). Beads were washed, and proteins were eluted into 20 μl of SDS-sample buffer containing 5% β-mercaptoethanol by heating at 65°C for 15 min. Proteins were resolved on 14% SDS-PAGE. Western blot analysis was done using a specific anti-ARF6 antibody.

### Cell lysis and Western blotting

For total soluble proteins, tissues or cells were lysed in buffer E as previously described. For the signaling pathways, cells were first lysed in a hypotonic buffer (20 mM Tris-HCl (pH 7.4), 10 mM NaCl, 3 mM MgCl_2_, 0.5% NP-40) for the cytosolic fraction, and then the nuclear pellet was resuspended in cell extraction buffer (10 mM Tris (pH 7.4), 100 mM NaCl, 1 mM EGTA, 1 mM EDTA, 1 mM NaF, 20 mM Na_4_P_2_O_7_, 2 mM Na_3_VO_4_, 1% Triton X-100, 10% glycerol, 0.1% SDS, 0.5% deoxycholate). Soluble proteins were run on SDS-PAGE gels and transferred onto nitrocellulose membranes. The membranes were blotted for relevant proteins using specific primary antibodies as described for each experiment. HRP-conjugated secondary antibodies were used and the chemiluminescence reaction was triggered using the Amersham ECL Prime Western detection reagent. Membranes were exposed to autoradiography films, which were scanned using an HP scanner. For Figures 2C and 3F, bands were quantified using ImageJ (NIH, Bethesda, MD, USA).For Figure 4D, membranes were digitalized using GE LAS 4000 mini.

### Lipid uptake

Post-infection, HASMCs were serum-starved for 16 h and then stimulated with vehicle (PBS) or high fat diet mouse serum for 72 h. Cells were fixed with a 10% formalin solution for 10 min at room temperature (RT), and then stained with ORO in an isopropanol solution for 30 min. After washing, samples were labelled with Alexa Fluor 488 phalloidin. Coverslips were mounted onto slides with Aqua-Mount and analyzed using a Zeiss inverted microscope Axio Observer. For fluorescence activated cell sorting (FACS) experiments, infected cells were serum-starved for 16 h and then stimulated with Alexa Fluor 488 acetylated LDL (AcLDL) (ThermoFisher) for 18 h. Cells were trypsinized, washed twice, and analyzed with a BD FACSCanto II flow cytometer. For the analysis of foam cell markers or signaling pathways, cells were stimulated with vehicle or irradiated-OxLDL (80 µg/ml) (Sigma Aldrich, Oakville, ON) for indicated times. The preparation of irradiated-OxLDL was performed as described in (48). Cells were lysed for total RNA or total proteins.

### Migration Assay

Transfected cells were serum starved and seeded into Boyden chambers (24-well inserts with 8-μm pore collagen I-coated membranes). Afterwards, cells were left untreated (non-stimulated; ns) or stimulated with IL-6 (10 ng/ml) for 4 h. Cells were subsequently fixed using paraformaldehyde (4%) for 20 min and incubated with crystal violet (0.1% in 20% MeOH, overnight). Membranes were washed three times in distilled H_2_O, and cells were removed from the upper chamber by scraping, leaving those that migrated through the membrane in the lower chamber. Pictures of five different fields per well were taken, and cell migration was quantified using ImageJ. The average number of migrating cells was determined for each condition.

### IL-6 ELISA

The concentration of IL-6 released into the medium was determined using an ELISA kit for human IL-6 (ThermoFisher Scientific inc). Briefly, post-infected HASMCs were incubated with IL-1β (10 ng/ml), TNFα (25 ng/ml), or OxLDL (80 µg/ml) for 24 h. When inhibitors were used, cells were pretreated 1 hour before stimulation with OxLDL. Supernatants were collected and processed to perform an IL-6 ELISA according to the manufacturer’s instructions.

### Mass spectroscopy

Culture media of serum-starved controlled-or ARF6-depleted cells were concentrated using Amicon™ Ultra-15 Centrifugal Filter Units, 30 kDa MWCO (Sigma Aldrich, Oakville, ON). One hundred micrograms of proteins was resuspended in laemmli buffer and subjected to electrophoresis. Bands were destained in 50% MeOH (Sigma-Aldrich). Each band was shrunk in 50% acetonitrile (ACN), reconstituted in 50 mM ammonium bicarbonate with 10 mM TCEP [Tris(2-carboxyethyl) phosphine hydrochloride; Thermo Fisher Scientific], and vortexed for 1 h at 37°C. Chloroacetamide (Sigma-Aldrich) was added for alkylation to a final concentration of 55 mM. Samples were vortexed for another hour at 37°C. One microgram of trypsin was added, and digestion was performed for 8 h at 37°C. Peptide extraction was conducted with 90% ACN. The extracted peptide samples were dried down and solubilized in 5% ACN-0.2% formic acid (FA). The samples were loaded on a C4 guard column (Optimize Technologies) connected directly to the switching valve. They were separated on a home-made reversed-phase column (150-μm i.d. by 200 mm packed with Jupiter C18, Phenomenex) with a 56-min gradient from 10 to 30% ACN-0.2% FA and a 600-nl/min flow rate on an Easy-nLC 1200 connected to an Orbitrap HF Biopharma (Thermo Fisher Scientific, San Jose, CA). Each full MS spectrum acquired at a resolution of 120,000 was followed by tandem-MS (MS-MS) spectrum acquisition for a maximum of 3 s. on the most abundant multiply charged precursor ions. Tandem-MS experiments were performed using higher energy collisional dissociation (HCD) at a collision energy of 34%. The data were processed using PEAKS X Pro (Bioinformatics Solutions, Waterloo, ON) and a Uniprot human database. Mass tolerances on precursor and fragment ions were 10 ppm and 0.3 Da, respectively. Variable selected posttranslational modifications were carbamidomethyl (C), oxidation (M), deamidation (NQ), acetyl (N-ter) and phosphorylation (STY). Data were viewed using Scaffold 5.3.1 (Proteome Software inc.) and extracted as Excel file (Supporting Information S3). Heat map was generated with Z scores from normalized data using GraphPad Prism software version 10.0 (GraphPad PRISM, San Diego, CA).

### Animal protocol

The local institutional animal ethics committee (CDEA) approved all animal studies according to the Canadian Council of Animal Protection. *Acta2-CreERT2/ApoE^−/−^*transgenic mice were first generated using *Acta2-CreERT2 and ApoE^−/−^*mice. These were crossed with *Arf6*^flox/flox^ mice (22) to generate parental lines that were further crossed to produce experimental cohorts (*Acta2-Cre-ERT2^+/−^/ApoE^−/−^/Arf6^flox/flox^*). All animals were numbered and kept in a temperature-controlled room on a 12-hour light/12-hour dark schedule. Mice were randomly divided into two groups. The first group was intraperitoneally injected with 100 µl of peanut oil (vehicle), while the other group was injected with Tamoxifen (75 mg/kg/day) in 100 µl of vehicle for 5 consecutive days, alternating sides each day. After one week of rest, both groups were fed a high-fat Western diet (Envigo, TD.96121) ad libitum. After 10 weeks, mice were anesthetized with isoflurane, and cardiac puncture followed by pneumothorax was used for euthanasia. Different organs were excised and used for protein or RNA extraction, immunofluorescence or FACS analysis. Serum was also collected using BD Microtainer tubes according to the manufacturer’s instruction (BD Biosciences, Mississauga, ON) and total cholesterol was measured according to the manufacturer’s instruction (Abcam Biochemicals, Cambridge, MA).

### En-face analysis of Atherosclerosis in Mice

After euthanasia, a visual examination of the organs showed no other pathology than fat accumulation around the aorta. Microdissection was then done as described in (49) under stereomicroscope. The entire aortas were degreased using forceps and microdissection scissors. Depending on the experiments, the aortas were dissected from the mice and either stored in formalin (10%) solution for 24 h for ORO staining and immunohistochemistry (IHC), or snap-frozen for entire protein and DNA/RNA extraction. The abdominal sections were used for VSMCs isolation. For *en-face* experiments, the aorta was cut from the animal at the iliac arteries and at 1-2 mm from the heart. The entire aorta was then placed in phosphate buffer saline (PBS) and any residual fat was removed using forceps. The thoracic aorta was then pinned onto paraffin paper coated with black electric tape for easy visualization using 1 mm thick pins, and placed in a Petri dish as described in (49). The aorta was then cut open and *en-face* pinned in a PBS-containing Petri dish. Tissues were fixed, washed three time in cold PBS and placed in ORO (0.7% in propylene glycol) at 37°C for 16 h. Aortas were washed in five consecutive solutions (H_2_O, methanol, PBS) and counterstained with Light Green (0.05%) solution. The pictures were obtained using a Nikon D800 camera with macro lens.

### VSMC isolation

Briefly, dissected and degreased 5mm-sections of abdominal aortas were bathed for 15 min in a digestion solution (collagenase II 0.15%, dispase II 0.2%, 5 mM CaCl_2_ in HBSS) at 37°C. The tunica was then removed under a stereomicroscope. Aortas were further incubated for 3 h at 37°C in the digestion solution and were passed through a 70 µm cell strainer. Cells were snap-frozen for RNA extraction.

### Immunohistochemistry and Immunofluorescence

IHC was performed on 4 µm paraffin sections, and anti-ARF6 antibody (dilution 1/350) was applied to every section for 2 h at RT. Sections were then incubated with a specific secondary biotinylated antibody. Streptavidin horseradish peroxidase and 3,3-diaminobenzidine were used according to the manufacturer’s instructions (DABmap detection kit, Ventana Medical Systems). For H&E staining of heart transversal sections 10 µm-thick sections were fixed for 10 min with ice-cold 10% formalin, washed with PBS, and stained with Mayers hematoxylin for 30 seconds

(sec). Sections were sequentially rinsed in PBS and tap water, and dipped in ethanol (EtOH) 80%. Next, sections were counterstained with eosin for 30 sec, dehydrated with EtOH and cleared in xylene. Sections were mounted on microscope slides using Leica mounting media. For H&E staining of *en-face* opened aorta transversal sections, Oil Red O-stained aortas were washed in PBS and processed for paraffin embedding. 4 µm-thick paraffin sections were deparaffinized, rehydrated and stained with H&E as described above.

For Masson trichome staining, heart sections were processed following Trichrome Stain kit (Sigma, HT15). Slides were washed in tap water and stained in working Weigert’s Iron Hematoxylin Solution for 5 min. After washes, sections were stained in Biebrich Scarlet-Acid Fucshin for 5 min. After rinses, sections were placed in a Phosphotungstic/Phosphomolybdic Acid solution for 5 min. Finally, sections were placed successively in Aniline Blue Solution for 5 min and Acetic Acid for 2 min before dehydration in EtOH and clearing in xylene. ARF6, CD45 and Sirius red colorations were performed at the IRIC Histology Core Facility (Université de Montréal).

For immunofluorescence labelling, heart sections were fixed in 4% paraformaldehyde (PFA) at RT for 15 min. After PBS washing, sections were incubated with glycine to quench free aldehyde groups. Sections were then blocked in a solution containing 5% normal serum and 0.3% Triton for 60 min. Specific antibodies diluted in 1% bovine serum albumin (BSA) and 0.3% Triton/PBS were incubated overnight at 4°C as described in figure legends. After several washes, the sections were incubated in the dark for 1 h in the presence of an anti-mouse or anti-rabbit Alexa Fluor 568 antibody (Invitrogen). Sections were then mounted using a solution of Aqua-mount (Fisher Scientific, Ottawa, Canada) and observed using a Zeiss inverted microscope Axio Observer.

## Statistical analysis

Statistical analyses were conducted with GraphPad Prism software version 10.0 (GraphPad PRISM, San Diego, CA) using the Student’s *t*-test, a one-way ANOVA, or a two-way ANOVA analysis. A P value <0.05 was considered statistically significant.

## Supporting information

This article contains supporting information.

## Acknowledgements

We would like to thank Professor Pierre Chambon from Université de Strasbourg for sharing the Acta2-CreERT2 mice. We also thank Dr. Louis Gaboury from Université de Montréal and Dr. Franck Ah-Pine from CHU de La Réunion site SUD for their expert advice on immunohistochemistry. We are grateful for the expert help of Dr Marc Saba-El Leil and Isabelle Caron from the In Vivo Biology IRIC platform. Proteomics analyses were performed by the Center for Advanced Proteomics Analyses, a Node of the Canadian Genomic Innovation Network that is supported by the Canadian Government through Genome Canada

## Abbreviations

ARF: ADP-ribosylation factors
SMC: smooth muscle cells
CVD: cardiovascular diseases
VSMC: vascular smooth muscle cells
HASMC: Human aortic smooth muscle cells
ECM: extracellular matrix
HFD: high fat diet
TAM: tamoxifen
IHC: immunohistochemistry
LDLR: low density lipoprotein receptor
VLDLR: very low density lipoprotein receptor
angiotensin II: Ang II
AT_1_R: angiotensin II type 1 receptor
PDGF: platelet-derived growth factor
Erk1/2: extracellular signal-regulated kinase 1/2
PAK: p21-activated kinase
ROS: reactive oxygen species
MMP: matrix metalloprotease
CKO: conditional knockout
KO: knockout
PBS: phosphate buffer saline
H&E: hematoxylin and eosin
ORO: oil red O
EtOH: ethanol
h: hour
min: minute
sec: second
PFA: paraformaldehyde
RT: room temperature
BSA: bovine serum albumin
IL: interleukin
TNF_α_: tumor necrosis factor alpha
OxLDL: oxidized LDL
FACS: fluorescence activated cell sorting
AcLDL: acetylated LDL

## References

1. Chen, R., McVey, D. G., Shen, D., Huang, X., and Ye, S. (2023) Phenotypic Switching of Vascular Smooth Muscle Cells in Atherosclerosis Journal of the American Heart Association 12, e031121 10.1161/JAHA.123.031121

2. Grootaert, M. O. J., and Bennett, M. R. (2021) Vascular smooth muscle cells in atherosclerosis:Time for a reassessment Cardiovasc Res 10.1093/cvr/cvab046

3. Wang, Y., Dubland, J. A., Allahverdian, S., Asonye, E., Sahin, B., Jaw, J. E. et al. (2019) Smooth Muscle Cells Contribute the Majority of Foam Cells in ApoE (Apolipoprotein E)-Deficient Mouse Atherosclerosis Arteriosclerosis, Thrombosis, and Vascular Biology 39, 876–887 doi:10.1161/ATVBAHA.119.312434

4. Doran, A. C., Meller, N., and McNamara, C. A. (2008) Role of Smooth Muscle Cells in the Initiation and Early Progression of Atherosclerosis Arteriosclerosis, Thrombosis, and Vascular Biology 28, 812–819 doi:10.1161/ATVBAHA.107.159327

5. Chattopadhyay, A., Kwartler, C. S., Kaw, K., Li, Y., Kaw, A., Chen, J. et al. (2021) Cholesterol-Induced Phenotypic Modulation of Smooth Muscle Cells to Macrophage/Fibroblast-like Cells Is Driven by an Unfolded Protein Response Arterioscler Thromb Vasc Biol 41, 302–316 10.1161/atvbaha.120.315164

6. Clarke, M. C. H., Talib, S., Figg, N. L., and Bennett, M. R. (2010) Vascular Smooth Muscle Cell Apoptosis Induces Interleukin-1–Directed Inflammation Circulation Research 106, 363–372 10.1161/CIRCRESAHA.109.208389

7. Martinet, W., Schrijvers, D. M., and De Meyer, G. R. (2011) Necrotic cell death in atherosclerosis Basic Res Cardiol 106, 749–760 10.1007/s00395-011-0192-x

8. Charles, R., Namkung, Y., Cotton, M., Laporte, S. A., and Claing, A. (2016) β-Arrestin-mediated Angiotensin II Signaling Controls the Activation of ARF6 Protein and Endocytosis in Migration of Vascular Smooth Muscle Cells J Biol Chem 291, 3967–3981 10.1074/jbc.M115.684357

9. Bourmoum, M., Charles, R., and Claing, A. (2016) The GTPase ARF6 Controls ROS Production to Mediate Angiotensin II-Induced Vascular Smooth Muscle Cell Proliferation PLOS ONE 11, e0148097 10.1371/journal.pone.0148097

10. Fiola-Masson, E., Artigalas, J., Campbell, S., and Claing, A. (2022) Activation of the GTPase ARF6 regulates invasion of human vascular smooth muscle cells by stimulating MMP14 activity Sci Rep 12, 9532 10.1038/s41598-022-13574-7

11. Jackson, C. L., and Bouvet, S. (2014) Arfs at a glance J Cell Sci 127, 4103–4109 10.1242/jcs.144899

12. Charles, R., Bourmoum, M., and Claing, A. (2018) ARF GTPases control phenotypic switching of vascular smooth muscle cells through the regulation of actin function and actin dependent gene expression Cell Signal 46, 64–75 10.1016/j.cellsig.2018.02.012

13. Suzuki, T., Kanai, Y., Hara, T., Sasaki, J., Sasaki, T., Kohara, M. et al. (2006) Crucial role of the small GTPase ARF6 in hepatic cord formation during liver development Mol Cell Biol 26, 6149–6156 26/16/6149 [pii] 10.1128/MCB.00298-06

14. Kruth, H. S., Jones, N. L., Huang, W., Zhao, B., Ishii, I., Chang, J. et al. (2005) Macropinocytosis is the endocytic pathway that mediates macrophage foam cell formation with native low density lipoprotein J Biol Chem 280, 2352–2360 10.1074/jbc.M407167200

15. Sorokin, V., Vickneson, K., Kofidis, T., Woo, C. C., Lin, X. Y., Foo, R., et al. (2020) Role of Vascular Smooth Muscle Cell Plasticity and Interactions in Vessel Wall Inflammation Frontiers in Immunology 11, 10.3389/fimmu.2020.599415

16. Yap, C., Mieremet, A., de Vries, C. J. M., Micha, D., and de Waard, V. (2021) Six Shades of Vascular Smooth Muscle Cells Illuminated by KLF4 (Krüppel-Like Factor 4) Arteriosclerosis, Thrombosis, and Vascular Biology 41, 2693–2707 10.1161/ATVBAHA.121.316600

17. Yin, Y. W., Liao, S. Q., Zhang, M. J., Liu, Y., Li, B. H., Zhou, Y. et al. (2014) TLR4-mediated inflammation promotes foam cell formation of vascular smooth muscle cell by upregulating ACAT1 expression Cell Death Dis 5, e1574 10.1038/cddis.2014.535

18. Akira, S., and Takeda, K. (2004) Toll-like receptor signalling Nature Reviews Immunology 4, 499–511 10.1038/nri1391

19. Klouche, M., Bhakdi, S., Hemmes, M., and Rose-John, S. (1999) Novel path to activation of vascular smooth muscle cells: up-regulation of gp130 creates an autocrine activation loop by IL-6 and its soluble receptor J Immunol 163, 4583–4589,

20. Doyon, P., Kizilay-Mancini, O., Dô, F., Huynh, D., Mayer, G., Lehoux, S., et al. (2024) Early IKKβ-dependent anabolic signature governs vascular smooth muscle cells fate and abdominal aortic aneurysm development bioRxiv 2024.2006.2012.598763 10.1101/2024.06.12.598763

21. Singh, B., Kosuru, R., Lakshmikanthan, S., Sorci-Thomas, M. G., Zhang, D. X., Sparapani, R. et al. (2021) Endothelial Rap1 (Ras-Association Proximate 1) Restricts Inflammatory Signaling to Protect From the Progression of Atherosclerosis Arteriosclerosis, Thrombosis, and Vascular Biology 41, 638–650 doi:10.1161/ATVBAHA.120.315401

22. Akiyama, M., Hasegawa, H., Hongu, T., Frohman, M. A., Harada, A., Sakagami, H., et al. (2014) Trans-regulation of oligodendrocyte myelination by neurons through small GTPase Arf6-regulated secretion of fibroblast growth factor-2 Nat Commun 5, 4744 10.1038/ncomms5744

23. Hongu, T., Funakoshi, Y., Fukuhara, S., Suzuki, T., Sakimoto, S., Takakura, N., et al. (2015) Arf6 regulates tumour angiogenesis and growth through HGF-induced endothelial β1 integrin recycling Nature Communications 6, 7925 10.1038/ncomms8925

24. Barnes, M. J., and Farndale, R. W. (1999) Collagens and atherosclerosis Exp Gerontol 34, 513–525 10.1016/s0531-5565(99)00038-8

25. Kuzan, A., Wisniewski, J., Maksymowicz, K., Kobielarz, M., Gamian, A., and Chwilkowska, A. (2021) Relationship between calcification, atherosclerosis and matrix proteins in the human aorta Folia Histochem Cytobiol 59, 8–21 10.5603/FHC.a2021.0002

26. Nitschke, Y., Weissen-Plenz, G., Terkeltaub, R., and Rutsch, F. (2011) Npp1 promotes atherosclerosis in ApoE knockout mice J Cell Mol Med 15, 2273–2283 10.1111/j.1582-4934.2011.01327.x

27. Huff, M. W., and Pickering, J. G. (2015) Can a Vascular Smooth Muscle–Derived Foam-Cell Really Change its Spots? Arteriosclerosis, Thrombosis, and Vascular Biology 35, 492–495 10.1161/ATVBAHA.115.305225

28. Zani, I. A., Stephen, S. L., Mughal, N. A., Russell, D., Homer-Vanniasinkam, S., Wheatcroft, S. B. et al. (2015) Scavenger receptor structure and function in health and disease Cells 4, 178–201 10.3390/cells4020178

29. Kiyan, Y., Tkachuk, S., Hilfiker-Kleiner, D., Haller, H., Fuhrman, B., and Dumler, I. (2014) OxLDL induces inflammatory responses in vascular smooth muscle cells via urokinase receptor association with CD36 and TLR4 J Mol Cell Cardiol 66, 72–82 10.1016/j.yjmcc.2013.11.005

30. Lee, K., Yim, J. H., Lee, H. K., and Pyo, S. (2016) Inhibition of VCAM-1 expression on mouse vascular smooth muscle cells by lobastin via downregulation of p38, ERK 1/2 and NF-κB signaling pathways Arch Pharm Res 39, 83–93 10.1007/s12272-015-0687-3

31. Ming Cao, W., Murao, K., Imachi, H., Sato, M., Nakano, T., Kodama, T., et al. (2001) Phosphatidylinositol 3-OH kinase-Akt/protein kinase B pathway mediates Gas6 induction of scavenger receptor a in immortalized human vascular smooth muscle cell line Arterioscler Thromb Vasc Biol 21, 1592–1597 10.1161/hq1001.097062

32. Schweitzer, J. K., Pietrini, S. D., and D’Souza-Schorey, C. (2009) ARF6-mediated endosome recycling reverses lipid accumulation defects in Niemann-Pick Type C disease PLoS One 4, e5193 10.1371/journal.pone.0005193

33. Twigg, M. W., Freestone, K., Homer-Vanniasinkam, S., and Ponnambalam, S. (2012) The LOX-1 Scavenger Receptor and Its Implications in the Treatment of Vascular Disease Cardiology Research and Practice 2012, 632408 10.1155/2012/632408

34. Claing, A., Perry, S. J., Achiriloaie, M., Walker, J. K. L., Albanesi, J. P., Lefkowitz, R. J. et al. (2000) Multiple endocytic pathways of G protein-coupled receptors delineated by GIT1 sensitivity Proceedings of the National Academy of Sciences 97, 1119–1124 doi:10.1073/pnas.97.3.1119

35. Houndolo, T., Boulay, P.-L., and Claing, A. (2005) G Protein-coupled Receptor Endocytosis in ADP-ribosylation Factor 6-depleted Cells * Journal of Biological Chemistry 280, 5598–5604 10.1074/jbc.M411456200

36. Poupart, M. E., Fessart, D., Cotton, M., Laporte, S. A., and Claing, A. (2007) ARF6 regulates angiotensin II type 1 receptor endocytosis by controlling the recruitment of AP-2 and clathrin Cell Signal 19, 2370–2378 10.1016/j.cellsig.2007.07.015

37. Pasterkamp, G., Van Keulen, J. K., and De Kleijn, D. P. (2004) Role of Toll-like receptor 4 in the initiation and progression of atherosclerotic disease Eur J Clin Invest 34, 328–334 10.1111/j.1365-2362.2004.01338.x

38. Higashimori, M., Tatro, J. B., Moore, K. J., Mendelsohn, M. E., Galper, J. B., and Beasley, D. (2011) Role of Toll-Like Receptor 4 in Intimal Foam Cell Accumulation in Apolipoprotein E–Deficient Mice Arteriosclerosis, Thrombosis, and Vascular Biology 31, 50–57 doi:10.1161/ATVBAHA.110.210971

39. Van Acker, T., Eyckerman, S., Vande Walle, L., Gerlo, S., Goethals, M., Lamkanfi, M. et al. (2014) The small GTPase Arf6 is essential for the Tram/Trif pathway in TLR4 signaling J Biol Chem 289, 1364–1376 10.1074/jbc.M113.499194

40. Bäck, M., Yurdagul, A., Tabas, I., Öörni, K., and Kovanen, P. T. (2019) Inflammation and its resolution in atherosclerosis: mediators and therapeutic opportunities Nature Reviews Cardiology 16, 389–406 10.1038/s41569-019-0169-2

41. Čejková, S., Králová-Lesná, I., and Poledne, R. (2016) Monocyte adhesion to the endothelium is an initial stage of atherosclerosis development Cor et Vasa 58, e419–e425 10.1016/j.crvasa.2015.08.002

42. Ley, K., Miller, Y. I., and Hedrick, C. C. (2011) Monocyte and macrophage dynamics during atherogenesis Arteriosclerosis, thrombosis, and vascular biology 31, 1506–1516 10.1161/ATVBAHA.110.221127

43. Shen, Z., Li, C., Frieler, R. A., Gerasimova, A. S., Lee, S. J., Wu, J. et al. (2012) Smooth muscle protein 22 alpha-Cre is expressed in myeloid cells in mice Biochem Biophys Res Commun 422, 639–642 10.1016/j.bbrc.2012.05.041

44. Chakraborty, R., Saddouk, F. Z., Carrao, A. C., Krause, D. S., Greif, D. M., and Martin, K. A. (2019) Promoters to Study Vascular Smooth Muscle Arteriosclerosis, Thrombosis, and Vascular Biology 39, 603–612 doi:10.1161/ATVBAHA.119.312449

45. Poller, W. C., Nahrendorf, M., and Swirski, F. K. (2020) Hematopoiesis and Cardiovascular Disease Circulation Research 126, 1061–1085 10.1161/CIRCRESAHA.120.315895

46. Ridker, P. M., and Rane, M. (2021) Interleukin-6 Signaling and Anti-Interleukin-6 Therapeutics in Cardiovascular Disease Circulation Research 128, 1728–1746 doi:10.1161/CIRCRESAHA.121.319077

47. Charles, R., Bourmoum, M., Campbell, S., and Claing, A. (2019) Methods to Investigate the β-Arrestin-Mediated Control of ARF6 Activation to Regulate Trafficking and Actin Cytoskeleton Remodeling Methods Mol Biol 1957, 159–168 10.1007/978-1-4939-9158-7_10

48. Yuan, X. M., Li, W., Olsson, A. G., and Brunk, U. T. (1997) The toxicity to macrophages of oxidized low-density lipoprotein is mediated through lysosomal damage Atherosclerosis 133, 153–161 10.1016/S0021-9150(97)00094-4

49. Centa, M., Jin, H., Hofste, L., Hellberg, S., Busch, A., Baumgartner, R. et al. (2019) Germinal Center–Derived Antibodies Promote Atherosclerosis Plaque Size and Stability Circulation 139, 2466–2482 10.1161/CIRCULATIONAHA.118.038534

