## Supplemental information for "ARF6 controls VSMC cell phenotypic switching upon lipid stimulation to promote inflammatory signaling contributing to the progression of atherosclerosis"

From the [1]Department of Pharmacology and Physiology, Faculty of Medicine, Université de Montréal, Montreal, Quebec H3T 1J4, Canada; [2]Department of Physiological Chemistry, Faculty of Medical Sciences and Graduate School of Comprehensive Human Sciences, University of Tsukuba, 1-1-1 Tennohdai, Tsukuba 305- 8575, Japan, [3]Faculty of Pharmacy, Université de Montréal, Montreal, Quebec H3T 1J4, Canada.

***Abstract***

Vascular smooth muscle cells (VSMCs) play a pivotal role in the development and progression of atherosclerosis. Traditionally viewed as contractile cells that maintain vascular tone and structure, VSMCs undergo phenotypic switching in response to atherogenic stimuli, such as high circulating levels of LDL, thus adopting synthetic, osteogenic, or macrophage-like phenotypes. This plasticity contributes to plaque formation, extracellular matrix remodeling, and inflammatory signaling. We have previously shown that ADP-ribosylation factor 1 (ARF1), a small GTP-binding protein, regulates the expression and function of actin, which is important for maintaining the contractile phenotype of VSMCs. However, the role of ARF6 in phenotypic switching remains to be elucidated. Here, we demonstrate that ARF6 knockdown in human aortic smooth muscle cells (HASMCs) reduced lipid uptake through alterations in the expression of scavenger receptors (LOX-1, MSR1), cytokine production (IL-6) as well as modulation of inflammatory markers and pathways (adhesion molecules, PI3K, NFkB, p38). To confirm our findings in an *in vivo* setting, we engineered a novel conditional smooth muscle cell-specific ARF6 KO mouse in an atheroprone background (*Acta2-Cre-ERT2*$^{+/-}$/*ApoE*$^{-/-}$/*Arf6*$^{f/f}$ mice). Mice were fed a high-fat diet to accelerate plaque formation. ARF6 knockout resulted in a significant reduction of atherosclerotic lesions in the aortic arch, which was associated with a reduction of collagen and foam cells. Furthermore, we observed that ARF6 regulates the expression of inflammatory markers. These findings highlight the importance of ARF6 expression in VSMCs and its role in the pathogenesis of atherosclerosis.

### Supporting Information S1

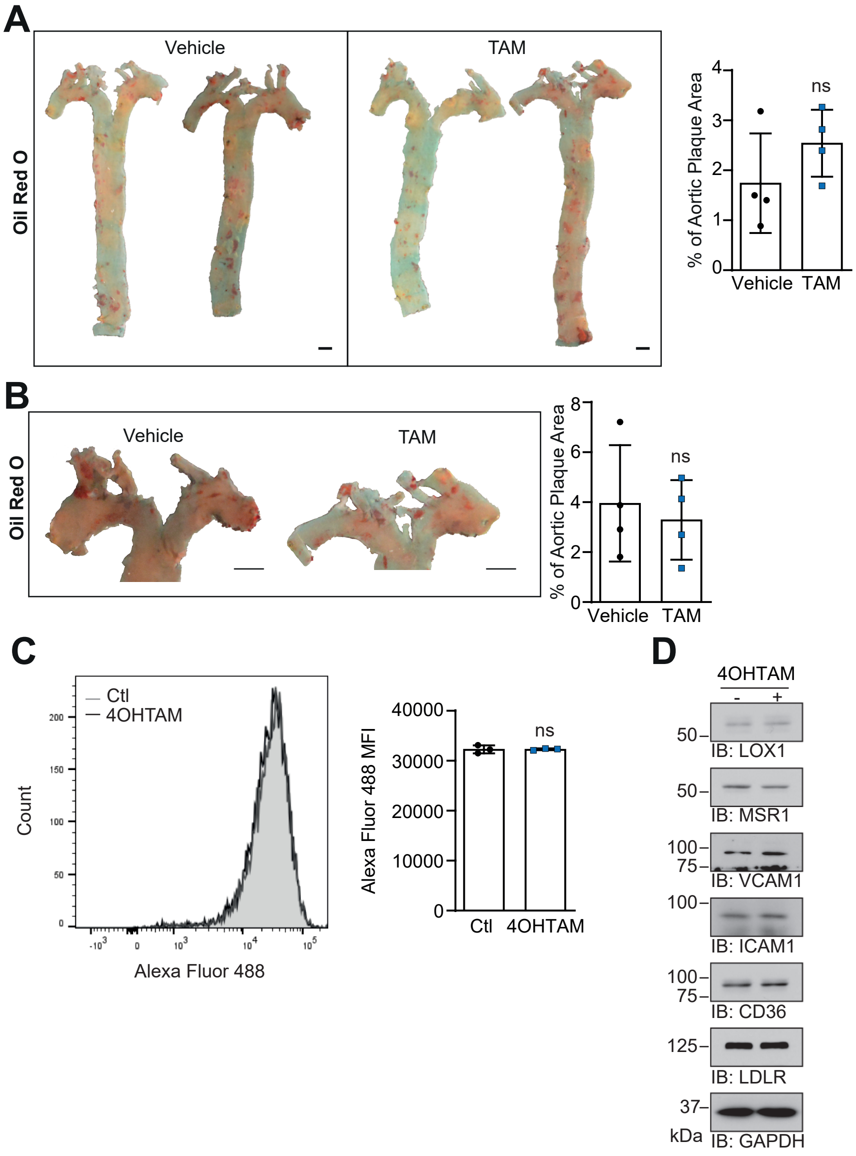


Figure S1 Tamoxifen does not alter atheroma formation or lipid uptake. A,B) Representative images of en-face aorta (A) or aortic arch (B) stained with Oil Red O (ORO). Quantifications are the mean ± SD of four animals per group. Ns not significant (Unpaired Student's t-test). Bar scale: 1 mm. C) Cells were pretreated with DMSO (Ctl) or 4-Hydroxytamoxifen (4OHTAM) (1 µM) for 72 h before incubation with Alexa Fluor 488-labelled AcLDL (1 µg/ml) for 18 h and analyzed by flow cytometry. Quantifications are mean ± SD of mean fluorescence intensity (MFI) of triplicates from one experiment, representative of three independent experiments. ns not significant (Unpaired Student's t-test). D) Cells were treated with DMSO (-) or 4-Hydroxytamoxifen (+) (4OHTAM) (1 µM) for 72 h and stimulated with oxLDL (80 µg/ml) for 72 h. Expression levels of indicated proteins were assessed by Western blotting using specific antibodies. Images are representative of three independent experiments.

### Supporting Information S2

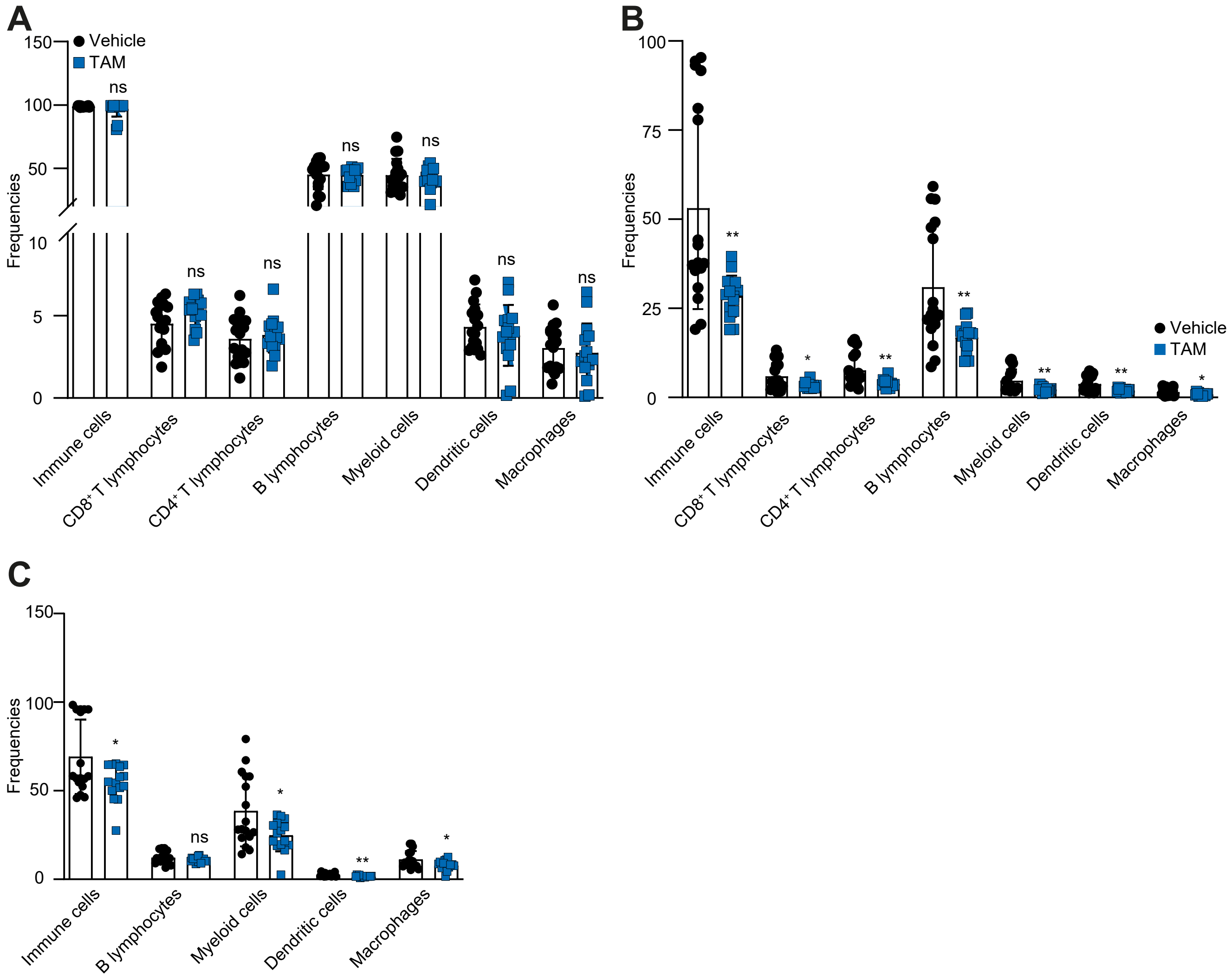


Figure S2 Leucocyte profiles of control and ARF6 KO mice. Frequencies of immune cell populations were assessed in the (A) blood, (B) spleen, and (C) bone marrow in Vehicle or Tamoxifen (TAM) treated mice after ten weeks of high-fat diet. Immune cell frequencies were analysed after exclusion of doublets and dead cells. Immune cells are defined as CD45+ cells, CD8+ T lymphocytes as CD8+ cells, CD4+ T lymphocyte as CD4+ cells, B lymphocytes as CD19+ cells, myeloid cells as CD11b+ cells, dendritic cells as CD11c+ cells, and macrophages as CD11b and F4/80 double positive cells. Quantifications represent the frequency in mean ± SD of 15 or 17 animals by group. **P<0.01,*P<0.05, ns not significant (Unpaired Student's t-test).

| # | Visible? | Starred? | Bio View:Ident | Accession Nu | Alternate ID | Molecular We | Protein Group | SCR | shARF6 |
|---|---|---|---|---|---|---|---|---|---|
| | VRAI | Orange | Collagen alph | CO1A2_HUM/ | COL1A2 | 129 kDa | | 160 | 101 |
| | VRAI | Orange | Collagen alph | CO1A1_HUM/ | COL1A1 | 139 kDa | | 44 | 25 |
| | VRAI | Orange | Collagen alph | CO5A1_HUM/ | COL5A1 | 184 kDa | | 14 | 7 |
| | VRAI | Orange | Collagen alph | CO5A2_HUM/ | COL5A2 | 145 kDa | | 12 | 5 |
| | VRAI | Orange | Collagen alph | CO4A2_HUM/ | COL4A2 | 168 kDa | | 13 | 3 |
| | VRAI | Orange | Collagen alph | CO3A1_HUM/ | COL3A1 | 139 kDa | | 8 | 5 |
| | VRAI | Orange | Collagen alph | CO6A1_HUM/ | COL6A1 | 109 kDa | | 6 | 5 |
| | VRAI | Orange | Collagen alph | CO6A3_HUM/ | COL6A3 | 344 kDa | | 8 | 2 |
| | VRAI | Empty | Complement | CFAH_HUMAI | CFH | 139 kDa | | 28 | 18 |
| | VRAI | Empty | Glucose-6-ph | G6PI_HUMAN | GPI | 63 kDa | | 20 | 22 |
| | VRAI | Empty | Decorin OS=H | PGS2_HUMAI | DCN | 40 kDa | | 20 | 20 |
| | VRAI | Empty | Heat shock cc | HSP7C_HUM/ | HSPA8 | 71 kDa | VRAI | 11 | 11 |
| | VRAI | Empty | 72 kDa type IV | MMP2_HUMA | MMP2 | 74 kDa | | 8 | 9 |
| | VRAI | Empty | Interstitial col | MMP1_HUMA | MMP1 | 54 kDa | | 9 | 5 |
| | VRAI | Empty | Versican core | CSPG2_HUM/ | VCAN | 373 kDa | | 11 | 3 |
| | VRAI | Empty | Procollagen-ly | PLOD2_HUM/ | PLOD2 | 85 kDa | | 7 | 5 |
| | VRAI | Empty | Glycogen pho | PYGB_HUMAI | PYGB | 97 kDa | | 6 | 5 |
| | VRAI | Empty | WD repeat-co | WDR1_HUMA | WDR1 | 66 kDa | | 4 | 7 |
| | VRAI | Empty | Procollagen-ly | PLOD1_HUM/ | PLOD1 | 84 kDa | | 3 | 5 |
| | VRAI | Empty | Staphylococc | SND1_HUMAI | SND1 | 102 kDa | | 2 | 3 |
| | VRAI | Empty | EGF-containir | FBLN3_HUMA | EFEMP1 | 55 kDa | | 3 | 2 |
| | VRAI | Empty | Glucose-6-ph | G6PD_HUMAI | G6PD | 59 kDa | | 2 | 3 |
| | VRAI | Empty | Plexin domain | PXDC2_HUM/ | PLXDC2 | 60 kDa | | 2 | 3 |
| | VRAI | Empty | N-sulphogluc | SPHM_HUMAI | SGSH | 57 kDa | | 2 | 2 |
| | VRAI | Empty | Complement | CO3_HUMAN | C3 | 187 kDa | | 2 | 1 |
| | VRAI | Empty | Basement me | PGBM_HUMAI | HSPG2 | 469 kDa | | 3 | 1 |
| | VRAI | Empty | Complement | C1S_HUMAN | C1S | 77 kDa | | 3 | 1 |
| | VRAI | Empty | Neutral alpha | GANAB_HUM/ | GANAB | 107 kDa | | 3 | 1 |
| | VRAI | Empty | Alpha-2-HS-gl | FETUA_HUMA | AHSG | 39 kDa | | 2 | 1 |
| | VRAI | Empty | Glucosidase 2 | GLU2B_HUM/ | PRKCSH | 59 kDa | | 1 | 1 |
| | VRAI | Empty | EGF-like repea | EDIL3_HUMAI | EDIL3 | 54 kDa | | 0 | 1 |
| | VRAI | Empty | Coronin-1C O | COR1C_HUM. | CORO1C | 53 kDa | | | 2 |
| | VRAI | Empty | T-complex prc | TCPE_HUMAN | CCT5 | 60 kDa | | | 2 |
| | VRAI | Empty | Macrophage c | CSF1_HUMAN | CSF1 | 60 kDa | | 2 | |

END OF FILE

#Audrey Caling SCR_ARF6 010319 (7), Samples Report created on 07/14/2025

| | | | | | | | | | Percent Cover | Percent Coverage |
|---|---|---|---|---|---|---|---|---|---|---|
| Audrey Caling SCR_ARF6 010319 (7) | | | | | | | | | SCR | shARF6 |
| # | Visible? | Starred? | Identified Prot | Accession Nu | Alternate ID | Molecular We | Protein Group | Taxonomy | SCR | shARF6 |
| 27 | VRAI | Orange | Collagen alph | CO1A2_HUM/ | COL1A2 | 129 kDa | | unknown | 57.70% | 52.00% |
| 31 | VRAI | Orange | Collagen alph | CO1A1_HUM/ | COL1A1 | 139 kDa | | unknown | 25.70% | 15.20% |
| 21 | VRAI | Empty | Complement | CFAH_HUMAI | CFH | 139 kDa | | unknown | 17.70% | 15.70% |
| 15 | VRAI | Empty | Glucose-6-ph | G6PI_HUMAN | GPI | 63 kDa | | unknown | 31.00% | 24.00% |
| 19 | VRAI | Empty | Decorin OS=H | PGS2_HUMAN | DCN | 40 kDa | | unknown | 37.00% | 37.30% |
| 12 | VRAI | Empty | Heat shock co | HSP7C_HUM/ | HSPA8 | 71 kDa | VRAI | unknown | 14.70% | 14.70% |
| 29 | VRAI | Orange | Collagen alph | CO5A1_HUM/ | COL5A1 | 184 kDa | | unknown | 7.78% | 4.68% |
| 25 | VRAI | Orange | Collagen alph | CO5A2_HUM/ | COL5A2 | 145 kDa | | unknown | 10.40% | 7.47% |
| 34 | VRAI | Empty | 72 kDa type IV | MMP2_HUMA | MMP2 | 74 kDa | | unknown | 12.10% | 9.24% |
| 26 | VRAI | Orange | Collagen alph | CO4A2_HUM/ | COL4A2 | 168 kDa | | unknown | 6.78% | 3.04% |
| 11 | VRAI | Empty | Interstitial col | MMP1_HUMA | MMP1 | 54 kDa | | unknown | 20.30% | 10.40% |
| 30 | VRAI | Orange | Collagen alph | CO3A1_HUM/ | COL3A1 | 139 kDa | | unknown | 5.46% | 5.05% |
| 2 | VRAI | Empty | Versican core | CSPG2_HUM/ | VCAN | 373 kDa | | unknown | 3.47% | 1.62% |
| 28 | VRAI | Orange | Collagen alph | CO6A1_HUM/ | COL6A1 | 109 kDa | | unknown | 7.10% | 5.64% |
| 5 | VRAI | Empty | Procollagen-ly | PLOD2_HUM/ | PLOD2 | 85 kDa | | unknown | 9.91% | 7.06% |
| 13 | VRAI | Empty | Glycogen pho | PYGB_HUMAI | PYGB | 97 kDa | | unknown | 5.93% | 5.93% |
| 1 | VRAI | Empty | WD repeat-co | WDR1_HUMA | WDR1 | 66 kDa | | unknown | 6.60% | 8.91% |
| 24 | VRAI | Orange | Collagen alph | CO6A3_HUM/ | COL6A3 | 344 kDa | | unknown | 2.80% | 0.57% |
| 6 | VRAI | Empty | Procollagen-ly | PLOD1_HUM/ | PLOD1 | 84 kDa | | unknown | 8.12% | 11.60% |
| 4 | VRAI | Empty | Staphylococc | SND1_HUMAI | SND1 | 102 kDa | | unknown | 2.53% | 3.41% |
| 18 | VRAI | Empty | EGF-containir | FBLN3_HUMA | EFEMP1 | 55 kDa | | unknown | 3.25% | 3.25% |
| 16 | VRAI | Empty | Glucose-6-ph | G6PD_HUMAI | G6PD | 59 kDa | | unknown | 5.05% | 7.96% |
| 7 | VRAI | Empty | Plexin domain | PXDC2_HUM/ | PLXDC2 | 60 kDa | | unknown | 5.10% | 5.10% |
| 9 | VRAI | Empty | N-sulphogluc | SPHM_HUMAI | SGSH | 57 kDa | | unknown | 4.78% | 4.78% |
| 22 | VRAI | Empty | Complement | CO3_HUMAN | C3 | 187 kDa | | unknown | 1.68% | 0.84% |
| 32 | VRAI | Empty | Basement me | PGBM_HUMA | HSPG2 | 469 kDa | | unknown | 0.71% | 0.25% |
| 23 | VRAI | Empty | Complement | C1S_HUMAN | C1S | 77 kDa | | unknown | 2.76% | 2.76% |
| 8 | VRAI | Empty | Neutral alpha- | GANAB_HUM/ | GANAB | 107 kDa | | unknown | 4.13% | 1.59% |
| 33 | VRAI | Empty | Alpha-2-HS-gl | FETUA_HUMA | AHSG | 39 kDa | | unknown | 3.54% | 3.54% |
| 14 | VRAI | Empty | Glucosidase 2 | GLU2B_HUM/ | PRKCSH | 59 kDa | | unknown | 1.52% | 1.89% |
| 17 | VRAI | Empty | EGF-like repe | EDIL3_HUMAI | EDIL3 | 54 kDa | | unknown | 0.00% | 2.08% |
| 20 | VRAI | Empty | Coronin-1C O | COR1C_HUM/ | CORO1C | 53 kDa | | unknown | 0 | 5.27% |
| 3 | VRAI | Empty | T-complex pro | TCPE_HUMAN | CCT5 | 60 kDa | | unknown | 0 | 8.13% |
| 10 | VRAI | Empty | Macrophage c | CSF1_HUMAN | CSF1 | 60 kDa | | unknown | 6.68% | 0 |

END OF FILE

| | | | | | | | | | | Percentage of | Percentage of Total Spectra |
|---|---|---|---|---|---|---|---|---|---|---|---|
| Audrey Caling | SCR_ARF6 010319 (7) | | | | | | | | | SCR | shARF6 |
| # | | Visible? | Starred? | Identified Proteins | Accession Nu | Alternate ID | Molecular We | Protein Group | Taxonomy | SCR | shARF6 |
| | 1 | VRAI | Orange | Collagen alpha-2( | CO1A2_HUM/ | COL1A2 | 129 kDa | | unknown | 5.90% | 4.30% |
| | 2 | VRAI | Orange | Collagen alpha-1( | CO1A1_HUM/ | COL1A1 | 139 kDa | | unknown | 1.60% | 1.10% |
| | 9 | VRAI | Empty | Complement fact | CFAH_HUMAI | CFH | 139 kDa | | unknown | 1.00% | 0.77% |
| | 10 | VRAI | Empty | Glucose-6-phospl | G6PI_HUMAN | GPI | 63 kDa | | unknown | 0.73% | 0.94% |
| | 11 | VRAI | Empty | Decorin OS=Hom | PGS2_HUMAI | DCN | 40 kDa | | unknown | 0.73% | 0.85% |
| | 12 | VRAI | Empty | Heat shock cogna | HSP7C_HUM/ | HSPA8 | 71 kDa | VRAI | unknown | 0.40% | 0.47% |
| | 3 | VRAI | Orange | Collagen alpha-1( | CO5A1_HUM/ | COL5A1 | 184 kDa | | unknown | 0.51% | 0.30% |
| | 4 | VRAI | Orange | Collagen alpha-2( | CO5A2_HUM/ | COL5A2 | 145 kDa | | unknown | 0.44% | 0.21% |
| | 13 | VRAI | Empty | 72 kDa type IV col | MMP2_HUMA | MMP2 | 74 kDa | | unknown | 0.29% | 0.38% |
| | 5 | VRAI | Orange | Collagen alpha-2( | CO4A2_HUM/ | COL4A2 | 168 kDa | | unknown | 0.48% | 0.13% |
| | 14 | VRAI | Empty | Interstitial collage | MMP1_HUMA | MMP1 | 54 kDa | | unknown | 0.33% | 0.21% |
| | 6 | VRAI | Orange | Collagen alpha-1( | CO3A1_HUM/ | COL3A1 | 139 kDa | | unknown | 0.29% | 0.21% |
| | 15 | VRAI | Empty | Versican core pro | CSPG2_HUM/ | VCAN | 373 kDa | | unknown | 0.40% | 0.13% |
| | 7 | VRAI | Orange | Collagen alpha-1( | CO6A1_HUM/ | COL6A1 | 109 kDa | | unknown | 0.22% | 0.21% |
| | 16 | VRAI | Empty | Procollagen-lysin | PLOD2_HUM/ | PLOD2 | 85 kDa | | unknown | 0.26% | 0.21% |
| | 17 | VRAI | Empty | Glycogen phosph | PYGB_HUMAI | PYGB | 97 kDa | | unknown | 0.22% | 0.21% |
| | 18 | VRAI | Empty | WD repeat-contai | WDR1_HUMA | WDR1 | 66 kDa | | unknown | 0.15% | 0.30% |
| | 8 | VRAI | Orange | Collagen alpha-3( | CO6A3_HUM/ | COL6A3 | 344 kDa | | unknown | 0.29% | 0.09% |
| | 19 | VRAI | Empty | Procollagen-lysin | PLOD1_HUM/ | PLOD1 | 84 kDa | | unknown | 0.11% | 0.21% |
| | 20 | VRAI | Empty | Staphylococcal nu | SND1_HUMAI | SND1 | 102 kDa | | unknown | 0.07% | 0.13% |
| | 21 | VRAI | Empty | EGF-containing fit | FBLN3_HUMA | EFEMP1 | 55 kDa | | unknown | 0.11% | 0.09% |
| | 22 | VRAI | Empty | Glucose-6-phospl | G6PD_HUMAI | G6PD | 59 kDa | | unknown | 0.07% | 0.13% |
| | 23 | VRAI | Empty | Plexin domain-co | PXDC2_HUM/ | PLXDC2 | 60 kDa | | unknown | 0.07% | 0.13% |
| | 24 | VRAI | Empty | N-sulphoglucosar | SPHM_HUMAI | SGSH | 57 kDa | | unknown | 0.07% | 0.09% |
| | 25 | VRAI | Empty | Complement C3 ( | CO3_HUMAN | C3 | 187 kDa | | unknown | 0.07% | 0.04% |
| | 26 | VRAI | Empty | Basement membr | PGBM_HUMA | HSPG2 | 469 kDa | | unknown | 0.11% | 0.04% |
| | 27 | VRAI | Empty | Complement C1s | C1S_HUMAN | C1S | 77 kDa | | unknown | 0.11% | 0.04% |
| | 28 | VRAI | Empty | Neutral alpha-glu | GANAB_HUM/ | GANAB | 107 kDa | | unknown | 0.11% | 0.04% |
| | 29 | VRAI | Empty | Alpha-2-HS-glyco | FETUA_HUMA | AHSG | 39 kDa | | unknown | 0.07% | 0.04% |
| | 30 | VRAI | Empty | Glucosidase 2 sul | GLU2B_HUM/ | PRKCSH | 59 kDa | | unknown | 0.04% | 0.04% |
| | 31 | VRAI | Empty | EGF-like repeat ar | EDIL3_HUMAI | EDIL3 | 54 kDa | | unknown | 0.00% | 0.04% |
| | 32 | VRAI | Empty | Coronin-1C OS=H | COR1C_HUM, | CORO1C | 53 kDa | | unknown | 0 | 0.09% |
| | 33 | VRAI | Empty | T-complex protein | TCPE_HUMAN | CCT5 | 60 kDa | | unknown | 0 | 0.09% |
| | 34 | VRAI | Empty | Macrophage color | CSF1_HUMAN | CSF1 | 60 kDa | | unknown | 0.07% | 0 |

| Experiment na | Biological sar | Biological sar | MS/MS sampl | Protein group | Protein acces | Protein name | Protein identi | Protein perce | Protein molec | Number of un | Number of un | Number of tot | Spectrum nar | Peptide seque | Previous amir | Next amino ac | Peptide identi | Modifications | Actual peptid | Spectrum cha | Actual minus | Actual minus | Peptide start i | Peptide stop index |
|---|---|---|---|---|---|---|---|---|---|---|---|---|---|---|---|---|---|---|---|---|---|---|---|---|
| Audrey Caling | SCR | SCR | Service_Clair | Collagen alph | CO1A1_HUM | Collagen alph | 100.00% | 1.61% | 138,923.40 | 28 | 35 | 44 | index=9951 | ADDANVVRD | R | S | 99.60% | | 2,144.08 | 4 | 0.01 | 4.7 | 1218 | 1236 |
| Audrey Caling | SCR | SCR | Service_Clair | Collagen alph | CO1A1_HUM | Collagen alph | 100.00% | 1.61% | 138,923.40 | 28 | 35 | 44 | index=9959 | ADDANVVRD | R | S | 99.70% | | 2,144.08 | 3 | 0.0086 | 4 | 1218 | 1236 |
| Audrey Caling | SCR | SCR | Service_Clair | Collagen alph | CO1A1_HUM | Collagen alph | 100.00% | 1.61% | 138,923.40 | 28 | 35 | 44 | index=6791 | GERGFPGER | R | G | 99.60% | | 2,074.06 | 4 | 0.0092 | 4.4 | 677 | 697 |
| Audrey Caling | SCR | SCR | Service_Clair | Collagen alph | CO1A1_HUM | Collagen alph | 100.00% | 1.61% | 138,923.40 | 28 | 35 | 44 | index=2436 | GESGPSGPA | K | G | 99.70% | | 1,296.61 | 2 | 0.0027 | 2.1 | 782 | 796 |
| Audrey Caling | SCR | SCR | Service_Clair | Collagen alph | CO1A1_HUM | Collagen alph | 100.00% | 1.61% | 138,923.40 | 28 | 35 | 44 | index=7847 | GFPGADGVA | R | G | 99.70% | | 1,071.54 | 2 | 0.0023 | 2.1 | 494 | 505 |
| Audrey Caling | SCR | SCR | Service_Clair | Collagen alph | CO1A1_HUM | Collagen alph | 100.00% | 1.61% | 138,923.40 | 28 | 35 | 44 | index=7846 | GFPGADGVA | R | G | 99.70% | | 1,071.54 | 2 | 0.0023 | 2.1 | 494 | 505 |
| Audrey Caling | SCR | SCR | Service_Clair | Collagen alph | CO1A1_HUM | Collagen alph | 100.00% | 1.61% | 138,923.40 | 28 | 35 | 44 | index=7711 | GFPGADGVA | R | G | 99.70% | | 1,638.82 | 3 | 0.0063 | 3.8 | 494 | 511 |
| Audrey Caling | SCR | SCR | Service_Clair | Collagen alph | CO1A1_HUM | Collagen alph | 100.00% | 1.61% | 138,923.40 | 28 | 35 | 44 | index=7585 | GFPGERGVQ | R | G | 99.70% | | 1,731.89 | 3 | 0.0064 | 3.7 | 680 | 697 |
| Audrey Caling | SCR | SCR | Service_Clair | Collagen alph | CO1A1_HUM | Collagen alph | 100.00% | 1.61% | 138,923.40 | 28 | 35 | 44 | index=7584 | GFPGERGVQ | R | G | 99.70% | | 1,731.89 | 3 | 0.0064 | 3.7 | 680 | 697 |
| Audrey Caling | SCR | SCR | Service_Clair | Collagen alph | CO1A1_HUM | Collagen alph | 100.00% | 1.61% | 138,923.40 | 28 | 35 | 44 | index=3993 | GLPGERGRP | R | G | 98.10% | | 1,671.90 | 4 | 0.0039 | 2.3 | 305 | 322 |
| Audrey Caling | SCR | SCR | Service_Clair | Collagen alph | CO1A1_HUM | Collagen alph | 100.00% | 1.61% | 138,923.40 | 28 | 35 | 44 | index=3992 | GLPGERGRP | R | G | 99.70% | | 1,671.90 | 4 | 0.0039 | 2.3 | 305 | 322 |
| Audrey Caling | SCR | SCR | Service_Clair | Collagen alph | CO1A1_HUM | Collagen alph | 100.00% | 1.61% | 138,923.40 | 28 | 35 | 44 | index=9653 | GLTGPIGPPG | R | G | 99.70% | | 2,836.42 | 3 | 0.0099 | 3.5 | 764 | 796 |
| Audrey Caling | SCR | SCR | Service_Clair | Collagen alph | CO1A1_HUM | Collagen alph | 100.00% | 1.61% | 138,923.40 | 28 | 35 | 44 | index=3728 | GLTGSPGSP | K | T | 99.70% | | 1,225.60 | 2 | 0.0025 | 2 | 539 | 552 |
| Audrey Caling | SCR | SCR | Service_Clair | Collagen alph | CO1A1_HUM | Collagen alph | 100.00% | 1.61% | 138,923.40 | 28 | 35 | 44 | index=1251 | GPAGPQGPR | R | G | 99.70% | | 2,064.96 | 4 | 0.0066 | 3.2 | 1085 | 1105 |
| Audrey Caling | SCR | SCR | Service_Clair | Collagen alph | CO1A1_HUM | Collagen alph | 100.00% | 1.61% | 138,923.40 | 28 | 35 | 44 | index=2810 | GPPGPPGKN | R | G | 98.90% | Deamidated ( | 3,116.52 | 5 | 0.0056 | 1.8 | 221 | 253 |
| Audrey Caling | SCR | SCR | Service_Clair | Collagen alph | CO1A1_HUM | Collagen alph | 100.00% | 1.61% | 138,923.40 | 28 | 35 | 44 | index=3059 | GPSGPQGPG | R | G | 99.70% | | 1,285.65 | 2 | 0.0044 | 3.4 | 416 | 430 |
| Audrey Caling | SCR | SCR | Service_Clair | Collagen alph | CO1A1_HUM | Collagen alph | 100.00% | 1.61% | 138,923.40 | 28 | 35 | 44 | index=11151 | GQAGVMGFP | R | G | 98.40% | | 1,144.57 | 2 | 0.0028 | 2.4 | 575 | 586 |
| Audrey Caling | SCR | SCR | Service_Clair | Collagen alph | CO1A1_HUM | Collagen alph | 100.00% | 1.61% | 138,923.40 | 28 | 35 | 44 | index=1371 | GRPGAPGPA | R | G | 99.70% | | 1,062.57 | 2 | 0.0023 | 2.2 | 311 | 322 |
| Audrey Caling | SCR | SCR | Service_Clair | Collagen alph | CO1A1_HUM | Collagen alph | 100.00% | 1.61% | 138,923.40 | 28 | 35 | 44 | index=1364 | GRPGAPGPA | R | G | 99.70% | | 1,062.57 | 3 | 0.0026 | 2.4 | 311 | 322 |
| Audrey Caling | SCR | SCR | Service_Clair | Collagen alph | CO1A1_HUM | Collagen alph | 100.00% | 1.61% | 138,923.40 | 28 | 35 | 44 | index=17125 | GRTGDAGPV | R | Y | 99.40% | Phospho (+80 | 4,533.16 | 5 | 0.027 | 6 | 1169 | 1214 |
| Audrey Caling | SCR | SCR | Service_Clair | Collagen alph | CO1A1_HUM | Collagen alph | 100.00% | 1.61% | 138,923.40 | 28 | 35 | 44 | index=17124 | GRTGDAGPV | R | Y | 99.70% | Phospho (+80 | 4,533.16 | 5 | 0.027 | 6 | 1169 | 1214 |
| Audrey Caling | SCR | SCR | Service_Clair | Collagen alph | CO1A1_HUM | Collagen alph | 100.00% | 1.61% | 138,923.40 | 28 | 35 | 44 | index=17126 | GRTGDAGPV | R | Y | 99.70% | Phospho (+80 | 4,533.16 | 5 | 0.027 | 6 | 1169 | 1214 |
| Audrey Caling | SCR | SCR | Service_Clair | Collagen alph | CO1A1_HUM | Collagen alph | 100.00% | 1.61% | 138,923.40 | 28 | 35 | 44 | index=17130 | GRTGDAGPV | R | Y | 99.70% | Phospho (+80 | 4,533.15 | 4 | 0.025 | 5.4 | 1169 | 1214 |
| Audrey Caling | SCR | SCR | Service_Clair | Collagen alph | CO1A1_HUM | Collagen alph | 100.00% | 1.61% | 138,923.40 | 28 | 35 | 44 | index=17123 | GRTGDAGPV | R | Y | 99.70% | Phospho (+80 | 4,533.16 | 5 | 0.027 | 6 | 1169 | 1214 |
| Audrey Caling | SCR | SCR | Service_Clair | Collagen alph | CO1A1_HUM | Collagen alph | 100.00% | 1.61% | 138,923.40 | 28 | 35 | 44 | index=7848 | GSAGPPGAT | R | V | 99.70% | | 1,426.70 | 2 | 0.0039 | 2.7 | 866 | 882 |
| Audrey Caling | SCR | SCR | Service_Clair | Collagen alph | CO1A1_HUM | Collagen alph | 100.00% | 1.61% | 138,923.40 | 28 | 35 | 44 | index=7849 | GSAGPPGAT | R | V | 99.70% | | 1,426.70 | 2 | 0.0039 | 2.7 | 866 | 882 |
| Audrey Caling | SCR | SCR | Service_Clair | Collagen alph | CO1A1_HUM | Collagen alph | 100.00% | 1.61% | 138,923.40 | 28 | 35 | 44 | index=5970 | GVPGPPGAV | R | D | 99.70% | | 1,159.64 | 2 | 0.0038 | 3.3 | 599 | 612 |
| Audrey Caling | SCR | SCR | Service_Clair | Collagen alph | CO1A1_HUM | Collagen alph | 100.00% | 1.61% | 138,923.40 | 28 | 35 | 44 | index=3678 | GVQGPPGPA | R | G | 99.70% | | 1,088.57 | 2 | 0.0017 | 1.5 | 686 | 697 |
| Audrey Caling | SCR | SCR | Service_Clair | Collagen alph | CO1A1_HUM | Collagen alph | 100.00% | 1.61% | 138,923.40 | 28 | 35 | 44 | index=3677 | GVQGPPGPA | R | G | 99.70% | | 1,088.57 | 2 | 0.0017 | 1.5 | 686 | 697 |
| Audrey Caling | SCR | SCR | Service_Clair | Collagen alph | CO1A1_HUM | Collagen alph | 100.00% | 1.61% | 138,923.40 | 28 | 35 | 44 | index=6498 | GVVGLPGQR | R | G | 99.70% | | 1,223.68 | 2 | 0.0033 | 2.7 | 959 | 970 |
| Audrey Caling | SCR | SCR | Service_Clair | Collagen alph | CO1A1_HUM | Collagen alph | 100.00% | 1.61% | 138,923.40 | 28 | 35 | 44 | index=10637 | SLSQQIENIR | K | K | 99.70% | | 1,799.92 | 2 | 0.0036 | 2 | 1237 | 1252 |
| Audrey Caling | SCR | SCR | Service_Clair | Collagen alph | CO1A1_HUM | Collagen alph | 100.00% | 1.61% | 138,923.40 | 28 | 35 | 44 | index=10630 | SLSQQIENIR | K | K | 99.70% | | 1,799.92 | 3 | 0.0046 | 2.6 | 1237 | 1252 |
| Audrey Caling | SCR | SCR | Service_Clair | Collagen alph | CO1A1_HUM | Collagen alph | 100.00% | 1.61% | 138,923.40 | 28 | 35 | 44 | index=10652 | STGGISVPGP | K | G | 99.70% | | 1,552.77 | 2 | 0.0032 | 2 | 171 | 187 |
| Audrey Caling | SCR | SCR | Service_Clair | Collagen alph | CO1A1_HUM | Collagen alph | 100.00% | 1.61% | 138,923.40 | 28 | 35 | 44 | index=8093 | STGGISVPGP | K | G | 99.70% | Oxidation (+1 | 1,568.77 | 2 | 0.0043 | 2.8 | 171 | 187 |
| Audrey Caling | SCR | SCR | Service_Clair | Collagen alph | CO1A1_HUM | Collagen alph | 100.00% | 1.61% | 138,923.40 | 28 | 35 | 44 | index=17561 | TGDAGPVGPI | R | Y | 95.60% | Phospho (+80 | 4,320.03 | 4 | 0.02 | 4.7 | 1171 | 1214 |
| Audrey Caling | SCR | SCR | Service_Clair | Collagen alph | CO1A1_HUM | Collagen alph | 100.00% | 1.61% | 138,923.40 | 28 | 35 | 44 | index=4900 | TGPPGPAGQI | K | G | 99.70% | | 1,991.99 | 3 | 0.0012 | 0.6 | 553 | 574 |
| Audrey Caling | SCR | SCR | Service_Clair | Collagen alph | CO1A1_HUM | Collagen alph | 100.00% | 1.61% | 138,923.40 | 28 | 35 | 44 | index=4093 | TVIEYKTTK | K | T | 99.70% | | 1,081.61 | 2 | 0.0038 | 3.5 | 1425 | 1433 |
| Audrey Caling | SCR | SCR | Service_Clair | Collagen alph | CO1A1_HUM | Collagen alph | 100.00% | 1.61% | 138,923.40 | 28 | 35 | 44 | index=4069 | TVIEYKTTK | K | T | 99.70% | | 1,081.60 | 3 | 0.002 | 1.8 | 1425 | 1433 |
| Audrey Caling | SCR | SCR | Service_Clair | Collagen alph | CO1A1_HUM | Collagen alph | 100.00% | 1.61% | 138,923.40 | 28 | 35 | 44 | index=3244 | TVIEYKTTKTS | K | L | 99.70% | | 1,425.79 | 3 | 0.0046 | 3.3 | 1425 | 1436 |
| Audrey Caling | SCR | SCR | Service_Clair | Collagen alph | CO1A1_HUM | Collagen alph | 100.00% | 1.61% | 138,923.40 | 28 | 35 | 44 | index=5449 | YHDRDVWKP | R | I | 99.40% | Carbamidom | 1,756.81 | 4 | 0.0022 | 1.2 | 47 | 59 |
| Audrey Caling | SCR | SCR | Service_Clair | Collagen alph | CO1A1_HUM | Collagen alph | 100.00% | 1.61% | 138,923.40 | 28 | 35 | 44 | index=4793 | YYRADDANV | R | D | 99.70% | | 1,340.65 | 3 | 0.0032 | 2.4 | 1215 | 1225 |
| Audrey Caling | SCR | SCR | Service_Clair | Collagen alph | CO1A1_HUM | Collagen alph | 100.00% | 1.61% | 138,923.40 | 28 | 35 | 44 | index=10314 | YYRADDANV | R | S | 96.60% | | 2,626.31 | 4 | 0.0093 | 3.5 | 1215 | 1236 |
| Audrey Caling | SCR | SCR | Service_Clair | Collagen alph | CO1A1_HUM | Collagen alph | 100.00% | 1.61% | 138,923.40 | 28 | 35 | 44 | index=10313 | YYRADDANV | R | S | 99.30% | | 2,626.31 | 4 | 0.0093 | 3.5 | 1215 | 1236 |
| Audrey Caling | SCR | SCR | Service_Clair | Collagen alph | CO1A1_HUM | Collagen alph | 100.00% | 1.61% | 138,923.40 | 28 | 35 | 44 | index=10318 | YYRADDANV | R | S | 99.70% | | 2,626.31 | 3 | 0.01 | 3.9 | 1215 | 1236 |
| Audrey Caling | shARF6 | shARF6 | Service_Clair | Collagen alph | CO1A1_HUM | Collagen alph | 100.00% | 1.07% | 138,923.40 | 18 | 23 | 25 | index=8954 | ADDANVVRD | R | S | 99.60% | | 2,144.08 | 3 | 0.0051 | 2.4 | 1218 | 1236 |
| Audrey Caling | shARF6 | shARF6 | Service_Clair | Collagen alph | CO1A1_HUM | Collagen alph | 100.00% | 1.07% | 138,923.40 | 18 | 23 | 25 | index=8953 | ADDANVVRD | R | S | 99.70% | | 2,144.08 | 3 | 0.0051 | 2.4 | 1218 | 1236 |
| Audrey Caling | shARF6 | shARF6 | Service_Clair | Collagen alph | CO1A1_HUM | Collagen alph | 100.00% | 1.07% | 138,923.40 | 18 | 23 | 25 | index=2129 | GESGPSGPA | K | G | 99.70% | | 1,296.61 | 2 | 0.0022 | 1.7 | 782 | 796 |
| Audrey Caling | shARF6 | shARF6 | Service_Clair | Collagen alph | CO1A1_HUM | Collagen alph | 100.00% | 1.07% | 138,923.40 | 18 | 23 | 25 | index=6834 | GFPGERGVQ | R | G | 95.90% | | 1,731.89 | 3 | 0.0055 | 3.2 | 680 | 697 |
| Audrey Caling | shARF6 | shARF6 | Service_Clair | Collagen alph | CO1A1_HUM | Collagen alph | 100.00% | 1.07% | 138,923.40 | 18 | 23 | 25 | index=6833 | GFPGERGVQ | R | G | 99.70% | | 1,731.89 | 3 | 0.0055 | 3.2 | 680 | 697 |
| Audrey Caling | shARF6 | shARF6 | Service_Clair | Collagen alph | CO1A1_HUM | Collagen alph | 100.00% | 1.07% | 138,923.40 | 18 | 23 | 25 | index=3547 | GLPGERGRP | R | G | 99.70% | | 1,671.90 | 4 | 0.004 | 2.4 | 305 | 322 |
| Audrey Caling | shARF6 | shARF6 | Service_Clair | Collagen alph | CO1A1_HUM | Collagen alph | 100.00% | 1.07% | 138,923.40 | 18 | 23 | 25 | index=3295 | GLTGSPGSP | K | T | 99.70% | | 1,225.60 | 2 | 0.0017 | 1.4 | 539 | 552 |
| Audrey Caling | shARF6 | shARF6 | Service_Clair | Collagen alph | CO1A1_HUM | Collagen alph | 100.00% | 1.07% | 138,923.40 | 18 | 23 | 25 | index=1248 | GPAGPQGPR | R | G | 96.00% | | 835.43 | 2 | 0.0012 | 1.5 | 1085 | 1093 |
| Audrey Caling | shARF6 | shARF6 | Service_Clair | Collagen alph | CO1A1_HUM | Collagen alph | 100.00% | 1.07% | 138,923.40 | 18 | 23 | 25 | index=1035 | GPAGPQGPR | R | G | 99.70% | | 2,064.96 | 4 | 0.0066 | 3.2 | 1085 | 1105 |
| Audrey Caling | shARF6 | shARF6 | Service_Clair | Collagen alph | CO1A1_HUM | Collagen alph | 100.00% | 1.07% | 138,923.40 | 18 | 23 | 25 | index=2663 | GPSGPQGPG | R | G | 99.70% | | 1,285.65 | 2 | 0.0046 | 3.6 | 416 | 430 |
| Audrey Caling | shARF6 | shARF6 | Service_Clair | Collagen alph | CO1A1_HUM | Collagen alph | 100.00% | 1.07% | 138,923.40 | 18 | 23 | 25 | index=1153 | GRPGAPGPA | R | G | 99.70% | | 1,062.57 | 3 | 0.0004 | 0.38 | 311 | 322 |
| Audrey Caling | shARF6 | shARF6 | Service_Clair | Collagen alph | CO1A1_HUM | Collagen alph | 100.00% | 1.07% | 138,923.40 | 18 | 23 | 25 | index=15654 | GRTGDAGPV | R | Y | 99.70% | Phospho (+80 | 4,533.15 | 5 | 0.02 | 4.5 | 1169 | 1214 |
| Audrey Caling | shARF6 | shARF6 | Service_Clair | Collagen alph | CO1A1_HUM | Collagen alph | 100.00% | 1.07% | 138,923.40 | 18 | 23 | 25 | index=15689 | GRTGDAGPV | R | Y | 99.70% | Phospho (+80 | 4,533.15 | 4 | 0.02 | 4.5 | 1169 | 1214 |
| Audrey Caling | shARF6 | shARF6 | Service_Clair | Collagen alph | CO1A1_HUM | Collagen alph | 100.00% | 1.07% | 138,923.40 | 18 | 23 | 25 | index=7012 | GSAGPPGAT | R | V | 99.70% | | 1,426.70 | 2 | 0.0037 | 2.6 | 866 | 882 |
| Audrey Caling | shARF6 | shARF6 | Service_Clair | Collagen alph | CO1A1_HUM | Collagen alph | 100.00% | 1.07% | 138,923.40 | 18 | 23 | 25 | index=3234 | GVQGPPGPA | R | G | 99.70% | | 1,088.57 | 2 | 0.0016 | 1.5 | 686 | 697 |
| Audrey Caling | shARF6 | shARF6 | Service_Clair | Collagen alph | CO1A1_HUM | Collagen alph | 100.00% | 1.07% | 138,923.40 | 18 | 23 | 25 | index=5821 | GVVGLPGQR | R | G | 96.80% | | 1,223.68 | 3 | 0.0017 | 1.3 | 959 | 970 |
| Audrey Caling | shARF6 | shARF6 | Service_Clair | Collagen alph | CO1A1_HUM | Collagen alph | 100.00% | 1.07% | 138,923.40 | 18 | 23 | 25 | index=5820 | GVVGLPGQR | R | G | 99.70% | | 1,223.68 | 2 | 0.0034 | 2.8 | 959 | 970 |
| Audrey Caling | shARF6 | shARF6 | Service_Clair | Collagen alph | CO1A1_HUM | Collagen alph | 100.00% | 1.07% | 138,923.40 | 18 | 23 | 25 | index=9586 | SLSQQIENIR | K | K | 99.70% | | 1,799.92 | 2 | 0.0047 | 2.6 | 1237 | 1252 |
| Audrey Caling | shARF6 | shARF6 | Service_Clair | Collagen alph | CO1A1_HUM | Collagen alph | 100.00% | 1.07% | 138,923.40 | 18 | 23 | 25 | index=9563 | SLSQQIENIR | K | K | 99.70% | | 1,799.91 | 3 | 0.0018 | 0.99 | 1237 | 1252 |
| Audrey Caling | shARF6 | shARF6 | Service_Clair | Collagen alph | CO1A1_HUM | Collagen alph | 100.00% | 1.07% | 138,923.40 | 18 | 23 | 25 | index=16111 | TGDAGPVGPI | R | Y | 99.70% | Phospho (+80 | 4,320.03 | 4 | 0.018 | 4.2 | 1171 | 1214 |
| Audrey Caling | shARF6 | shARF6 | Service_Clair | Collagen alph | CO1A1_HUM | Collagen alph | 100.00% | 1.07% | 138,923.40 | 18 | 23 | 25 | index=3608 | TVIEYKTTK | K | T | 99.70% | | 1,081.60 | 2 | 0.0028 | 2.6 | 1425 | 1433 |
| Audrey Caling | shARF6 | shARF6 | Service_Clair | Collagen alph | CO1A1_HUM | Collagen alph | 100.00% | 1.07% | 138,923.40 | 18 | 23 | 25 | index=3621 | TVIEYKTTK | K | T | 99.70% | | 1,081.60 | 3 | 0.0023 | 2.1 | 1425 | 1433 |
| Audrey Caling | shARF6 | shARF6 | Service_Clair | Collagen alph | CO1A1_HUM | Collagen alph | 100.00% | 1.07% | 138,923.40 | 18 | 23 | 25 | index=4281 | YYRADDANV | R | D | 99.60% | | 1,340.65 | 3 | 0.0013 | 0.99 | 1215 | 1225 |
| Audrey Caling | shARF6 | shARF6 | Service_Clair | Collagen alph | CO1A1_HUM | Collagen alph | 100.00% | 1.07% | 138,923.40 | 18 | 23 | 25 | index=9281 | YYRADDANV | R | S | 99.70% | | 2,626.31 | 4 | 0.0069 | 2.6 | 1215 | 1236 |
| Audrey Caling | shARF6 | shARF6 | Service_Clair | Collagen alph | CO1A1_HUM | Collagen alph | 100.00% | 1.07% | 138,923.40 | 18 | 23 | 25 | index=9278 | YYRADDANV | R | S | 99.70% | | 2,626.31 | 3 | 0.0075 | 2.8 | 1215 | 1236 |

| Experiment na | Biological sar | Biological sar | MS/MS sampl | Protein group | Protein acces | Protein name | Protein identi | Protein perce | Protein molec | Number of un | Number of un | Number of tot | Spectrum nar | Peptide seque | Previous amir | Next amino ac | Peptide identi | Modifications | Actual peptide | Spectrum cha | Actual minus | Actual minus | Peptide start i | Peptide stop index |
|---|---|---|---|---|---|---|---|---|---|---|---|---|---|---|---|---|---|---|---|---|---|---|---|---|
| Audrey Caling | SCR | SCR | Service_Clair | Collagen alph | CO1A2_HUM | Collagen alph | 100.00% | 5.86% | 129,296.60 | 72 | 120 | 160 | index=2677 | AGEPGEPGQ | P | G | 99.70% | | 1,450.69 | 2 | 0.0063 | 4.3 | 117 | 132 |
| Audrey Caling | SCR | SCR | Service_Clair | Collagen alph | CO1A2_HUM | Collagen alph | 100.00% | 5.86% | 129,296.60 | 72 | 120 | 160 | index=6449 | AGPNGFAGP | P | G | 99.70% | Deamidated ( | 1,937.94 | 3 | 0.0066 | 3.4 | 720 | 741 |
| Audrey Caling | SCR | SCR | Service_Clair | Collagen alph | CO1A2_HUM | Collagen alph | 100.00% | 5.86% | 129,296.60 | 72 | 120 | 160 | index=5812 | AGPNGFAGP | P | G | 99.70% | Deamidated ( | 1,937.94 | 3 | 0.0066 | 3.4 | 720 | 741 |
| Audrey Caling | SCR | SCR | Service_Clair | Collagen alph | CO1A2_HUM | Collagen alph | 100.00% | 5.86% | 129,296.60 | 72 | 120 | 160 | index=2096 | AGVMGPPGS | R | G | 99.00% | Oxidation (+1 | 943.46 | 2 | 0.0022 | 2.3 | 414 | 423 |
| Audrey Caling | SCR | SCR | Service_Clair | Collagen alph | CO1A2_HUM | Collagen alph | 100.00% | 5.86% | 129,296.60 | 72 | 120 | 160 | index=4337 | AGVMGPPGS | R | G | 99.70% | | 927.46 | 2 | 0.0015 | 1.6 | 414 | 423 |
| Audrey Caling | SCR | SCR | Service_Clair | Collagen alph | CO1A2_HUM | Collagen alph | 100.00% | 5.86% | 129,296.60 | 72 | 120 | 160 | index=4336 | AGVMGPPGS | R | G | 99.70% | | 927.46 | 2 | 0.0015 | 1.6 | 414 | 423 |
| Audrey Caling | SCR | SCR | Service_Clair | Collagen alph | CO1A2_HUM | Collagen alph | 100.00% | 5.86% | 129,296.60 | 72 | 120 | 160 | index=2095 | AGVMGPPGS | R | G | 99.70% | Oxidation (+1 | 943.46 | 2 | 0.0022 | 2.3 | 414 | 423 |
| Audrey Caling | SCR | SCR | Service_Clair | Collagen alph | CO1A2_HUM | Collagen alph | 100.00% | 5.86% | 129,296.60 | 72 | 120 | 160 | index=8028 | AGVMGPPGS | R | G | 97.30% | Deamidated ( | 3,412.66 | 4 | 0.011 | 3.1 | 414 | 450 |
| Audrey Caling | SCR | SCR | Service_Clair | Collagen alph | CO1A2_HUM | Collagen alph | 100.00% | 5.86% | 129,296.60 | 72 | 120 | 160 | index=7019 | AGVMGPPGS | R | G | 97.90% | Oxidation (+1 | 3,428.65 | 5 | 0.0082 | 2.4 | 414 | 450 |
| Audrey Caling | SCR | SCR | Service_Clair | Collagen alph | CO1A2_HUM | Collagen alph | 100.00% | 5.86% | 129,296.60 | 72 | 120 | 160 | index=8046 | AGVMGPPGS | R | G | 99.00% | Deamidated ( | 3,412.66 | 5 | 0.012 | 3.6 | 414 | 450 |
| Audrey Caling | SCR | SCR | Service_Clair | Collagen alph | CO1A2_HUM | Collagen alph | 100.00% | 5.86% | 129,296.60 | 72 | 120 | 160 | index=9459 | AGVMGPPGS | R | G | 99.70% | Deamidated ( | 3,396.67 | 5 | 0.016 | 4.8 | 414 | 450 |
| Audrey Caling | SCR | SCR | Service_Clair | Collagen alph | CO1A2_HUM | Collagen alph | 100.00% | 5.86% | 129,296.60 | 72 | 120 | 160 | index=9432 | AGVMGPPGS | R | G | 99.70% | Deamidated ( | 3,396.67 | 4 | 0.015 | 4.3 | 414 | 450 |
| Audrey Caling | SCR | SCR | Service_Clair | Collagen alph | CO1A2_HUM | Collagen alph | 100.00% | 5.86% | 129,296.60 | 72 | 120 | 160 | index=9130 | AQPENIPAKN | R | S | 99.70% | | 1,585.81 | 2 | 0.0075 | 4.8 | 1206 | 1218 |
| Audrey Caling | SCR | SCR | Service_Clair | Collagen alph | CO1A2_HUM | Collagen alph | 100.00% | 5.86% | 129,296.60 | 72 | 120 | 160 | index=9104 | AQPENIPAKN | R | S | 99.70% | | 1,585.81 | 3 | 0.0076 | 4.8 | 1206 | 1218 |
| Audrey Caling | SCR | SCR | Service_Clair | Collagen alph | CO1A2_HUM | Collagen alph | 100.00% | 5.86% | 129,296.60 | 72 | 120 | 160 | index=9105 | AQPENIPAKN | R | S | 99.70% | | 1,585.81 | 3 | 0.0076 | 4.8 | 1206 | 1218 |
| Audrey Caling | SCR | SCR | Service_Clair | Collagen alph | CO1A2_HUM | Collagen alph | 100.00% | 5.86% | 129,296.60 | 72 | 120 | 160 | index=9131 | AQPENIPAKN | R | S | 99.70% | | 1,585.81 | 2 | 0.0075 | 4.8 | 1206 | 1218 |
| Audrey Caling | SCR | SCR | Service_Clair | Collagen alph | CO1A2_HUM | Collagen alph | 100.00% | 5.86% | 129,296.60 | 72 | 120 | 160 | index=5933 | DGARGAPGA | R | G | 99.70% | | 3,093.52 | 4 | 0.0093 | 3 | 672 | 708 |
| Audrey Caling | SCR | SCR | Service_Clair | Collagen alph | CO1A2_HUM | Collagen alph | 100.00% | 5.86% | 129,296.60 | 72 | 120 | 160 | index=20444 | DYEVDATLKS | K | K | 99.70% | | 2,805.40 | 3 | 0.0009 | 0.32 | 1132 | 1156 |
| Audrey Caling | SCR | SCR | Service_Clair | Collagen alph | CO1A2_HUM | Collagen alph | 100.00% | 5.86% | 129,296.60 | 72 | 120 | 160 | index=3570 | EGLRGPRGD | K | T | 98.70% | | 1,549.81 | 3 | 0.0049 | 3.1 | 816 | 830 |
| Audrey Caling | SCR | SCR | Service_Clair | Collagen alph | CO1A2_HUM | Collagen alph | 100.00% | 5.86% | 129,296.60 | 72 | 120 | 160 | index=3569 | EGLRGPRGD | K | T | 99.70% | | 1,549.81 | 3 | 0.0049 | 3.1 | 816 | 830 |
| Audrey Caling | SCR | SCR | Service_Clair | Collagen alph | CO1A2_HUM | Collagen alph | 100.00% | 5.86% | 129,296.60 | 72 | 120 | 160 | index=12684 | EMATQLAFMF | K | L | 99.70% | Oxidation (+1 | 1,212.57 | 2 | 0.0045 | 3.7 | 1249 | 1258 |
| Audrey Caling | SCR | SCR | Service_Clair | Collagen alph | CO1A2_HUM | Collagen alph | 100.00% | 5.86% | 129,296.60 | 72 | 120 | 160 | index=11032 | EMATQLAFMF | K | L | 99.70% | Oxidation (+1 | 1,212.56 | 2 | 0.0014 | 1.1 | 1249 | 1258 |
| Audrey Caling | SCR | SCR | Service_Clair | Collagen alph | CO1A2_HUM | Collagen alph | 100.00% | 5.86% | 129,296.60 | 72 | 120 | 160 | index=14818 | EMATQLAFMF | K | L | 99.70% | | 1,196.57 | 2 | 0.0038 | 3.2 | 1249 | 1258 |
| Audrey Caling | SCR | SCR | Service_Clair | Collagen alph | CO1A2_HUM | Collagen alph | 100.00% | 5.86% | 129,296.60 | 72 | 120 | 160 | index=12755 | GAAGLPGVA( | K | G | 99.70% | | 1,513.84 | 2 | 0.0047 | 3.1 | 310 | 327 |
| Audrey Caling | SCR | SCR | Service_Clair | Collagen alph | CO1A2_HUM | Collagen alph | 100.00% | 5.86% | 129,296.60 | 72 | 120 | 160 | index=14639 | GAAGLPGVA( | K | G | 99.70% | | 2,746.51 | 3 | 0.0074 | 2.7 | 310 | 342 |
| Audrey Caling | SCR | SCR | Service_Clair | Collagen alph | CO1A2_HUM | Collagen alph | 100.00% | 5.86% | 129,296.60 | 72 | 120 | 160 | index=9845 | GAKGPKGEN | R | T | 99.70% | Deamidated ( | 4,589.25 | 5 | 0.038 | 8.3 | 742 | 794 |
| Audrey Caling | SCR | SCR | Service_Clair | Collagen alph | CO1A2_HUM | Collagen alph | 100.00% | 5.86% | 129,296.60 | 72 | 120 | 160 | index=9844 | GAKGPKGEN | R | T | 99.70% | Deamidated ( | 4,589.25 | 5 | 0.038 | 8.3 | 742 | 794 |
| Audrey Caling | SCR | SCR | Service_Clair | Collagen alph | CO1A2_HUM | Collagen alph | 100.00% | 5.86% | 129,296.60 | 72 | 120 | 160 | index=11529 | GAKGPKGEN | R | T | 99.70% | Deamidated ( | 4,572.25 | 5 | 0.021 | 4.7 | 742 | 794 |
| Audrey Caling | SCR | SCR | Service_Clair | Collagen alph | CO1A2_HUM | Collagen alph | 100.00% | 5.86% | 129,296.60 | 72 | 120 | 160 | index=11669 | GAKGPKGEN | R | T | 99.70% | Deamidated ( | 4,573.23 | 5 | 0.015 | 3.3 | 742 | 794 |
| Audrey Caling | SCR | SCR | Service_Clair | Collagen alph | CO1A2_HUM | Collagen alph | 100.00% | 5.86% | 129,296.60 | 72 | 120 | 160 | index=9677 | GAKGPKGEN | R | T | 99.70% | Deamidated ( | 4,588.25 | 5 | 0.027 | 5.8 | 742 | 794 |
| Audrey Caling | SCR | SCR | Service_Clair | Collagen alph | CO1A2_HUM | Collagen alph | 100.00% | 5.86% | 129,296.60 | 72 | 120 | 160 | index=6147 | GAPGAVGAP( | R | A | 99.70% | | 2,088.01 | 2 | 0.0076 | 3.6 | 676 | 701 |
| Audrey Caling | SCR | SCR | Service_Clair | Collagen alph | CO1A2_HUM | Collagen alph | 100.00% | 5.86% | 129,296.60 | 72 | 120 | 160 | index=6173 | GAPGAVGAP( | R | G | 99.70% | | 2,694.33 | 3 | 0.0066 | 2.5 | 676 | 708 |
| Audrey Caling | SCR | SCR | Service_Clair | Collagen alph | CO1A2_HUM | Collagen alph | 100.00% | 5.86% | 129,296.60 | 72 | 120 | 160 | index=6172 | GAPGAVGAP( | R | G | 99.70% | | 2,694.33 | 3 | 0.0066 | 2.5 | 676 | 708 |
| Audrey Caling | SCR | SCR | Service_Clair | Collagen alph | CO1A2_HUM | Collagen alph | 100.00% | 5.86% | 129,296.60 | 72 | 120 | 160 | index=6171 | GAPGAVGAP( | R | G | 99.70% | | 2,694.33 | 4 | 0.0045 | 1.7 | 676 | 708 |
| Audrey Caling | SCR | SCR | Service_Clair | Collagen alph | CO1A2_HUM | Collagen alph | 100.00% | 5.86% | 129,296.60 | 72 | 120 | 160 | index=7221 | GAPGAVGAP( | R | A | 99.70% | | 3,716.82 | 4 | 0.015 | 4.1 | 676 | 719 |
| Audrey Caling | SCR | SCR | Service_Clair | Collagen alph | CO1A2_HUM | Collagen alph | 100.00% | 5.86% | 129,296.60 | 72 | 120 | 160 | index=1760 | GASGPAGVR | R | G | 98.60% | | 770.41 | 2 | 0.0018 | 2.3 | 424 | 432 |
| Audrey Caling | SCR | SCR | Service_Clair | Collagen alph | CO1A2_HUM | Collagen alph | 100.00% | 5.86% | 129,296.60 | 72 | 120 | 160 | index=8135 | GASGPAGVR( | R | G | 96.60% | Deamidated ( | 2,487.21 | 4 | 0.0025 | 1 | 424 | 450 |
| Audrey Caling | SCR | SCR | Service_Clair | Collagen alph | CO1A2_HUM | Collagen alph | 100.00% | 5.86% | 129,296.60 | 72 | 120 | 160 | index=6541 | GASGPAGVR( | R | G | 99.70% | Deamidated ( | 2,503.21 | 3 | 0.0092 | 3.7 | 424 | 450 |
| Audrey Caling | SCR | SCR | Service_Clair | Collagen alph | CO1A2_HUM | Collagen alph | 100.00% | 5.86% | 129,296.60 | 72 | 120 | 160 | index=7722 | GASGPAGVR( | R | G | 99.70% | Deamidated ( | 2,503.20 | 3 | 0.0058 | 2.3 | 424 | 450 |
| Audrey Caling | SCR | SCR | Service_Clair | Collagen alph | CO1A2_HUM | Collagen alph | 100.00% | 5.86% | 129,296.60 | 72 | 120 | 160 | index=7503 | GASGPAGVR( | R | G | 99.70% | Deamidated ( | 2,503.21 | 4 | 0.0066 | 2.6 | 424 | 450 |
| Audrey Caling | SCR | SCR | Service_Clair | Collagen alph | CO1A2_HUM | Collagen alph | 100.00% | 5.86% | 129,296.60 | 72 | 120 | 160 | index=6265 | GASGPAGVR( | R | G | 99.70% | Oxidation (+1 | 2,502.22 | 4 | 0.0082 | 3.3 | 424 | 450 |
| Audrey Caling | SCR | SCR | Service_Clair | Collagen alph | CO1A2_HUM | Collagen alph | 100.00% | 5.86% | 129,296.60 | 72 | 120 | 160 | index=7507 | GASGPAGVR( | R | G | 99.70% | Deamidated ( | 2,503.21 | 3 | 0.0086 | 3.4 | 424 | 450 |
| Audrey Caling | SCR | SCR | Service_Clair | Collagen alph | CO1A2_HUM | Collagen alph | 100.00% | 5.86% | 129,296.60 | 72 | 120 | 160 | index=6540 | GASGPAGVR( | R | G | 99.70% | Deamidated ( | 2,503.21 | 3 | 0.0092 | 3.7 | 424 | 450 |
| Audrey Caling | SCR | SCR | Service_Clair | Collagen alph | CO1A2_HUM | Collagen alph | 100.00% | 5.86% | 129,296.60 | 72 | 120 | 160 | index=8140 | GASGPAGVR( | R | G | 99.70% | Deamidated ( | 2,487.21 | 3 | 0.0044 | 1.8 | 424 | 450 |
| Audrey Caling | SCR | SCR | Service_Clair | Collagen alph | CO1A2_HUM | Collagen alph | 100.00% | 5.86% | 129,296.60 | 72 | 120 | 160 | index=10605 | GDGGPPGMT | R | T | 99.70% | | 1,500.68 | 2 | 0.0029 | 1.9 | 778 | 794 |
| Audrey Caling | SCR | SCR | Service_Clair | Collagen alph | CO1A2_HUM | Collagen alph | 100.00% | 5.86% | 129,296.60 | 72 | 120 | 160 | index=1942 | GEIGNPGRD( | R | G | 95.10% | | 1,197.59 | 3 | 0.0028 | 2.3 | 664 | 675 |
| Audrey Caling | SCR | SCR | Service_Clair | Collagen alph | CO1A2_HUM | Collagen alph | 100.00% | 5.86% | 129,296.60 | 72 | 120 | 160 | index=2863 | GEPGAPGEN( | K | G | 99.70% | Deamidated ( | 1,652.74 | 2 | 0.0051 | 3.1 | 199 | 216 |
| Audrey Caling | SCR | SCR | Service_Clair | Collagen alph | CO1A2_HUM | Collagen alph | 100.00% | 5.86% | 129,296.60 | 72 | 120 | 160 | index=6015 | GEPGLRGEIG | K | G | 99.00% | | 1,806.92 | 3 | 0.0075 | 4.2 | 658 | 675 |
| Audrey Caling | SCR | SCR | Service_Clair | Collagen alph | CO1A2_HUM | Collagen alph | 100.00% | 5.86% | 129,296.60 | 72 | 120 | 160 | index=10048 | GEPGNIGFPG | R | G | 99.70% | | 1,168.59 | 2 | 0.0032 | 2.7 | 487 | 498 |
| Audrey Caling | SCR | SCR | Service_Clair | Collagen alph | CO1A2_HUM | Collagen alph | 100.00% | 5.86% | 129,296.60 | 72 | 120 | 160 | index=10047 | GEPGNIGFPG | R | G | 99.70% | | 1,168.59 | 2 | 0.0032 | 2.7 | 487 | 498 |
| Audrey Caling | SCR | SCR | Service_Clair | Collagen alph | CO1A2_HUM | Collagen alph | 100.00% | 5.86% | 129,296.60 | 72 | 120 | 160 | index=8500 | GEPGNIGFPG | R | G | 99.70% | Deamidated ( | 3,083.53 | 4 | 0.0076 | 2.5 | 487 | 519 |
| Audrey Caling | SCR | SCR | Service_Clair | Collagen alph | CO1A2_HUM | Collagen alph | 100.00% | 5.86% | 129,296.60 | 72 | 120 | 160 | index=8486 | GEPGNIGFPG | R | G | 99.70% | Deamidated ( | 3,083.52 | 5 | 0.0057 | 1.9 | 487 | 519 |
| Audrey Caling | SCR | SCR | Service_Clair | Collagen alph | CO1A2_HUM | Collagen alph | 100.00% | 5.86% | 129,296.60 | 72 | 120 | 160 | index=5296 | GERGPPGES( | R | G | 99.70% | | 1,905.94 | 3 | 0.0069 | 3.6 | 589 | 609 |
| Audrey Caling | SCR | SCR | Service_Clair | Collagen alph | CO1A2_HUM | Collagen alph | 100.00% | 5.86% | 129,296.60 | 72 | 120 | 160 | index=9014 | GERGYPGNIG | K | H | 99.70% | | 2,593.33 | 4 | 0.012 | 4.5 | 946 | 974 |
| Audrey Caling | SCR | SCR | Service_Clair | Collagen alph | CO1A2_HUM | Collagen alph | 100.00% | 5.86% | 129,296.60 | 72 | 120 | 160 | index=7347 | GIPGPVGAAG | R | G | 99.70% | | 1,250.68 | 2 | 0.0025 | 2 | 328 | 342 |
| Audrey Caling | SCR | SCR | Service_Clair | Collagen alph | CO1A2_HUM | Collagen alph | 100.00% | 5.86% | 129,296.60 | 72 | 120 | 160 | index=7336 | GIPGPVGAAG | R | G | 99.70% | | 1,250.68 | 3 | 0.0024 | 1.9 | 328 | 342 |
| Audrey Caling | SCR | SCR | Service_Clair | Collagen alph | CO1A2_HUM | Collagen alph | 100.00% | 5.86% | 129,296.60 | 72 | 120 | 160 | index=7346 | GIPGPVGAAG | R | G | 99.70% | | 1,250.68 | 2 | 0.0025 | 2 | 328 | 342 |
| Audrey Caling | SCR | SCR | Service_Clair | Collagen alph | CO1A2_HUM | Collagen alph | 100.00% | 5.86% | 129,296.60 | 72 | 120 | 160 | index=11823 | GLHGEFGLP( | R | G | 99.70% | | 1,460.76 | 3 | 0.0058 | 4 | 574 | 588 |
| Audrey Caling | SCR | SCR | Service_Clair | Collagen alph | CO1A2_HUM | Collagen alph | 100.00% | 5.86% | 129,296.60 | 72 | 120 | 160 | index=10397 | GLHGEFGLP( | R | G | 99.70% | | 1,802.92 | 4 | 0.0031 | 1.7 | 574 | 591 |
| Audrey Caling | SCR | SCR | Service_Clair | Collagen alph | CO1A2_HUM | Collagen alph | 100.00% | 5.86% | 129,296.60 | 72 | 120 | 160 | index=7930 | GLPGADGRA | R | G | 99.10% | | 1,650.83 | 3 | 0.0044 | 2.7 | 406 | 423 |
| Audrey Caling | SCR | SCR | Service_Clair | Collagen alph | CO1A2_HUM | Collagen alph | 100.00% | 5.86% | 129,296.60 | 72 | 120 | 160 | index=7931 | GLPGADGRA | R | G | 99.70% | | 1,650.83 | 3 | 0.0044 | 2.7 | 406 | 423 |
| Audrey Caling | SCR | SCR | Service_Clair | Collagen alph | CO1A2_HUM | Collagen alph | 100.00% | 5.86% | 129,296.60 | 72 | 120 | 160 | index=7108 | GLPGADGRA | R | G | 99.70% | Oxidation (+1 | 1,666.83 | 2 | 0.0066 | 3.9 | 406 | 423 |
| Audrey Caling | SCR | SCR | Service_Clair | Collagen alph | CO1A2_HUM | Collagen alph | 100.00% | 5.86% | 129,296.60 | 72 | 120 | 160 | index=7101 | GLPGADGRA | R | G | 99.70% | Oxidation (+1 | 1,666.83 | 3 | 0.0044 | 2.6 | 406 | 423 |
| Audrey Caling | SCR | SCR | Service_Clair | Collagen alph | CO1A2_HUM | Collagen alph | 100.00% | 5.86% | 129,296.60 | 72 | 120 | 160 | index=7927 | GLPGADGRA | R | G | 99.70% | | 1,650.83 | 2 | 0.0055 | 3.3 | 406 | 423 |
| Audrey Caling | SCR | SCR | Service_Clair | Collagen alph | CO1A2_HUM | Collagen alph | 100.00% | 5.86% | 129,296.60 | 72 | 120 | 160 | index=9614 | GLPGADGRA | R | G | 97.50% | Oxidation (+1 | 4,152.03 | 5 | 0.019 | 4.5 | 406 | 450 |
| Audrey Caling | SCR | SCR | Service_Clair | Collagen alph | CO1A2_HUM | Collagen alph | 100.00% | 5.86% | 129,296.60 | 72 | 120 | 160 | index=10215 | GLPGADGRA | R | G | 99.70% | Deamidated ( | 4,120.04 | 6 | 0.02 | 4.8 | 406 | 450 |
| Audrey Caling | SCR | SCR | Service_Clair | Collagen alph | CO1A2_HUM | Collagen alph | 100.00% | 5.86% | 129,296.60 | 72 | 120 | 160 | index=4656 | GLPGERGRV( | R | G | 99.70% | | 1,673.91 | 4 | 0.004 | 2.4 | 217 | 234 |
| Audrey Caling | SCR | SCR | Service_Clair | Collagen alph | CO1A2_HUM | Collagen alph | 100.00% | 5.86% | 129,296.60 | 72 | 120 | 160 | index=4655 | GLPGERGRV( | R | G | 99.70% | | 1,673.91 | 4 | 0.004 | 2.4 | 217 | 234 |
| Audrey Caling | SCR | SCR | Service_Clair | Collagen alph | CO1A2_HUM | Collagen alph | 100.00% | 5.86% | 129,296.60 | 72 | 120 | 160 | index=4662 | GLPGERGRV( | R | G | 99.70% | | 1,673.91 | 3 | 0.0036 | 2.2 | 217 | 234 |
| Audrey Caling | SCR | SCR | Service_Clair | Collagen alph | CO1A2_HUM | Collagen alph | 100.00% | 5.86% | 129,296.60 | 72 | 120 | 160 | index=12459 | GLPGLKGHN | R | G | 98.60% | Deamidated ( | 3,333.72 | 5 | 0.012 | 3.7 | 1018 | 1053 |
| Audrey Caling | SCR | SCR | Service_Clair | Collagen alph | CO1A2_HUM | Collagen alph | 100.00% | 5.86% | 129,296.60 | 72 | 120 | 160 | index=12118 | GLPGLKGHN | R | G | 99.70% | Deamidated ( | 3,333.72 | 5 | 0.0092 | 2.8 | 1018 | 1053 |
| Audrey Caling | SCR | SCR | Service_Clair | Collagen alph | CO1A2_HUM | Collagen alph | 100.00% | 5.86% | 129,296.60 | 72 | 120 | 160 | index=7678 | GLPGSPGNIG | R | E | 99.70% | | 1,220.66 | 2 | 0.0042 | 3.4 | 451 | 464 |
| Audrey Caling | SCR | SCR | Service_Clair | Collagen alph | CO1A2_HUM | Collagen alph | 100.00% | 5.86% | 129,296.60 | 72 | 120 | 160 | index=7677 | GLPGSPGNIG | R | E | 99.70% | | 1,220.66 | 2 | 0.0042 | 3.4 | 451 | 464 |
| Audrey Caling | SCR | SCR | Service_Clair | Collagen alph | CO1A2_HUM | Collagen alph | 100.00% | 5.86% | 129,296.60 | 72 | 120 | 160 | index=16014 | GLPGVAGAV( | R | G | 99.70% | | 2,066.14 | 2 | 0.0089 | 4.3 | 883 | 906 |
| Audrey Caling | SCR | SCR | Service_Clair | Collagen alph | CO1A2_HUM | Collagen alph | 100.00% | 5.86% | 129,296.60 | 72 | 120 | 160 | index=16009 | GLPGVAGAV( | R | G | 99.70% | | 2,066.14 | 3 | 0.0077 | 3.7 | 883 | 906 |
| Audrey Caling | SCR | SCR | Service_Clair | Collagen alph | CO1A2_HUM | Collagen alph | 100.00% | 5.86% | 129,296.60 | 72 | 120 | 160 | index=16008 | GLPGVAGAV( | R | G | 99.70% | | 2,066.14 | 3 | 0.0077 | 3.7 | 883 | 906 |
| Audrey Caling | SCR | SCR | Service_Clair | Collagen alph | CO1A2_HUM | Collagen alph | 100.00% | 5.86% | 129,296.60 | 72 | 120 | 160 | index=16013 | GLPGVAGAV( | R | G | 99.70% | | 2,066.14 | 2 | 0.0089 | 4.3 | 883 | 906 |
| Audrey Caling | SCR | SCR | Service_Clair | Collagen alph | CO1A2_HUM | Collagen alph | 100.00% | 5.86% | 129,296.60 | 72 | 120 | 160 | index=1257 | GPAGPPGKA( | R | G | 98.10% | | 2,277.14 | 4 | 0.0076 | 3.3 | 133 | 156 |
| Audrey Caling | SCR | SCR | Service_Clair | Collagen alph | CO1A2_HUM | Collagen alph | 100.00% | 5.86% | 129,296.60 | 72 | 120 | 160 | index=1253 | GPAGPPGKA( | R | G | 99.70% | | 2,277.15 | 3 | 0.0096 | 4.2 | 133 | 156 |
| Audrey Caling | SCR | SCR | Service_Clair | Collagen alph | CO1A2_HUM | Collagen alph | 100.00% | 5.86% | 129,296.60 | 72 | 120 | 160 | index=1256 | GPAGPPGKA( | R | G | 99.70% | | 2,277.14 | 4 | 0.0076 | 3.3 | 133 | 156 |
| Audrey Caling | SCR | SCR | Service_Clair | Collagen alph | CO1A2_HUM | Collagen alph | 100.00% | 5.86% | 129,296.60 | 72 | 120 | 160 | index=2622 | GPAGPPGKA( | R | G | 99.50% | | 3,098.59 | 7 | 0.0065 | 2.1 | 133 | 165 |
| Audrey Caling | SCR | SCR | Service_Clair | Collagen alph | CO1A2_HUM | Collagen alph | 100.00% | 5.86% | 129,296.60 | 72 | 120 | 160 | index=2629 | GPAGPPGKA( | R | G | 99.70% | | 3,098.60 | 4 | 0.011 | 3.6 | 133 | 165 |
| Audrey Caling | SCR | SCR | Service_Clair | Collagen alph | CO1A2_HUM | Collagen alph | 100.00% | 5.86% | 129,296.60 | 72 | 120 | 160 | index=1314 | GPAGPSGPA( | R | D | 99.70% | | 894.46 | 2 | 0.0028 | 3.1 | 1054 | 1064 |
| Audrey Caling | SCR | SCR | Service_Clair | Collagen alph | CO1A2_HUM | Collagen alph | 100.00% | 5.86% | 129,296.60 | 72 | 120 | 160 | index=1313 | GPAGPSGPA( | R | D | 99.70% | | 894.46 | 2 | 0.0028 | 3.1 | 1054 | 1064 |
| Audrey Caling | SCR | SCR | Service_Clair | Collagen alph | CO1A2_HUM | Collagen alph | 100.00% | 5.86% | 129,296.60 | 72 | 120 | 160 | index=1148 | GPAGPSGPA( | R | T | 99.70% | | 1,222.61 | 3 | 0.0022 | 1.8 | 1054 | 1067 |
| Audrey Caling | SCR | SCR | Service_Clair | Collagen alph | CO1A2_HUM | Collagen alph | 100.00% | 5.86% | 129,296.60 | 72 | 120 | 160 | index=1147 | GPAGPSGPA( | R | T | 99.70% | | 1,222.61 | 3 | 0.0022 | 1.8 | 1054 | 1067 |
| Audrey Caling | SCR | SCR | Service_Clair | Collagen alph | CO1A2_HUM | Collagen alph | 100.00% | 5.86% | 129,296.60 | 72 | 120 | 160 | index=1154 | GPAGPSGPA( | R | T | 99.70% | | 1,222.61 | 2 | 0.0037 | 3 | 1054 | 1067 |
| Audrey Caling | SCR | SCR | Service_Clair | Collagen alph | CO1A2_HUM | Collagen alph | 100.00% | 5.86% | 129,296.60 | 72 | 120 | 160 | index=9045 | GPPGAAGAP( | R | G | 99.10% | | 2,894.38 | 3 | 0.0066 | 2.3 | 100 | 132 |
| Audrey Caling | SCR | SCR | Service_Clair | Collagen alph | CO1A2_HUM | Collagen alph | 100.00% | 5.86% | 129,296.60 | 72 | 120 | 160 | index=4667 | GPPGAVGSP( | R | G | 99.70% | Deamidated ( | 3,899.82 | 6 | 0.029 | 7.5 | 907 | 948 |
| Audrey Caling | SCR | SCR | Service_Clair | Collagen alph | CO1A2_HUM | Collagen alph | 100.00% | 5.86% | 129,296.60 | 72 | 120 | 160 | index=4668 | GPPGAVGSP( | R | G | 99.70% | Deamidated ( | 3,899.82 | 6 | 0.029 | 7.5 | 907 | 948 |
| Audrey Caling | SCR | SCR | Service_Clair | Collagen alph | CO1A2_HUM | Collagen alph | 100.00% | 5.86% | 129,296.60 | 72 | 120 | 160 | index=4880 | GPPGAVGSP( | R | G | 99.70% | Deamidated ( | 3,898.81 | 6 | 0.0028 | 0.71 | 907 | 948 |
| Audrey Caling | SCR | SCR | Service_Clair | Collagen alph | CO1A2_HUM | Collagen alph | 100.00% | 5.86% | 129,296.60 | 72 | 120 | 160 | index=4588 | GPPGAVGSP( | R | G | 99.70% | | 3,897.83 | 6 | 0.0097 | 2.5 | 907 | 948 |
| Audrey Caling | SCR | SCR | Service_Clair | Collagen alph | CO1A2_HUM | Collagen alph | 100.00% | 5.86% | 129,296.60 | 72 | 120 | 160 | index=4685 | GPPGAVGSP( | R | G | 99.70% | Deamidated ( | 3,898.82 | 5 | 0.013 | 3.3 | 907 | 948 |
| Audrey Caling | SCR | SCR | Service_Clair | Collagen alph | CO1A2_HUM | Collagen alph | 100.00% | 5.86% | 129,296.60 | 72 | 120 | 160 | index=1268 | GPRGDQGPV | R | T | 99.70% | | 1,094.56 | 3 | 0.0027 | 2.4 | 820 | 830 |
| Audrey Caling | SCR | SCR | Service_Clair | Collagen alph | CO1A2_HUM | Collagen alph | 100.00% | 5.86% | 129,296.60 | 72 | 120 | 160 | index=2514 | GPSGPQGIR( | R | G | 99.70% | | 2,075.05 | 4 | 0.0025 | 1.2 | 997 | 1017 |
| Audrey Caling | SCR | SCR | Service_Clair | Collagen alph | CO1A2_HUM | Collagen alph | 100.00% | 5.86% | 129,296.60 | 72 | 120 | 160 | index=2520 | GPSGPQGIR( | R | G | 99.70% | | 2,075.05 | 3 | 0.0022 | 1.1 | 997 | 1017 |
| Audrey Caling | SCR | SCR | Service_Clair | Collagen alph | CO1A2_HUM | Collagen alph | 100.00% | 5.86% | 129,296.60 | 72 | 120 | 160 | index=1911 | GPTGDPGKN | K | G | 98.70% | Deamidated ( | 1,932.95 | 4 | 0.0059 | 3.1 | 499 | 519 |
| Audrey Caling | SCR | SCR | Service_Clair | Collagen alph | CO1A2_HUM | Collagen alph | 100.00% | 5.86% | 129,296.60 | 72 | 120 | 160 | index=1905 | GPTGDPGKN | K | G | 99.20% | Deamidated ( | 1,932.94 | 5 | 0.0031 | 1.6 | 499 | 519 |
| Audrey Caling | SCR | SCR | Service_Clair | Collagen alph | CO1A2_HUM | Collagen alph | 100.00% | 5.86% | 129,296.60 | 72 | 120 | 160 | index=1647 | GPTGDPGKN | K | G | 99.70% | | 1,931.96 | 3 | 0.0065 | 3.4 | 499 | 519 |
| Audrey Caling | SCR | SCR | Service_Clair | Collagen alph | CO1A2_HUM | Collagen alph | 100.00% | 5.86% | 129,296.60 | 72 | 120 | 160 | index=1910 | GPTGDPGKN | K | G | 99.70% | Deamidated ( | 1,932.95 | 4 | 0.0059 | 3.1 | 499 | 519 |
| Audrey Caling | SCR | SCR | Service_Clair | Collagen alph | CO1A2_HUM | Collagen alph | 100.00% | 5.86% | 129,296.60 | 72 | 120 | 160 | index=1918 | GPTGDPGKN | K | G | 99.70% | Deamidated ( | 1,932.95 | 3 | 0.0054 | 2.8 | 499 | 519 |
| Audrey Caling | SCR | SCR | Service_Clair | Collagen alph | CO1A2_HUM | Collagen alph | 100.00% | 5.86% | 129,296.60 | 72 | 120 | 160 | index=1849 | GRVGAPGPA( | R | G | 99.70% | | 1,064.59 | 3 | 0.0032 | 3 | 223 | 234 |
| Audrey Caling | SCR | SCR | Service_Clair | Collagen alph | CO1A2_HUM | Collagen alph | 100.00% | 5.86% | 129,296.60 | 72 | 120 | 160 | index=1863 | GRVGAPGPA( | R | G | 99.70% | | 1,064.59 | 2 | 0.0025 | 2.3 | 223 | 234 |
| Audrey Caling | SCR | SCR | Service_Clair | Collagen alph | CO1A2_HUM | Collagen alph | 100.00% | 5.86% | 129,296.60 | 72 | 120 | 160 | index=8096 | GSPGERGEV( | R | G | 99.70% | Deamidated ( | 2,618.26 | 3 | 0.0075 | 2.9 | 709 | 738 |
| Audrey Caling | SCR | SCR | Service_Clair | Collagen alph | CO1A2_HUM | Collagen alph | 100.00% | 5.86% | 129,296.60 | 72 | 120 | 160 | index=7504 | GSPGERGEV( | R | G | 99.70% | Deamidated ( | 2,960.42 | 4 | 0.011 | 3.6 | 709 | 741 |
| Audrey Caling | SCR | SCR | Service_Clair | Collagen alph | CO1A2_HUM | Collagen alph | 100.00% | 5.86% | 129,296.60 | 72 | 120 | 160 | index=7505 | GSPGERGEV( | R | G | 99.70% | Deamidated ( | 2,960.42 | 4 | 0.011 | 3.6 | 709 | 741 |
| Audrey Caling | SCR | SCR | Service_Clair | Collagen alph | CO1A2_HUM | Collagen alph | 100.00% | 5.86% | 129,296.60 | 72 | 120 | 160 | index=7875 | GSPGERGEV( | R | G | 99.70% | Deamidated ( | 2,960.42 | 4 | 0.0091 | 3.1 | 709 | 741 |
| Audrey Caling | SCR | SCR | Service_Clair | Collagen alph | CO1A2_HUM | Collagen alph | 100.00% | 5.86% | 129,296.60 | 72 | 120 | 160 | index=3534 | GSPGSRGLP( | R | A | 99.70% | | 1,282.64 | 3 | 0.0044 | 3.5 | 400 | 413 |
| Audrey Caling | SCR | SCR | Service_Clair | Collagen alph | CO1A2_HUM | Collagen alph | 100.00% | 5.86% | 129,296.60 | 72 | 120 | 160 | index=7267 | GSPGSRGLP( | R | G | 97.80% | | 2,192.09 | 3 | 0.0076 | 3.5 | 400 | 423 |
| Audrey Caling | SCR | SCR | Service_Clair | Collagen alph | CO1A2_HUM | Collagen alph | 100.00% | 5.86% | 129,296.60 | 72 | 120 | 160 | index=5452 | GSPGSRGLP( | R | G | 99.70% | Oxidation (+1 | 2,208.08 | 4 | 0.0015 | 0.68 | 400 | 423 |
| Audrey Caling | SCR | SCR | Service_Clair | Collagen alph | CO1A2_HUM | Collagen alph | 100.00% | 5.86% | 129,296.60 | 72 | 120 | 160 | index=2548 | GVVGPQGAR | R | G | 99.70% | | 839.46 | 2 | 0.0015 | 1.8 | 157 | 165 |
| Audrey Caling | SCR | SCR | Service_Clair | Collagen alph | CO1A2_HUM | Collagen alph | 100.00% | 5.86% | 129,296.60 | 72 | 120 | 160 | index=2549 | GVVGPQGAR | R | G | 99.70% | | 839.46 | 2 | 0.0015 | 1.8 | 157 | 165 |
| Audrey Caling | SCR | SCR | Service_Clair | Collagen alph | CO1A2_HUM | Collagen alph | 100.00% | 5.86% | 129,296.60 | 72 | 120 | 160 | index=9838 | GYPGNIGPVG | R | H | 99.70% | | 2,251.16 | 3 | 0.0082 | 3.6 | 949 | 974 |
| Audrey Caling | SCR | SCR | Service_Clair | Collagen alph | CO1A2_HUM | Collagen alph | 100.00% | 5.86% | 129,296.60 | 72 | 120 | 160 | index=9839 | GYPGNIGPVG | R | H | 99.70% | | 2,251.16 | 3 | 0.0082 | 3.6 | 949 | 974 |
| Audrey Caling | SCR | SCR | Service_Clair | Collagen alph | CO1A2_HUM | Collagen alph | 100.00% | 5.86% | 129,296.60 | 72 | 120 | 160 | index=9840 | GYPGNIGPVG | R | H | 99.70% | | 2,251.16 | 3 | 0.0082 | 3.6 | 949 | 974 |
| Audrey Caling | SCR | SCR | Service_Clair | Collagen alph | CO1A2_HUM | Collagen alph | 100.00% | 5.86% | 129,296.60 | 72 | 120 | 160 | index=9846 | GYPGNIGPVG | R | H | 99.70% | | 2,251.16 | 2 | 0.01 | 4.4 | 949 | 974 |
| Audrey Caling | SCR | SCR | Service_Clair | Collagen alph | CO1A2_HUM | Collagen alph | 100.00% | 5.86% | 129,296.60 | 72 | 120 | 160 | index=9352 | GYPGNIGPVG | R | G | 99.70% | | 4,259.16 | 6 | 0.015 | 3.6 | 949 | 996 |
| Audrey Caling | SCR | SCR | Service_Clair | Collagen alph | CO1A2_HUM | Collagen alph | 100.00% | 5.86% | 129,296.60 | 72 | 120 | 160 | index=9321 | GYPGNIGPVG | R | G | 99.70% | | 4,259.17 | 5 | 0.017 | 3.9 | 949 | 996 |
| Audrey Caling | SCR | SCR | Service_Clair | Collagen alph | CO1A2_HUM | Collagen alph | 100.00% | 5.86% | 129,296.60 | 72 | 120 | 160 | index=5301 | HGNRGETGP | K | G | 99.70% | | 2,026.02 | 3 | 0.0065 | 3.2 | 975 | 996 |
| Audrey Caling | SCR | SCR | Service_Clair | Collagen alph | CO1A2_HUM | Collagen alph | 100.00% | 5.86% | 129,296.60 | 72 | 120 | 160 | index=5585 | HGNRGETGP | K | G | 99.70% | Deamidated ( | 2,027.00 | 4 | 0.0078 | 3.9 | 975 | 996 |
| Audrey Caling | SCR | SCR | Service_Clair | Collagen alph | CO1A2_HUM | Collagen alph | 100.00% | 5.86% | 129,296.60 | 72 | 120 | 160 | index=5313 | HGNRGETGP | K | G | 99.70% | | 2,026.02 | 4 | 0.0087 | 4.3 | 975 | 996 |
| Audrey Caling | SCR | SCR | Service_Clair | Collagen alph | CO1A2_HUM | Collagen alph | 100.00% | 5.86% | 129,296.60 | 72 | 120 | 160 | index=5312 | HGNRGETGP | K | G | 99.70% | | 2,026.02 | 4 | 0.0087 | 4.3 | 975 | 996 |
| Audrey Caling | SCR | SCR | Service_Clair | Collagen alph | CO1A2_HUM | Collagen alph | 100.00% | 5.86% | 129,296.60 | 72 | 120 | 160 | index=5300 | HGNRGETGP | K | G | 99.70% | | 2,026.02 | 3 | 0.0065 | 3.2 | 975 | 996 |
| Audrey Caling | SCR | SCR | Service_Clair | Collagen alph | CO1A2_HUM | Collagen alph | 100.00% | 5.86% | 129,296.60 | 72 | 120 | 160 | index=5578 | HGNRGETGP | K | G | 99.70% | Deamidated ( | 2,027.00 | 3 | 0.0036 | 1.8 | 975 | 996 |
| Audrey Caling | SCR | SCR | Service_Clair | Collagen alph | CO1A2_HUM | Collagen alph | 100.00% | 5.86% | 129,296.60 | 72 | 120 | 160 | index=6170 | HGNRGETGP | K | G | 99.70% | | 2,549.25 | 4 | 0.0024 | 0.94 | 975 | 1002 |
| Audrey Caling | SCR | SCR | Service_Clair | Collagen alph | CO1A2_HUM | Collagen alph | 100.00% | 5.86% | 129,296.60 | 72 | 120 | 160 | index=6149 | HGNRGETGP | K | I | 99.70% | | 2,606.27 | 4 | 0.0038 | 1.5 | 975 | 1003 |
| Audrey Caling | SCR | SCR | Service_Clair | Collagen alph | CO1A2_HUM | Collagen alph | 100.00% | 5.86% | 129,296.60 | 72 | 120 | 160 | index=10117 | LPGLKGHNGI | G | T | 99.70% | Acetyl (+42), | 4,602.30 | 7 | 0.024 | 5.3 | 1019 | 1067 |
| Audrey Caling | SCR | SCR | Service_Clair | Collagen alph | CO1A2_HUM | Collagen alph | 100.00% | 5.86% | 129,296.60 | 72 | 120 | 160 | index=10116 | LPGLKGHNGI | G | T | 99.70% | Acetyl (+42), | 4,602.30 | 7 | 0.024 | 5.3 | 1019 | 1067 |
| Audrey Caling | SCR | SCR | Service_Clair | Collagen alph | CO1A2_HUM | Collagen alph | 100.00% | 5.86% | 129,296.60 | 72 | 120 | 160 | index=1605 | NGDKGHAGL | K | G | 99.70% | Deamidated ( | 1,223.60 | 3 | 0.0022 | 1.8 | 507 | 519 |
| Audrey Caling | SCR | SCR | Service_Clair | Collagen alph | CO1A2_HUM | Collagen alph | 100.00% | 5.86% | 129,296.60 | 72 | 120 | 160 | index=10704 | NGPPGPAGS | P | T | 99.70% | Deamidated ( | 2,392.10 | 3 | 0.0066 | 2.8 | 768 | 794 |
| Audrey Caling | SCR | SCR | Service_Clair | Collagen alph | CO1A2_HUM | Collagen alph | 100.00% | 5.86% | 129,296.60 | 72 | 120 | 160 | index=5675 | NSIAYMDEET | K | A | 99.70% | Oxidation (+1 | 1,727.81 | 3 | 0.0031 | 1.8 | 1274 | 1288 |
| Audrey Caling | SCR | SCR | Service_Clair | Collagen alph | CO1A2_HUM | Collagen alph | 100.00% | 5.86% | 129,296.60 | 72 | 120 | 160 | index=9298 | NSIAYMDEET | K | A | 99.70% | | 1,711.82 | 3 | 0.007 | 4.1 | 1274 | 1288 |
| Audrey Caling | SCR | SCR | Service_Clair | Collagen alph | CO1A2_HUM | Collagen alph | 100.00% | 5.86% | 129,296.60 | 72 | 120 | 160 | index=6069 | RGETGPSGP\ | N | G | 99.70% | | 1,717.89 | 3 | 0.0076 | 4.4 | 978 | 996 |
| Audrey Caling | SCR | SCR | Service_Clair | Collagen alph | CO1A2_HUM | Collagen alph | 100.00% | 5.86% | 129,296.60 | 72 | 120 | 160 | index=2644 | SAPSLRPK | R | D | 98.30% | | 854.5 | 2 | 0.0023 | 2.6 | 1124 | 1131 |
| Audrey Caling | SCR | SCR | Service_Clair | Collagen alph | CO1A2_HUM | Collagen alph | 100.00% | 5.86% | 129,296.60 | 72 | 120 | 160 | index=9797 | SAPSLRPKDY | R | S | 99.70% | | 1,888.99 | 4 | 0.0043 | 2.3 | 1124 | 1140 |
| Audrey Caling | SCR | SCR | Service_Clair | Collagen alph | CO1A2_HUM | Collagen alph | 100.00% | 5.86% | 129,296.60 | 72 | 120 | 160 | index=9806 | SAPSLRPKDY | R | S | 99.70% | Phospho (+80 | 1,968.95 | 3 | -0.0052 | -2.7 | 1124 | 1140 |
| Audrey Caling | SCR | SCR | Service_Clair | Collagen alph | CO1A2_HUM | Collagen alph | 100.00% | 5.86% | 129,296.60 | 72 | 120 | 160 | index=9796 | SAPSLRPKDY | R | S | 99.70% | | 1,888.99 | 4 | 0.0043 | 2.3 | 1124 | 1140 |
| Audrey Caling | SCR | SCR | Service_Clair | Collagen alph | CO1A2_HUM | Collagen alph | 100.00% | 5.86% | 129,296.60 | 72 | 120 | 160 | index=9803 | SAPSLRPKDY | R | S | 99.70% | | 1,888.99 | 3 | 0.0045 | 2.4 | 1124 | 1140 |
| Audrey Caling | SCR | SCR | Service_Clair | Collagen alph | CO1A2_HUM | Collagen alph | 100.00% | 5.86% | 129,296.60 | 72 | 120 | 160 | index=9816 | SAPSLRPKDY | R | S | 99.70% | | 1,888.99 | 2 | 0.0046 | 2.4 | 1124 | 1140 |
| Audrey Caling | SCR | SCR | Service_Clair | Collagen alph | CO1A2_HUM | Collagen alph | 100.00% | 5.86% | 129,296.60 | 72 | 120 | 160 | index=9804 | SAPSLRPKDY | R | S | 99.70% | | 1,888.99 | 3 | 0.0045 | 2.4 | 1124 | 1140 |
| Audrey Caling | SCR | SCR | Service_Clair | Collagen alph | CO1A2_HUM | Collagen alph | 100.00% | 5.86% | 129,296.60 | 72 | 120 | 160 | index=19462 | SAPSLRPKDY | R | K | 99.70% | | 3,641.90 | 4 | 0.013 | 3.6 | 1124 | 1156 |
| Audrey Caling | SCR | SCR | Service_Clair | Collagen alph | CO1A2_HUM | Collagen alph | 100.00% | 5.86% | 129,296.60 | 72 | 120 | 160 | index=17850 | SAPSLRPKDY | R | K | 99.70% | Deamidated ( | 3,642.89 | 4 | 0.018 | 4.9 | 1124 | 1156 |
| Audrey Caling | SCR | SCR | Service_Clair | Collagen alph | CO1A2_HUM | Collagen alph | 100.00% | 5.86% | 129,296.60 | 72 | 120 | 160 | index=17435 | SLNNQIETLLT | K | K | 99.70% | | 1,770.92 | 3 | 0.006 | 3.4 | 1141 | 1156 |
| Audrey Caling | SCR | SCR | Service_Clair | Collagen alph | CO1A2_HUM | Collagen alph | 100.00% | 5.86% | 129,296.60 | 72 | 120 | 160 | index=16436 | SLNNQIETLLT | K | K | 99.70% | Deamidated ( | 1,771.90 | 2 | 0.0083 | 4.7 | 1141 | 1156 |
| Audrey Caling | SCR | SCR | Service_Clair | Collagen alph | CO1A2_HUM | Collagen alph | 100.00% | 5.86% | 129,296.60 | 72 | 120 | 160 | index=17436 | SLNNQIETLLT | K | K | 99.70% | | 1,770.92 | 3 | 0.006 | 3.4 | 1141 | 1156 |
| Audrey Caling | SCR | SCR | Service_Clair | Collagen alph | CO1A2_HUM | Collagen alph | 100.00% | 5.86% | 129,296.60 | 72 | 120 | 160 | index=17443 | SLNNQIETLLT | K | K | 99.70% | | 1,770.92 | 2 | 0.0057 | 3.2 | 1141 | 1156 |
| Audrey Caling | SCR | SCR | Service_Clair | Collagen alph | CO1A2_HUM | Collagen alph | 100.00% | 5.86% | 129,296.60 | 72 | 120 | 160 | index=17442 | SLNNQIETLLT | K | K | 99.70% | | 1,770.92 | 2 | 0.0057 | 3.2 | 1141 | 1156 |
| Audrey Caling | SCR | SCR | Service_Clair | Collagen alph | CO1A2_HUM | Collagen alph | 100.00% | 5.86% | 129,296.60 | 72 | 120 | 160 | index=15759 | SLNNQIETLLT | K | T | 99.70% | | 2,337.25 | 4 | 0.0086 | 3.7 | 1141 | 1161 |

| | | | | | | | | | | | | | | | | | | | | | | | | |
|---|---|---|---|---|---|---|---|---|---|---|---|---|---|---|---|---|---|---|---|---|---|---|---|---|
| Audrey Caling | SCR | SCR | Service_Clair | Collagen alph | CO1A2_HUM | Collagen alph | 100.00% | 5.86% | 129,296.60 | 72 | 120 | 160 | index=15758 | SLNNQIETLLT | K | T | 99.70% | | 2,337.25 | 4 | 0.0086 | 3.7 | 1141 | 1161 |
| Audrey Caling | SCR | SCR | Service_Clair | Collagen alph | CO1A2_HUM | Collagen alph | 100.00% | 5.86% | 129,296.60 | 72 | 120 | 160 | index=12931 | TGHPGTVGPA | R | S | 99.70% | Phospho (+80 | 5,592.50 | 5 | 0.015 | 2.7 | 1068 | 1123 |
| Audrey Caling | SCR | SCR | Service_Clair | Collagen alph | CO1A2_HUM | Collagen alph | 100.00% | 5.86% | 129,296.60 | 72 | 120 | 160 | index=7719 | TGPPGPSGIS | R | E | 99.70% | | 1,780.92 | 2 | 0.0043 | 2.4 | 795 | 815 |
| Audrey Caling | SCR | SCR | Service_Clair | Collagen alph | CO1A2_HUM | Collagen alph | 100.00% | 5.86% | 129,296.60 | 72 | 120 | 160 | index=7718 | TGPPGPSGIS | R | E | 99.70% | | 1,780.92 | 2 | 0.0043 | 2.4 | 795 | 815 |
| Audrey Caling | SCR | SCR | Service_Clair | Collagen alph | CO1A2_HUM | Collagen alph | 100.00% | 5.86% | 129,296.60 | 72 | 120 | 160 | index=8877 | TGPPGPSGIS | R | G | 99.70% | | 2,236.17 | 3 | 0.0049 | 2.2 | 795 | 819 |
| Audrey Caling | SCR | SCR | Service_Clair | Collagen alph | CO1A2_HUM | Collagen alph | 100.00% | 5.86% | 129,296.60 | 72 | 120 | 160 | index=8262 | TGPPGPSGIS | R | G | 98.00% | | 2,546.34 | 3 | 0.0016 | 0.63 | 795 | 822 |
| Audrey Caling | SCR | SCR | Service_Clair | Collagen alph | CO1A2_HUM | Collagen alph | 100.00% | 5.86% | 129,296.60 | 72 | 120 | 160 | index=8259 | TGPPGPSGIS | R | G | 99.70% | | 2,546.34 | 4 | 0.0036 | 1.4 | 795 | 822 |
| Audrey Caling | SCR | SCR | Service_Clair | Collagen alph | CO1A2_HUM | Collagen alph | 100.00% | 5.86% | 129,296.60 | 72 | 120 | 160 | index=8181 | TGPPGPSGIS | R | T | 96.60% | | 3,312.71 | 5 | 0.003 | 0.89 | 795 | 830 |
| Audrey Caling | SCR | SCR | Service_Clair | Collagen alph | CO1A2_HUM | Collagen alph | 100.00% | 5.86% | 129,296.60 | 72 | 120 | 160 | index=2132 | VGAPGPAGAI | R | G | 99.70% | | 851.46 | 2 | 0.0016 | 1.9 | 225 | 234 |
| Audrey Caling | SCR | SCR | Service_Clair | Collagen alph | CO1A2_HUM | Collagen alph | 100.00% | 5.86% | 129,296.60 | 72 | 120 | 160 | index=2131 | VGAPGPAGAI | R | G | 99.70% | | 851.46 | 2 | 0.0016 | 1.9 | 225 | 234 |
| Audrey Caling | SCR | SCR | Service_Clair | Collagen alph | CO1A2_HUM | Collagen alph | 100.00% | 5.86% | 129,296.60 | 72 | 120 | 160 | index=13882 | VGPPGNPGA | P | G | 99.70% | Deamidated ( | 2,902.55 | 3 | 0.011 | 3.7 | 294 | 327 |
| Audrey Caling | SCR | SCR | Service_Clair | Collagen alph | CO1A2_HUM | Collagen alph | 100.00% | 5.86% | 129,296.60 | 72 | 120 | 160 | index=10001 | VYCDFSTGET | K | A | 99.70% | Carbamidom | 1,606.68 | 2 | 0.0027 | 1.7 | 1193 | 1205 |
| Audrey Caling | shARF6 | shARF6 | Service_Clair | Collagen alph | CO1A2_HUM | Collagen alph | 100.00% | 4.32% | 129,296.60 | 52 | 82 | 101 | index=1949 | AGEDGHPGK | K | G | 97.60% | | 2,437.24 | 4 | 0.0047 | 1.9 | 141 | 165 |
| Audrey Caling | shARF6 | shARF6 | Service_Clair | Collagen alph | CO1A2_HUM | Collagen alph | 100.00% | 4.32% | 129,296.60 | 52 | 82 | 101 | index=1860 | AGVMGPPGS | R | G | 99.70% | Oxidation (+1 | 943.46 | 2 | 0.001 | 1.1 | 414 | 423 |
| Audrey Caling | shARF6 | shARF6 | Service_Clair | Collagen alph | CO1A2_HUM | Collagen alph | 100.00% | 4.32% | 129,296.60 | 52 | 82 | 101 | index=3862 | AGVMGPPGS | R | G | 99.70% | | 927.46 | 2 | 0.0015 | 1.6 | 414 | 423 |
| Audrey Caling | shARF6 | shARF6 | Service_Clair | Collagen alph | CO1A2_HUM | Collagen alph | 100.00% | 4.32% | 129,296.60 | 52 | 82 | 101 | index=7197 | AGVMGPPGS | R | G | 96.40% | Deamidated ( | 3,412.66 | 5 | 0.01 | 2.9 | 414 | 450 |
| Audrey Caling | shARF6 | shARF6 | Service_Clair | Collagen alph | CO1A2_HUM | Collagen alph | 100.00% | 4.32% | 129,296.60 | 52 | 82 | 101 | index=8494 | AGVMGPPGS | R | G | 99.70% | Deamidated ( | 3,396.66 | 4 | 0.011 | 3.2 | 414 | 450 |
| Audrey Caling | shARF6 | shARF6 | Service_Clair | Collagen alph | CO1A2_HUM | Collagen alph | 100.00% | 4.32% | 129,296.60 | 52 | 82 | 101 | index=8177 | AQPENIPAKN | R | S | 99.70% | | 1,585.80 | 3 | 0.0032 | 2 | 1206 | 1218 |
| Audrey Caling | shARF6 | shARF6 | Service_Clair | Collagen alph | CO1A2_HUM | Collagen alph | 100.00% | 4.32% | 129,296.60 | 52 | 82 | 101 | index=8176 | AQPENIPAKN | R | S | 99.70% | | 1,585.80 | 3 | 0.0032 | 2 | 1206 | 1218 |
| Audrey Caling | shARF6 | shARF6 | Service_Clair | Collagen alph | CO1A2_HUM | Collagen alph | 100.00% | 4.32% | 129,296.60 | 52 | 82 | 101 | index=8164 | AQPENIPAKN | R | S | 99.70% | | 1,585.81 | 2 | 0.0049 | 3.1 | 1206 | 1218 |
| Audrey Caling | shARF6 | shARF6 | Service_Clair | Collagen alph | CO1A2_HUM | Collagen alph | 100.00% | 4.32% | 129,296.60 | 52 | 82 | 101 | index=8175 | AQPENIPAKN | R | S | 99.70% | | 1,585.80 | 3 | 0.0032 | 2 | 1206 | 1218 |
| Audrey Caling | shARF6 | shARF6 | Service_Clair | Collagen alph | CO1A2_HUM | Collagen alph | 100.00% | 4.32% | 129,296.60 | 52 | 82 | 101 | index=18976 | DYEVDATLKS | K | K | 99.50% | | 2,805.41 | 3 | 0.0042 | 1.5 | 1132 | 1156 |
| Audrey Caling | shARF6 | shARF6 | Service_Clair | Collagen alph | CO1A2_HUM | Collagen alph | 100.00% | 4.32% | 129,296.60 | 52 | 82 | 101 | index=3151 | EGLRGPRGD | K | T | 95.60% | | 1,549.81 | 2 | 0.004 | 2.6 | 816 | 830 |
| Audrey Caling | shARF6 | shARF6 | Service_Clair | Collagen alph | CO1A2_HUM | Collagen alph | 100.00% | 4.32% | 129,296.60 | 52 | 82 | 101 | index=3145 | EGLRGPRGD | K | T | 99.70% | | 1,549.81 | 3 | 0.0047 | 3 | 816 | 830 |
| Audrey Caling | shARF6 | shARF6 | Service_Clair | Collagen alph | CO1A2_HUM | Collagen alph | 100.00% | 4.32% | 129,296.60 | 52 | 82 | 101 | index=3144 | EGLRGPRGD | K | T | 99.70% | | 1,549.81 | 3 | 0.0047 | 3 | 816 | 830 |
| Audrey Caling | shARF6 | shARF6 | Service_Clair | Collagen alph | CO1A2_HUM | Collagen alph | 100.00% | 4.32% | 129,296.60 | 52 | 82 | 101 | index=11499 | EMATQLAFMF | K | L | 99.70% | Oxidation (+1 | 1,212.57 | 2 | 0.0042 | 3.4 | 1249 | 1258 |
| Audrey Caling | shARF6 | shARF6 | Service_Clair | Collagen alph | CO1A2_HUM | Collagen alph | 100.00% | 4.32% | 129,296.60 | 52 | 82 | 101 | index=13457 | EMATQLAFMF | K | L | 99.70% | | 1,196.57 | 2 | 0.0051 | 4.2 | 1249 | 1258 |
| Audrey Caling | shARF6 | shARF6 | Service_Clair | Collagen alph | CO1A2_HUM | Collagen alph | 100.00% | 4.32% | 129,296.60 | 52 | 82 | 101 | index=11532 | GAAGLPGVA( | K | G | 99.70% | | 1,513.84 | 2 | 0.0058 | 3.8 | 310 | 327 |
| Audrey Caling | shARF6 | shARF6 | Service_Clair | Collagen alph | CO1A2_HUM | Collagen alph | 100.00% | 4.32% | 129,296.60 | 52 | 82 | 101 | index=8842 | GAKGPKGEN | R | T | 99.70% | Deamidated ( | 4,589.24 | 5 | 0.027 | 5.9 | 742 | 794 |
| Audrey Caling | shARF6 | shARF6 | Service_Clair | Collagen alph | CO1A2_HUM | Collagen alph | 100.00% | 4.32% | 129,296.60 | 52 | 82 | 101 | index=10515 | GAKGPKGEN | R | T | 99.70% | Deamidated ( | 4,573.26 | 5 | 0.042 | 9.1 | 742 | 794 |
| Audrey Caling | shARF6 | shARF6 | Service_Clair | Collagen alph | CO1A2_HUM | Collagen alph | 100.00% | 4.32% | 129,296.60 | 52 | 82 | 101 | index=10422 | GAKGPKGEN | R | T | 99.70% | Deamidated ( | 4,572.26 | 5 | 0.025 | 5.5 | 742 | 794 |
| Audrey Caling | shARF6 | shARF6 | Service_Clair | Collagen alph | CO1A2_HUM | Collagen alph | 100.00% | 4.32% | 129,296.60 | 52 | 82 | 101 | index=8841 | GAKGPKGEN | R | T | 99.70% | Deamidated ( | 4,589.24 | 5 | 0.027 | 5.9 | 742 | 794 |
| Audrey Caling | shARF6 | shARF6 | Service_Clair | Collagen alph | CO1A2_HUM | Collagen alph | 100.00% | 4.32% | 129,296.60 | 52 | 82 | 101 | index=5470 | GAPGAVGAP( | R | A | 98.10% | | 2,088.00 | 2 | 0.0052 | 2.5 | 676 | 701 |
| Audrey Caling | shARF6 | shARF6 | Service_Clair | Collagen alph | CO1A2_HUM | Collagen alph | 100.00% | 4.32% | 129,296.60 | 52 | 82 | 101 | index=5517 | GAPGAVGAP( | R | G | 99.70% | | 2,694.33 | 3 | 0.0053 | 2 | 676 | 708 |
| Audrey Caling | shARF6 | shARF6 | Service_Clair | Collagen alph | CO1A2_HUM | Collagen alph | 100.00% | 4.32% | 129,296.60 | 52 | 82 | 101 | index=5518 | GAPGAVGAP( | R | G | 99.70% | | 2,694.33 | 3 | 0.0053 | 2 | 676 | 708 |
| Audrey Caling | shARF6 | shARF6 | Service_Clair | Collagen alph | CO1A2_HUM | Collagen alph | 100.00% | 4.32% | 129,296.60 | 52 | 82 | 101 | index=5516 | GAPGAVGAP( | R | G | 99.70% | | 2,694.33 | 3 | 0.0053 | 2 | 676 | 708 |
| Audrey Caling | shARF6 | shARF6 | Service_Clair | Collagen alph | CO1A2_HUM | Collagen alph | 100.00% | 4.32% | 129,296.60 | 52 | 82 | 101 | index=5513 | GAPGAVGAP( | R | G | 99.70% | | 2,694.33 | 4 | 0.0045 | 1.7 | 676 | 708 |
| Audrey Caling | shARF6 | shARF6 | Service_Clair | Collagen alph | CO1A2_HUM | Collagen alph | 100.00% | 4.32% | 129,296.60 | 52 | 82 | 101 | index=6514 | GAPGAVGAP( | R | A | 99.20% | | 3,716.81 | 4 | 0.011 | 2.9 | 676 | 719 |
| Audrey Caling | shARF6 | shARF6 | Service_Clair | Collagen alph | CO1A2_HUM | Collagen alph | 100.00% | 4.32% | 129,296.60 | 52 | 82 | 101 | index=1531 | GASGPAGVR | R | G | 99.60% | | 770.41 | 2 | 0.0017 | 2.2 | 424 | 432 |
| Audrey Caling | shARF6 | shARF6 | Service_Clair | Collagen alph | CO1A2_HUM | Collagen alph | 100.00% | 4.32% | 129,296.60 | 52 | 82 | 101 | index=6759 | GASGPAGVR( | R | G | 99.70% | Deamidated ( | 2,503.21 | 3 | 0.0076 | 3 | 424 | 450 |
| Audrey Caling | shARF6 | shARF6 | Service_Clair | Collagen alph | CO1A2_HUM | Collagen alph | 100.00% | 4.32% | 129,296.60 | 52 | 82 | 101 | index=6889 | GASGPAGVR( | R | G | 99.70% | Deamidated ( | 2,503.20 | 3 | 0.0034 | 1.3 | 424 | 450 |
| Audrey Caling | shARF6 | shARF6 | Service_Clair | Collagen alph | CO1A2_HUM | Collagen alph | 100.00% | 4.32% | 129,296.60 | 52 | 82 | 101 | index=5357 | GEPGLRGEIG | K | G | 99.20% | | 1,806.92 | 3 | 0.007 | 3.9 | 658 | 675 |
| Audrey Caling | shARF6 | shARF6 | Service_Clair | Collagen alph | CO1A2_HUM | Collagen alph | 100.00% | 4.32% | 129,296.60 | 52 | 82 | 101 | index=6601 | GIPGPVGAAG | R | G | 99.70% | | 1,250.67 | 3 | 0.0013 | 1 | 328 | 342 |
| Audrey Caling | shARF6 | shARF6 | Service_Clair | Collagen alph | CO1A2_HUM | Collagen alph | 100.00% | 4.32% | 129,296.60 | 52 | 82 | 101 | index=6609 | GIPGPVGAAG | R | G | 99.70% | | 1,250.67 | 2 | -0.00012 | -0.097 | 328 | 342 |
| Audrey Caling | shARF6 | shARF6 | Service_Clair | Collagen alph | CO1A2_HUM | Collagen alph | 100.00% | 4.32% | 129,296.60 | 52 | 82 | 101 | index=6608 | GIPGPVGAAG | R | G | 99.70% | | 1,250.67 | 2 | -0.00012 | -0.097 | 328 | 342 |
| Audrey Caling | shARF6 | shARF6 | Service_Clair | Collagen alph | CO1A2_HUM | Collagen alph | 100.00% | 4.32% | 129,296.60 | 52 | 82 | 101 | index=5167 | GLPGADGRA | R | G | 99.30% | Oxidation (+1 | 1,666.83 | 3 | 0.0061 | 3.6 | 406 | 423 |
| Audrey Caling | shARF6 | shARF6 | Service_Clair | Collagen alph | CO1A2_HUM | Collagen alph | 100.00% | 4.32% | 129,296.60 | 52 | 82 | 101 | index=7087 | GLPGADGRA | R | G | 99.70% | | 1,650.83 | 3 | 0.0016 | 0.96 | 406 | 423 |
| Audrey Caling | shARF6 | shARF6 | Service_Clair | Collagen alph | CO1A2_HUM | Collagen alph | 100.00% | 4.32% | 129,296.60 | 52 | 82 | 101 | index=6360 | GLPGADGRA | R | G | 99.70% | Oxidation (+1 | 1,666.83 | 2 | 0.0067 | 4 | 406 | 423 |
| Audrey Caling | shARF6 | shARF6 | Service_Clair | Collagen alph | CO1A2_HUM | Collagen alph | 100.00% | 4.32% | 129,296.60 | 52 | 82 | 101 | index=7052 | GLPGADGRA | R | G | 99.70% | | 1,650.83 | 3 | 0.004 | 2.4 | 406 | 423 |
| Audrey Caling | shARF6 | shARF6 | Service_Clair | Collagen alph | CO1A2_HUM | Collagen alph | 100.00% | 4.32% | 129,296.60 | 52 | 82 | 101 | index=7092 | GLPGADGRA | R | G | 99.70% | | 1,650.83 | 2 | 0.0043 | 2.6 | 406 | 423 |
| Audrey Caling | shARF6 | shARF6 | Service_Clair | Collagen alph | CO1A2_HUM | Collagen alph | 100.00% | 4.32% | 129,296.60 | 52 | 82 | 101 | index=8180 | GLPGADGRA | R | G | 95.30% | Oxidation (+1 | 4,136.02 | 6 | 0.009 | 2.2 | 406 | 450 |
| Audrey Caling | shARF6 | shARF6 | Service_Clair | Collagen alph | CO1A2_HUM | Collagen alph | 100.00% | 4.32% | 129,296.60 | 52 | 82 | 101 | index=9188 | GLPGADGRA | R | G | 99.70% | Deamidated ( | 4,120.03 | 6 | 0.008 | 1.9 | 406 | 450 |
| Audrey Caling | shARF6 | shARF6 | Service_Clair | Collagen alph | CO1A2_HUM | Collagen alph | 100.00% | 4.32% | 129,296.60 | 52 | 82 | 101 | index=9195 | GLPGADGRA | R | G | 99.70% | Deamidated ( | 4,120.04 | 5 | 0.016 | 3.9 | 406 | 450 |
| Audrey Caling | shARF6 | shARF6 | Service_Clair | Collagen alph | CO1A2_HUM | Collagen alph | 100.00% | 4.32% | 129,296.60 | 52 | 82 | 101 | index=4180 | GLPGERGRV( | R | G | 99.70% | | 1,673.91 | 4 | 0.0056 | 3.4 | 217 | 234 |
| Audrey Caling | shARF6 | shARF6 | Service_Clair | Collagen alph | CO1A2_HUM | Collagen alph | 100.00% | 4.32% | 129,296.60 | 52 | 82 | 101 | index=4176 | GLPGERGRV( | R | G | 99.70% | | 1,673.91 | 3 | 0.0063 | 3.8 | 217 | 234 |
| Audrey Caling | shARF6 | shARF6 | Service_Clair | Collagen alph | CO1A2_HUM | Collagen alph | 100.00% | 4.32% | 129,296.60 | 52 | 82 | 101 | index=6877 | GLPGSPGNIG | R | E | 99.70% | | 1,220.65 | 2 | 0.0015 | 1.2 | 451 | 464 |
| Audrey Caling | shARF6 | shARF6 | Service_Clair | Collagen alph | CO1A2_HUM | Collagen alph | 100.00% | 4.32% | 129,296.60 | 52 | 82 | 101 | index=13166 | GLPGSPGNIG | R | G | 99.70% | | 3,241.74 | 4 | 0.0027 | 0.82 | 451 | 486 |
| Audrey Caling | shARF6 | shARF6 | Service_Clair | Collagen alph | CO1A2_HUM | Collagen alph | 100.00% | 4.32% | 129,296.60 | 52 | 82 | 101 | index=14593 | GLPGVAGAV( | R | G | 99.70% | | 2,066.14 | 2 | 0.0077 | 3.7 | 883 | 906 |
| Audrey Caling | shARF6 | shARF6 | Service_Clair | Collagen alph | CO1A2_HUM | Collagen alph | 100.00% | 4.32% | 129,296.60 | 52 | 82 | 101 | index=5311 | GLVGEPGPA( | R | R | 99.70% | | 3,412.62 | 4 | 0.0094 | 2.8 | 343 | 380 |
| Audrey Caling | shARF6 | shARF6 | Service_Clair | Collagen alph | CO1A2_HUM | Collagen alph | 100.00% | 4.32% | 129,296.60 | 52 | 82 | 101 | index=1115 | GPAGPSGPA( | R | D | 99.70% | | 894.46 | 2 | 0.0015 | 1.7 | 1054 | 1064 |
| Audrey Caling | shARF6 | shARF6 | Service_Clair | Collagen alph | CO1A2_HUM | Collagen alph | 100.00% | 4.32% | 129,296.60 | 52 | 82 | 101 | index=960 | GPAGPSGPA( | R | T | 99.70% | | 1,222.61 | 2 | 0.002 | 1.6 | 1054 | 1067 |
| Audrey Caling | shARF6 | shARF6 | Service_Clair | Collagen alph | CO1A2_HUM | Collagen alph | 100.00% | 4.32% | 129,296.60 | 52 | 82 | 101 | index=957 | GPAGPSGPA( | R | T | 99.70% | | 1,222.61 | 3 | 0.0025 | 2.1 | 1054 | 1067 |
| Audrey Caling | shARF6 | shARF6 | Service_Clair | Collagen alph | CO1A2_HUM | Collagen alph | 100.00% | 4.32% | 129,296.60 | 52 | 82 | 101 | index=4134 | GPPGAVGSP( | R | G | 99.70% | Deamidated ( | 3,898.83 | 5 | 0.02 | 5 | 907 | 948 |
| Audrey Caling | shARF6 | shARF6 | Service_Clair | Collagen alph | CO1A2_HUM | Collagen alph | 100.00% | 4.32% | 129,296.60 | 52 | 82 | 101 | index=1059 | GPRGDQGPV | R | T | 99.70% | | 1,094.56 | 3 | 0.0029 | 2.6 | 820 | 830 |
| Audrey Caling | shARF6 | shARF6 | Service_Clair | Collagen alph | CO1A2_HUM | Collagen alph | 100.00% | 4.32% | 129,296.60 | 52 | 82 | 101 | index=2787 | GPSGPQGIR | R | G | 95.30% | | 867.46 | 2 | 0.0025 | 2.9 | 997 | 1005 |
| Audrey Caling | shARF6 | shARF6 | Service_Clair | Collagen alph | CO1A2_HUM | Collagen alph | 100.00% | 4.32% | 129,296.60 | 52 | 82 | 101 | index=2211 | GPSGPQGIR( | R | G | 99.70% | | 2,075.05 | 4 | 0.0015 | 0.74 | 997 | 1017 |
| Audrey Caling | shARF6 | shARF6 | Service_Clair | Collagen alph | CO1A2_HUM | Collagen alph | 100.00% | 4.32% | 129,296.60 | 52 | 82 | 101 | index=2217 | GPSGPQGIR( | R | G | 99.70% | | 2,075.05 | 3 | 0.0023 | 1.1 | 997 | 1017 |
| Audrey Caling | shARF6 | shARF6 | Service_Clair | Collagen alph | CO1A2_HUM | Collagen alph | 100.00% | 4.32% | 129,296.60 | 52 | 82 | 101 | index=1695 | GPTGDPGKN | K | G | 96.60% | Deamidated ( | 1,932.94 | 5 | 0.0031 | 1.6 | 499 | 519 |
| Audrey Caling | shARF6 | shARF6 | Service_Clair | Collagen alph | CO1A2_HUM | Collagen alph | 100.00% | 4.32% | 129,296.60 | 52 | 82 | 101 | index=1714 | GPTGDPGKN | K | G | 99.70% | Deamidated ( | 1,932.94 | 4 | 0.0026 | 1.3 | 499 | 519 |
| Audrey Caling | shARF6 | shARF6 | Service_Clair | Collagen alph | CO1A2_HUM | Collagen alph | 100.00% | 4.32% | 129,296.60 | 52 | 82 | 101 | index=1694 | GPTGDPGKN | K | G | 99.70% | Deamidated ( | 1,932.94 | 3 | 0.0032 | 1.6 | 499 | 519 |
| Audrey Caling | shARF6 | shARF6 | Service_Clair | Collagen alph | CO1A2_HUM | Collagen alph | 100.00% | 4.32% | 129,296.60 | 52 | 82 | 101 | index=1609 | GRVGAPGPA( | R | G | 99.70% | | 1,064.59 | 2 | 0.003 | 2.8 | 223 | 234 |
| Audrey Caling | shARF6 | shARF6 | Service_Clair | Collagen alph | CO1A2_HUM | Collagen alph | 100.00% | 4.32% | 129,296.60 | 52 | 82 | 101 | index=1603 | GRVGAPGPA( | R | G | 99.70% | | 1,064.59 | 3 | 0.0012 | 1.1 | 223 | 234 |
| Audrey Caling | shARF6 | shARF6 | Service_Clair | Collagen alph | CO1A2_HUM | Collagen alph | 100.00% | 4.32% | 129,296.60 | 52 | 82 | 101 | index=6735 | GSPGERGEV( | R | G | 99.70% | Deamidated ( | 2,960.42 | 4 | 0.0043 | 1.5 | 709 | 741 |
| Audrey Caling | shARF6 | shARF6 | Service_Clair | Collagen alph | CO1A2_HUM | Collagen alph | 100.00% | 4.32% | 129,296.60 | 52 | 82 | 101 | index=3108 | GSPGSRGLP( | R | A | 99.70% | | 1,282.64 | 3 | 0.0038 | 3 | 400 | 413 |
| Audrey Caling | shARF6 | shARF6 | Service_Clair | Collagen alph | CO1A2_HUM | Collagen alph | 100.00% | 4.32% | 129,296.60 | 52 | 82 | 101 | index=6544 | GSPGSRGLP( | R | G | 99.70% | | 2,192.09 | 4 | 0.004 | 1.8 | 400 | 423 |
| Audrey Caling | shARF6 | shARF6 | Service_Clair | Collagen alph | CO1A2_HUM | Collagen alph | 100.00% | 4.32% | 129,296.60 | 52 | 82 | 101 | index=6470 | GVRGPNGDA | A | G | 99.70% | Deamidated ( | 2,047.01 | 4 | 0.0061 | 3 | 430 | 450 |
| Audrey Caling | shARF6 | shARF6 | Service_Clair | Collagen alph | CO1A2_HUM | Collagen alph | 100.00% | 4.32% | 129,296.60 | 52 | 82 | 101 | index=2242 | GVVGPQGAR | R | G | 99.70% | | 839.46 | 2 | 0.0019 | 2.3 | 157 | 165 |
| Audrey Caling | shARF6 | shARF6 | Service_Clair | Collagen alph | CO1A2_HUM | Collagen alph | 100.00% | 4.32% | 129,296.60 | 52 | 82 | 101 | index=2241 | GVVGPQGAR | R | G | 99.70% | | 839.46 | 2 | 0.0019 | 2.3 | 157 | 165 |
| Audrey Caling | shARF6 | shARF6 | Service_Clair | Collagen alph | CO1A2_HUM | Collagen alph | 100.00% | 4.32% | 129,296.60 | 52 | 82 | 101 | index=8838 | GYPGNIGPVG | R | H | 99.70% | | 2,251.16 | 3 | 0.006 | 2.7 | 949 | 974 |
| Audrey Caling | shARF6 | shARF6 | Service_Clair | Collagen alph | CO1A2_HUM | Collagen alph | 100.00% | 4.32% | 129,296.60 | 52 | 82 | 101 | index=8839 | GYPGNIGPVG | R | H | 99.70% | | 2,251.16 | 3 | 0.006 | 2.7 | 949 | 974 |
| Audrey Caling | shARF6 | shARF6 | Service_Clair | Collagen alph | CO1A2_HUM | Collagen alph | 100.00% | 4.32% | 129,296.60 | 52 | 82 | 101 | index=8399 | GYPGNIGPVG | R | G | 99.70% | | 4,259.17 | 5 | 0.025 | 5.8 | 949 | 996 |
| Audrey Caling | shARF6 | shARF6 | Service_Clair | Collagen alph | CO1A2_HUM | Collagen alph | 100.00% | 4.32% | 129,296.60 | 52 | 82 | 101 | index=4742 | HGNRGETGP | K | G | 99.70% | | 2,026.01 | 3 | 0.0054 | 2.7 | 975 | 996 |
| Audrey Caling | shARF6 | shARF6 | Service_Clair | Collagen alph | CO1A2_HUM | Collagen alph | 100.00% | 4.32% | 129,296.60 | 52 | 82 | 101 | index=4736 | HGNRGETGP | K | G | 99.70% | | 2,026.02 | 4 | 0.0061 | 3 | 975 | 996 |
| Audrey Caling | shARF6 | shARF6 | Service_Clair | Collagen alph | CO1A2_HUM | Collagen alph | 100.00% | 4.32% | 129,296.60 | 52 | 82 | 101 | index=4735 | HGNRGETGP | K | G | 99.70% | | 2,026.02 | 4 | 0.0061 | 3 | 975 | 996 |
| Audrey Caling | shARF6 | shARF6 | Service_Clair | Collagen alph | CO1A2_HUM | Collagen alph | 100.00% | 4.32% | 129,296.60 | 52 | 82 | 101 | index=4741 | HGNRGETGP | K | G | 99.70% | | 2,026.01 | 3 | 0.0054 | 2.7 | 975 | 996 |
| Audrey Caling | shARF6 | shARF6 | Service_Clair | Collagen alph | CO1A2_HUM | Collagen alph | 100.00% | 4.32% | 129,296.60 | 52 | 82 | 101 | index=4984 | HGNRGETGP | K | G | 99.70% | Deamidated ( | 2,027.00 | 3 | 0.0041 | 2 | 975 | 996 |
| Audrey Caling | shARF6 | shARF6 | Service_Clair | Collagen alph | CO1A2_HUM | Collagen alph | 100.00% | 4.32% | 129,296.60 | 52 | 82 | 101 | index=5467 | HGNRGETGP | K | G | 99.70% | | 2,549.26 | 4 | 0.0081 | 3.2 | 975 | 1002 |
| Audrey Caling | shARF6 | shARF6 | Service_Clair | Collagen alph | CO1A2_HUM | Collagen alph | 100.00% | 4.32% | 129,296.60 | 52 | 82 | 101 | index=5469 | HGNRGETGP | K | G | 99.70% | | 2,549.26 | 3 | 0.0093 | 3.6 | 975 | 1002 |
| Audrey Caling | shARF6 | shARF6 | Service_Clair | Collagen alph | CO1A2_HUM | Collagen alph | 100.00% | 4.32% | 129,296.60 | 52 | 82 | 101 | index=9115 | LPGLKGHNGI | G | T | 98.30% | Acetyl (+42), | 4,602.30 | 7 | 0.02 | 4.5 | 1019 | 1067 |
| Audrey Caling | shARF6 | shARF6 | Service_Clair | Collagen alph | CO1A2_HUM | Collagen alph | 100.00% | 4.32% | 129,296.60 | 52 | 82 | 101 | index=9116 | LPGLKGHNGI | G | T | 99.10% | Acetyl (+42), | 4,602.30 | 7 | 0.02 | 4.5 | 1019 | 1067 |
| Audrey Caling | shARF6 | shARF6 | Service_Clair | Collagen alph | CO1A2_HUM | Collagen alph | 100.00% | 4.32% | 129,296.60 | 52 | 82 | 101 | index=8347 | NSIAYMDEET | K | A | 99.70% | | 1,711.81 | 3 | 0.0044 | 2.6 | 1274 | 1288 |
| Audrey Caling | shARF6 | shARF6 | Service_Clair | Collagen alph | CO1A2_HUM | Collagen alph | 100.00% | 4.32% | 129,296.60 | 52 | 82 | 101 | index=5409 | RGETGPSGPV | N | G | 99.70% | | 1,717.89 | 3 | 0.005 | 2.9 | 978 | 996 |
| Audrey Caling | shARF6 | shARF6 | Service_Clair | Collagen alph | CO1A2_HUM | Collagen alph | 100.00% | 4.32% | 129,296.60 | 52 | 82 | 101 | index=8820 | SAPSLRPKDY | R | S | 99.70% | | 1,888.99 | 4 | 0.0038 | 2 | 1124 | 1140 |
| Audrey Caling | shARF6 | shARF6 | Service_Clair | Collagen alph | CO1A2_HUM | Collagen alph | 100.00% | 4.32% | 129,296.60 | 52 | 82 | 101 | index=8821 | SAPSLRPKDY | R | S | 99.70% | | 1,888.99 | 4 | 0.0038 | 2 | 1124 | 1140 |
| Audrey Caling | shARF6 | shARF6 | Service_Clair | Collagen alph | CO1A2_HUM | Collagen alph | 100.00% | 4.32% | 129,296.60 | 52 | 82 | 101 | index=8832 | SAPSLRPKDY | R | S | 99.70% | | 1,889.00 | 3 | 0.0056 | 3 | 1124 | 1140 |
| Audrey Caling | shARF6 | shARF6 | Service_Clair | Collagen alph | CO1A2_HUM | Collagen alph | 100.00% | 4.32% | 129,296.60 | 52 | 82 | 101 | index=8835 | SAPSLRPKDY | R | S | 99.70% | Phospho (+80 | 1,968.95 | 3 | -0.0039 | -2 | 1124 | 1140 |
| Audrey Caling | shARF6 | shARF6 | Service_Clair | Collagen alph | CO1A2_HUM | Collagen alph | 100.00% | 4.32% | 129,296.60 | 52 | 82 | 101 | index=8831 | SAPSLRPKDY | R | S | 99.70% | | 1,889.00 | 3 | 0.0056 | 3 | 1124 | 1140 |
| Audrey Caling | shARF6 | shARF6 | Service_Clair | Collagen alph | CO1A2_HUM | Collagen alph | 100.00% | 4.32% | 129,296.60 | 52 | 82 | 101 | index=8807 | SAPSLRPKDY | R | S | 99.70% | | 1,889.00 | 2 | 0.0063 | 3.3 | 1124 | 1140 |
| Audrey Caling | shARF6 | shARF6 | Service_Clair | Collagen alph | CO1A2_HUM | Collagen alph | 100.00% | 4.32% | 129,296.60 | 52 | 82 | 101 | index=8833 | SAPSLRPKDY | R | S | 99.70% | | 1,889.00 | 3 | 0.0056 | 3 | 1124 | 1140 |
| Audrey Caling | shARF6 | shARF6 | Service_Clair | Collagen alph | CO1A2_HUM | Collagen alph | 100.00% | 4.32% | 129,296.60 | 52 | 82 | 101 | index=17997 | SAPSLRPKDY | R | K | 97.00% | Deamidated ( | 3,642.91 | 5 | 0.033 | 9 | 1124 | 1156 |
| Audrey Caling | shARF6 | shARF6 | Service_Clair | Collagen alph | CO1A2_HUM | Collagen alph | 100.00% | 4.32% | 129,296.60 | 52 | 82 | 101 | index=18006 | SAPSLRPKDY | R | K | 99.10% | | 3,641.90 | 5 | 0.0092 | 2.5 | 1124 | 1156 |
| Audrey Caling | shARF6 | shARF6 | Service_Clair | Collagen alph | CO1A2_HUM | Collagen alph | 100.00% | 4.32% | 129,296.60 | 52 | 82 | 101 | index=18001 | SAPSLRPKDY | R | K | 99.70% | | 3,641.91 | 4 | 0.02 | 5.5 | 1124 | 1156 |
| Audrey Caling | shARF6 | shARF6 | Service_Clair | Collagen alph | CO1A2_HUM | Collagen alph | 100.00% | 4.32% | 129,296.60 | 52 | 82 | 101 | index=16366 | SAPSLRPKDY | R | K | 99.70% | Deamidated ( | 3,642.89 | 4 | 0.014 | 4 | 1124 | 1156 |
| Audrey Caling | shARF6 | shARF6 | Service_Clair | Collagen alph | CO1A2_HUM | Collagen alph | 100.00% | 4.32% | 129,296.60 | 52 | 82 | 101 | index=15978 | SLNNQIETLLT | K | K | 99.70% | | 1,770.92 | 3 | 0.0043 | 2.4 | 1141 | 1156 |
| Audrey Caling | shARF6 | shARF6 | Service_Clair | Collagen alph | CO1A2_HUM | Collagen alph | 100.00% | 4.32% | 129,296.60 | 52 | 82 | 101 | index=15975 | SLNNQIETLLT | K | K | 99.70% | | 1,770.92 | 2 | 0.0047 | 2.7 | 1141 | 1156 |
| Audrey Caling | shARF6 | shARF6 | Service_Clair | Collagen alph | CO1A2_HUM | Collagen alph | 100.00% | 4.32% | 129,296.60 | 52 | 82 | 101 | index=14372 | SLNNQIETLLT | K | T | 99.70% | | 2,337.25 | 4 | 0.0054 | 2.3 | 1141 | 1161 |
| Audrey Caling | shARF6 | shARF6 | Service_Clair | Collagen alph | CO1A2_HUM | Collagen alph | 100.00% | 4.32% | 129,296.60 | 52 | 82 | 101 | index=11601 | TGHPGTVGPA | R | S | 99.70% | Phospho (+80 | 5,512.55 | 6 | 0.026 | 4.7 | 1068 | 1123 |
| Audrey Caling | shARF6 | shARF6 | Service_Clair | Collagen alph | CO1A2_HUM | Collagen alph | 100.00% | 4.32% | 129,296.60 | 52 | 82 | 101 | index=6890 | TGPPGPSGIS | R | E | 99.70% | | 1,780.91 | 2 | 0.0013 | 0.72 | 795 | 815 |
| Audrey Caling | shARF6 | shARF6 | Service_Clair | Collagen alph | CO1A2_HUM | Collagen alph | 100.00% | 4.32% | 129,296.60 | 52 | 82 | 101 | index=6891 | TGPPGPSGIS | R | E | 99.70% | | 1,780.91 | 2 | 0.0013 | 0.72 | 795 | 815 |
| Audrey Caling | shARF6 | shARF6 | Service_Clair | Collagen alph | CO1A2_HUM | Collagen alph | 100.00% | 4.32% | 129,296.60 | 52 | 82 | 101 | index=7390 | TGPPGPSGIS | R | G | 99.70% | | 2,546.34 | 4 | 0.0012 | 0.48 | 795 | 822 |
| Audrey Caling | shARF6 | shARF6 | Service_Clair | Collagen alph | CO1A2_HUM | Collagen alph | 100.00% | 4.32% | 129,296.60 | 52 | 82 | 101 | index=7295 | TGPPGPSGIS | R | T | 99.70% | | 3,312.71 | 5 | 0.003 | 0.89 | 795 | 830 |
| Audrey Caling | shARF6 | shARF6 | Service_Clair | Collagen alph | CO1A2_HUM | Collagen alph | 100.00% | 4.32% | 129,296.60 | 52 | 82 | 101 | index=1857 | VGAPGPAGAI | R | G | 99.70% | | 851.46 | 2 | 0.0012 | 1.4 | 225 | 234 |
| Audrey Caling | shARF6 | shARF6 | Service_Clair | Collagen alph | CO1A2_HUM | Collagen alph | 100.00% | 4.32% | 129,296.60 | 52 | 82 | 101 | index=1856 | VGAPGPAGAI | R | G | 99.70% | | 851.46 | 2 | 0.0012 | 1.4 | 225 | 234 |

| Experiment na | Biological sar | Biological sar | MS/MS sampl | Protein group | Protein acces | Protein name | Protein identi | Protein perce | Protein mole | Number of un | Number of un | Number of tot | Spectrum nar | Peptide seque | Previous amir | Next amino ac | Peptide identi | Modifications | Actual peptid | Spectrum cha | Actual minus | Actual minus | Peptide start i | Peptide stop index |
|---|---|---|---|---|---|---|---|---|---|---|---|---|---|---|---|---|---|---|---|---|---|---|---|---|
| Audrey Caling | SCR | SCR | Service_Clair | Collagen alph | CO3A1_HUM | Collagen alph | 100.00% | 0.29% | 138,547.00 | 6 | 7 | 8 | index=4377 | GAAGEPGRD | R | G | 99.70% | Oxidation (+1 | 1,652.77 | 3 | 0.0016 | 0.96 | 519 | 536 |
| Audrey Caling | SCR | SCR | Service_Clair | Collagen alph | CO3A1_HUM | Collagen alph | 100.00% | 0.29% | 138,547.00 | 6 | 7 | 8 | index=5323 | GAAGEPGRD | R | G | 99.70% | | 1,636.78 | 3 | 0.0039 | 2.4 | 519 | 536 |
| Audrey Caling | SCR | SCR | Service_Clair | Collagen alph | CO3A1_HUM | Collagen alph | 100.00% | 0.29% | 138,547.00 | 6 | 7 | 8 | index=4376 | GAPGERGRP | R | G | 99.70% | | 1,645.88 | 3 | 0.0043 | 2.6 | 303 | 320 |
| Audrey Caling | SCR | SCR | Service_Clair | Collagen alph | CO3A1_HUM | Collagen alph | 100.00% | 0.29% | 138,547.00 | 6 | 7 | 8 | index=4316 | GRPGLPGAA | R | G | 99.70% | | 1,078.60 | 3 | 0.0023 | 2.1 | 309 | 320 |
| Audrey Caling | SCR | SCR | Service_Clair | Collagen alph | CO3A1_HUM | Collagen alph | 100.00% | 0.29% | 138,547.00 | 6 | 7 | 8 | index=3607 | GRPGLPGAA | R | G | 99.70% | | 1,648.86 | 3 | 0.0063 | 3.8 | 309 | 326 |
| Audrey Caling | SCR | SCR | Service_Clair | Collagen alph | CO3A1_HUM | Collagen alph | 100.00% | 0.29% | 138,547.00 | 6 | 7 | 8 | index=3058 | SGDRGESGP | K | G | 99.70% | | 1,906.90 | 3 | 0.0072 | 3.8 | 1061 | 1082 |
| Audrey Caling | SCR | SCR | Service_Clair | Collagen alph | CO3A1_HUM | Collagen alph | 100.00% | 0.29% | 138,547.00 | 6 | 7 | 8 | index=3057 | SGDRGESGP | K | G | 99.70% | | 1,906.90 | 3 | 0.0072 | 3.8 | 1061 | 1082 |
| Audrey Caling | SCR | SCR | Service_Clair | Collagen alph | CO3A1_HUM | Collagen alph | 100.00% | 0.29% | 138,547.00 | 6 | 7 | 8 | index=13677 | SVNGQIESLIS | K | K | 99.70% | Deamidated ( | 1,658.82 | 2 | 0.0038 | 2.3 | 1240 | 1255 |
| Audrey Caling | shARF6 | shARF6 | Service_Clair | Collagen alph | CO3A1_HUM | Collagen alph | 100.00% | 0.21% | 138,547.00 | 5 | 5 | 5 | index=4737 | GAAGEPGRD | R | G | 99.70% | | 1,636.78 | 3 | 0.0041 | 2.5 | 519 | 536 |
| Audrey Caling | shARF6 | shARF6 | Service_Clair | Collagen alph | CO3A1_HUM | Collagen alph | 100.00% | 0.21% | 138,547.00 | 5 | 5 | 5 | index=3943 | GAPGERGRP | R | G | 99.70% | | 1,645.88 | 3 | 0.0065 | 3.9 | 303 | 320 |
| Audrey Caling | shARF6 | shARF6 | Service_Clair | Collagen alph | CO3A1_HUM | Collagen alph | 100.00% | 0.21% | 138,547.00 | 5 | 5 | 5 | index=3856 | GRPGLPGAA | R | G | 99.70% | | 1,078.60 | 3 | 0.0019 | 1.8 | 309 | 320 |
| Audrey Caling | shARF6 | shARF6 | Service_Clair | Collagen alph | CO3A1_HUM | Collagen alph | 100.00% | 0.21% | 138,547.00 | 5 | 5 | 5 | index=2649 | SGDRGESGP | K | G | 99.70% | | 1,906.90 | 3 | 0.0076 | 4 | 1061 | 1082 |
| Audrey Caling | shARF6 | shARF6 | Service_Clair | Collagen alph | CO3A1_HUM | Collagen alph | 100.00% | 0.21% | 138,547.00 | 5 | 5 | 5 | index=12401 | SVNGQIESLIS | K | K | 99.70% | Deamidated ( | 1,658.82 | 2 | 0.0056 | 3.4 | 1240 | 1255 |

| Experiment na | Biological sar | Biological sar | MS/MS sampl | Protein group | Protein acces | Protein name | Protein identi | Protein perce | Protein mole | Number of un | Number of un | Number of tot | Spectrum nar | Peptide seque | Previous amir | Next amino ac | Peptide identi | Modifications | Actual peptid | Spectrum cha | Actual minus | Actual minus | Peptide start i | Peptide stop index |
|---|---|---|---|---|---|---|---|---|---|---|---|---|---|---|---|---|---|---|---|---|---|---|---|---|
| Audrey Calin | SCR | SCR | Service_Clair | Collagen alph | CO4A2_HUM | Collagen alph | 100.00% | 0.48% | 167,539.60 | 9 | 10 | 13 | index=15155 | EGFPGPPGFI | R | G | 98.80% | | 1,326.67 | 2 | 0.0021 | 1.6 | 728 | 740 |
| Audrey Calin | SCR | SCR | Service_Clair | Collagen alph | CO4A2_HUM | Collagen alph | 100.00% | 0.48% | 167,539.60 | 9 | 10 | 13 | index=15049 | GLPGDAGRE | R | G | 99.50% | | 2,050.05 | 3 | 0.0081 | 4 | 720 | 740 |
| Audrey Calin | SCR | SCR | Service_Clair | Collagen alph | CO4A2_HUM | Collagen alph | 100.00% | 0.48% | 167,539.60 | 9 | 10 | 13 | index=12553 | GLRGIPGFAG | K | G | 99.70% | | 1,650.86 | 3 | 0.0031 | 1.9 | 702 | 719 |
| Audrey Calin | SCR | SCR | Service_Clair | Collagen alph | CO4A2_HUM | Collagen alph | 100.00% | 0.48% | 167,539.60 | 9 | 10 | 13 | index=10312 | GVSGFPGAD | R | G | 99.70% | | 1,915.94 | 3 | 0.0062 | 3.2 | 109 | 129 |
| Audrey Calin | SCR | SCR | Service_Clair | Collagen alph | CO4A2_HUM | Collagen alph | 100.00% | 0.48% | 167,539.60 | 9 | 10 | 13 | index=10311 | GVSGFPGAD | R | G | 99.70% | | 1,915.94 | 3 | 0.0062 | 3.2 | 109 | 129 |
| Audrey Calin | SCR | SCR | Service_Clair | Collagen alph | CO4A2_HUM | Collagen alph | 100.00% | 0.48% | 167,539.60 | 9 | 10 | 13 | index=6017 | IAVQPGTVGP | K | R | 99.70% | | 1,278.71 | 2 | 0.0049 | 3.8 | 1397 | 1409 |
| Audrey Calin | SCR | SCR | Service_Clair | Collagen alph | CO4A2_HUM | Collagen alph | 100.00% | 0.48% | 167,539.60 | 9 | 10 | 13 | index=7728 | RGPPGAPGE | R | G | 99.70% | Oxidation (+1 | 2,160.02 | 3 | 0.0038 | 1.8 | 1410 | 1431 |
| Audrey Calin | SCR | SCR | Service_Clair | Collagen alph | CO4A2_HUM | Collagen alph | 100.00% | 0.48% | 167,539.60 | 9 | 10 | 13 | index=6854 | RGPPGAPGE | R | A | 99.70% | Oxidation (+1 | 2,570.25 | 4 | 0.0078 | 3 | 1410 | 1436 |
| Audrey Calin | SCR | SCR | Service_Clair | Collagen alph | CO4A2_HUM | Collagen alph | 100.00% | 0.48% | 167,539.60 | 9 | 10 | 13 | index=6801 | RGPPGAPGE | R | A | 99.70% | Oxidation (+1 | 2,570.25 | 4 | 0.0082 | 3.2 | 1410 | 1436 |
| Audrey Calin | SCR | SCR | Service_Clair | Collagen alph | CO4A2_HUM | Collagen alph | 100.00% | 0.48% | 167,539.60 | 9 | 10 | 13 | index=7034 | RGPPGAPGE | R | G | 99.30% | Oxidation (+1 | 3,136.55 | 5 | 0.013 | 4.3 | 1410 | 1442 |
| Audrey Calin | SCR | SCR | Service_Clair | Collagen alph | CO4A2_HUM | Collagen alph | 100.00% | 0.48% | 167,539.60 | 9 | 10 | 13 | index=6436 | RGPPGAPGE | R | G | 99.70% | Oxidation (+1 | 3,136.54 | 5 | 0.0063 | 2 | 1410 | 1442 |
| Audrey Calin | SCR | SCR | Service_Clair | Collagen alph | CO4A2_HUM | Collagen alph | 100.00% | 0.48% | 167,539.60 | 9 | 10 | 13 | index=7632 | RGPPGAPGE | R | G | 99.70% | | 3,120.56 | 4 | 0.016 | 5 | 1410 | 1442 |
| Audrey Calin | SCR | SCR | Service_Clair | Collagen alph | CO4A2_HUM | Collagen alph | 100.00% | 0.48% | 167,539.60 | 9 | 10 | 13 | index=17014 | SVSIGYLLVK | R | H | 99.70% | | 1,077.65 | 2 | 0.0042 | 3.9 | 1485 | 1494 |
| Audrey Calin | shARF6 | shARF6 | Service_Clair | Collagen alph | CO4A2_HUM | Collagen alph | 100.00% | 0.13% | 167,539.60 | 3 | 3 | 3 | index=11320 | GLRGIPGFAG | K | G | 99.70% | | 1,650.86 | 3 | 0.002 | 1.2 | 702 | 719 |
| Audrey Calin | shARF6 | shARF6 | Service_Clair | Collagen alph | CO4A2_HUM | Collagen alph | 100.00% | 0.13% | 167,539.60 | 3 | 3 | 3 | index=9274 | GVSGFPGAD | R | G | 99.70% | | 1,915.94 | 3 | 0.0062 | 3.2 | 109 | 129 |
| Audrey Calin | shARF6 | shARF6 | Service_Clair | Collagen alph | CO4A2_HUM | Collagen alph | 100.00% | 0.13% | 167,539.60 | 3 | 3 | 3 | index=5351 | IAVQPGTVGP | K | R | 99.70% | | 1,278.71 | 2 | 0.0056 | 4.4 | 1397 | 1409 |

| Experiment na | Biological sar | Biological sar | MS/MS sampl | Protein group | Protein acces | Protein name | Protein identi | Protein perce | Protein molec | Number of un | Number of un | Number of tot | Spectrum nar | Peptide seque | Previous amir | Next amino ac | Peptide identi | Modifications | Actual peptide | Spectrum cha | Actual minus | Actual minus | Peptide start i | Peptide stop index |
|---|---|---|---|---|---|---|---|---|---|---|---|---|---|---|---|---|---|---|---|---|---|---|---|---|
| Audrey Caling | SCR | SCR | Service_Clair | Collagen alpł | CO5A1_HUM. | Collagen alpł | 100.00% | 0.51% | 183,544.70 | 13 | 14 | 14 | index=19075 | DAQLSAPTKQ | K | A | 99.60% | | 3,037.57 | 3 | 0.0096 | 3.2 | 80 | 107 |
| Audrey Caling | SCR | SCR | Service_Clair | Collagen alpł | CO5A1_HUM. | Collagen alpł | 100.00% | 0.51% | 183,544.70 | 13 | 14 | 14 | index=18506 | DAQLSAPTKQ | K | K | 99.70% | | 3,236.71 | 4 | 0.01 | 3.2 | 80 | 109 |
| Audrey Caling | SCR | SCR | Service_Clair | Collagen alpł | CO5A1_HUM. | Collagen alpł | 100.00% | 0.51% | 183,544.70 | 13 | 14 | 14 | index=13460 | FGGGGDAGS | R | L | 99.70% | Oxidation (+1 | 2,861.38 | 3 | 0.012 | 4.2 | 526 | 554 |
| Audrey Caling | SCR | SCR | Service_Clair | Collagen alpł | CO5A1_HUM. | Collagen alpł | 100.00% | 0.51% | 183,544.70 | 13 | 14 | 14 | index=2274 | GETGFQGK | R | T | 99.00% | | 822.39 | 2 | 0.0018 | 2.2 | 982 | 989 |
| Audrey Caling | SCR | SCR | Service_Clair | Collagen alpł | CO5A1_HUM. | Collagen alpł | 100.00% | 0.51% | 183,544.70 | 13 | 14 | 14 | index=8102 | GINLSDGKWI | R | I | 99.70% | | 1,281.66 | 3 | 0.0014 | 1.1 | 157 | 167 |
| Audrey Caling | SCR | SCR | Service_Clair | Collagen alpł | CO5A1_HUM. | Collagen alpł | 100.00% | 0.51% | 183,544.70 | 13 | 14 | 14 | index=5123 | GLPGEPGPR | R | G | 99.30% | | 878.46 | 2 | 0.0023 | 2.6 | 673 | 681 |
| Audrey Caling | SCR | SCR | Service_Clair | Collagen alpł | CO5A1_HUM. | Collagen alpł | 100.00% | 0.51% | 183,544.70 | 13 | 14 | 14 | index=20198 | ILDEEVFEGDI | R | A | 99.70% | | 2,501.25 | 3 | 0.0078 | 3.1 | 211 | 231 |
| Audrey Caling | SCR | SCR | Service_Clair | Collagen alpł | CO5A1_HUM. | Collagen alpł | 100.00% | 0.51% | 183,544.70 | 13 | 14 | 14 | index=2416 | RSSKGPDVAY | R | V | 99.70% | | 1,234.65 | 3 | 0.0038 | 3 | 66 | 76 |
| Audrey Caling | SCR | SCR | Service_Clair | Collagen alpł | CO5A1_HUM. | Collagen alpł | 100.00% | 0.51% | 183,544.70 | 13 | 14 | 14 | index=4398 | SSKGPDVAYF | R | D | 99.70% | | 1,406.76 | 3 | 0.0033 | 2.3 | 67 | 79 |
| Audrey Caling | SCR | SCR | Service_Clair | Collagen alpł | CO5A1_HUM. | Collagen alpł | 100.00% | 0.51% | 183,544.70 | 13 | 14 | 14 | index=4405 | SSKGPDVAYF | R | D | 99.70% | | 1,406.76 | 2 | 0.0046 | 3.3 | 67 | 79 |
| Audrey Caling | SCR | SCR | Service_Clair | Collagen alpł | CO5A1_HUM. | Collagen alpł | 100.00% | 0.51% | 183,544.70 | 13 | 14 | 14 | index=3945 | TTGFCATR | K | R | 99.00% | Carbamidom | 912.42 | 2 | 0.0027 | 2.9 | 58 | 65 |
| Audrey Caling | SCR | SCR | Service_Clair | Collagen alpł | CO5A1_HUM. | Collagen alpł | 100.00% | 0.51% | 183,544.70 | 13 | 14 | 14 | index=2931 | TTGFCATRR | K | S | 99.00% | Carbamidom | 1,068.52 | 3 | 0.0019 | 1.8 | 58 | 66 |
| Audrey Caling | SCR | SCR | Service_Clair | Collagen alpł | CO5A1_HUM. | Collagen alpł | 100.00% | 0.51% | 183,544.70 | 13 | 14 | 14 | index=14255 | VLDFHNLPD( | K | T | 99.70% | | 1,467.78 | 3 | 0.003 | 2 | 45 | 57 |
| Audrey Caling | SCR | SCR | Service_Clair | Collagen alpł | CO5A1_HUM. | Collagen alpł | 100.00% | 0.51% | 183,544.70 | 13 | 14 | 14 | index=15549 | VLDFHNLPD( | K | R | 99.70% | Carbamidom | 2,362.18 | 4 | 0.006 | 2.6 | 45 | 65 |
| Audrey Caling | shARF6 | shARF6 | Service_Clair | Collagen alpł | CO5A1_HUM. | Collagen alpł | 100.00% | 0.30% | 183,544.70 | 6 | 7 | 7 | index=12195 | FGGGGDAGS | R | L | 99.70% | Oxidation (+1 | 2,861.39 | 3 | 0.013 | 4.4 | 526 | 554 |
| Audrey Caling | shARF6 | shARF6 | Service_Clair | Collagen alpł | CO5A1_HUM. | Collagen alpł | 100.00% | 0.30% | 183,544.70 | 6 | 7 | 7 | index=4590 | GLPGEPGPR | R | G | 99.60% | | 878.46 | 2 | 0.0014 | 1.6 | 673 | 681 |
| Audrey Caling | shARF6 | shARF6 | Service_Clair | Collagen alpł | CO5A1_HUM. | Collagen alpł | 100.00% | 0.30% | 183,544.70 | 6 | 7 | 7 | index=18740 | ILDEEVFEGDI | R | A | 99.70% | | 2,501.25 | 3 | 0.0091 | 3.6 | 211 | 231 |
| Audrey Caling | shARF6 | shARF6 | Service_Clair | Collagen alpł | CO5A1_HUM. | Collagen alpł | 100.00% | 0.30% | 183,544.70 | 6 | 7 | 7 | index=2115 | RSSKGPDVAY | R | V | 99.70% | | 1,234.64 | 3 | 0.0028 | 2.3 | 66 | 76 |
| Audrey Caling | shARF6 | shARF6 | Service_Clair | Collagen alpł | CO5A1_HUM. | Collagen alpł | 100.00% | 0.30% | 183,544.70 | 6 | 7 | 7 | index=3930 | SSKGPDVAYF | R | D | 99.70% | | 1,406.76 | 3 | 0.0039 | 2.8 | 67 | 79 |
| Audrey Caling | shARF6 | shARF6 | Service_Clair | Collagen alpł | CO5A1_HUM. | Collagen alpł | 100.00% | 0.30% | 183,544.70 | 6 | 7 | 7 | index=3922 | SSKGPDVAYF | R | D | 99.70% | | 1,406.75 | 4 | 0.003 | 2.1 | 67 | 79 |
| Audrey Caling | shARF6 | shARF6 | Service_Clair | Collagen alpł | CO5A1_HUM. | Collagen alpł | 100.00% | 0.30% | 183,544.70 | 6 | 7 | 7 | index=12921 | VLDFHNLPD( | K | T | 99.70% | | 1,467.78 | 3 | 0.0033 | 2.3 | 45 | 57 |

| Experiment na | Biological sar | Biological sar | MS/MS sampl | Protein group | Protein acces | Protein name | Protein identi | Protein perce | Protein molec | Number of un | Number of un | Number of tot | Spectrum nar | Peptide seque | Previous amir | Next amino ac | Peptide identi | Modifications | Actual peptid | Spectrum cha | Actual minus | Actual minus | Peptide start i | Peptide stop index |
|---|---|---|---|---|---|---|---|---|---|---|---|---|---|---|---|---|---|---|---|---|---|---|---|---|
| Audrey Caling | SCR | SCR | Service_Clair | Collagen alph | CO5A2_HUM | Collagen alph | 100.00% | 0.44% | 144,891.60 | 8 | 9 | 12 | index=5113 | GIVGMPGQR | R | G | 99.70% | Oxidation (+1 | 1,271.64 | 2 | 0.0026 | 2 | 993 | 1004 |
| Audrey Caling | SCR | SCR | Service_Clair | Collagen alph | CO5A2_HUM | Collagen alph | 100.00% | 0.44% | 144,891.60 | 8 | 9 | 12 | index=10413 | GLPGPLGPP | R | G | 99.70% | | 2,191.15 | 3 | 0.0077 | 3.5 | 798 | 821 |
| Audrey Caling | SCR | SCR | Service_Clair | Collagen alph | CO5A2_HUM | Collagen alph | 100.00% | 0.44% | 144,891.60 | 8 | 9 | 12 | index=5236 | GLTGNPGVQ | R | L | 99.70% | | 1,309.67 | 2 | 0.0043 | 3.3 | 573 | 586 |
| Audrey Caling | SCR | SCR | Service_Clair | Collagen alph | CO5A2_HUM | Collagen alph | 100.00% | 0.44% | 144,891.60 | 8 | 9 | 12 | index=5235 | GLTGNPGVQ | R | L | 99.70% | | 1,309.67 | 2 | 0.0043 | 3.3 | 573 | 586 |
| Audrey Caling | SCR | SCR | Service_Clair | Collagen alph | CO5A2_HUM | Collagen alph | 100.00% | 0.44% | 144,891.60 | 8 | 9 | 12 | index=4700 | GMPGERGRL | R | G | 99.70% | | 1,819.93 | 3 | 0.004 | 2.2 | 339 | 356 |
| Audrey Caling | SCR | SCR | Service_Clair | Collagen alph | CO5A2_HUM | Collagen alph | 100.00% | 0.44% | 144,891.60 | 8 | 9 | 12 | index=3064 | LGPQGAPGQ | R | G | 98.20% | | 979.52 | 2 | 0.0029 | 2.9 | 347 | 356 |
| Audrey Caling | SCR | SCR | Service_Clair | Collagen alph | CO5A2_HUM | Collagen alph | 100.00% | 0.44% | 144,891.60 | 8 | 9 | 12 | index=3065 | LGPQGAPGQ | R | G | 99.70% | | 979.52 | 2 | 0.0029 | 2.9 | 347 | 356 |
| Audrey Caling | SCR | SCR | Service_Clair | Collagen alph | CO5A2_HUM | Collagen alph | 100.00% | 0.44% | 144,891.60 | 8 | 9 | 12 | index=3986 | TCDDLKLCHS | R | Q | 96.90% | Carbamidom | 1,446.66 | 4 | 0.0037 | 2.6 | 1295 | 1306 |
| Audrey Caling | SCR | SCR | Service_Clair | Collagen alph | CO5A2_HUM | Collagen alph | 100.00% | 0.44% | 144,891.60 | 8 | 9 | 12 | index=5360 | VGDRGPAGP | R | G | 99.70% | | 3,630.68 | 4 | 0.0084 | 2.3 | 953 | 992 |
| Audrey Caling | SCR | SCR | Service_Clair | Collagen alph | CO5A2_HUM | Collagen alph | 100.00% | 0.44% | 144,891.60 | 8 | 9 | 12 | index=9595 | VGPPGPAGA | R | V | 99.70% | | 3,248.66 | 5 | 0.011 | 3.2 | 917 | 952 |
| Audrey Caling | SCR | SCR | Service_Clair | Collagen alph | CO5A2_HUM | Collagen alph | 100.00% | 0.44% | 144,891.60 | 8 | 9 | 12 | index=9594 | VGPPGPAGA | R | V | 99.70% | | 3,248.66 | 5 | 0.011 | 3.2 | 917 | 952 |
| Audrey Caling | SCR | SCR | Service_Clair | Collagen alph | CO5A2_HUM | Collagen alph | 100.00% | 0.44% | 144,891.60 | 8 | 9 | 12 | index=9586 | VGPPGPAGA | R | V | 99.70% | | 3,248.66 | 4 | 0.014 | 4.2 | 917 | 952 |
| Audrey Caling | shARF6 | shARF6 | Service_Clair | Collagen alph | CO5A2_HUM | Collagen alph | 100.00% | 0.21% | 144,891.60 | 5 | 5 | 5 | index=4568 | GIVGMPGQR | R | G | 99.70% | Oxidation (+1 | 1,271.64 | 2 | 0.0024 | 1.9 | 993 | 1004 |
| Audrey Caling | shARF6 | shARF6 | Service_Clair | Collagen alph | CO5A2_HUM | Collagen alph | 100.00% | 0.21% | 144,891.60 | 5 | 5 | 5 | index=4664 | GLTGNPGVQ | R | L | 99.70% | | 1,309.67 | 2 | 0.0041 | 3.1 | 573 | 586 |
| Audrey Caling | shARF6 | shARF6 | Service_Clair | Collagen alph | CO5A2_HUM | Collagen alph | 100.00% | 0.21% | 144,891.60 | 5 | 5 | 5 | index=2658 | LGPQGAPGQ | R | G | 98.90% | | 979.52 | 2 | 0.0025 | 2.6 | 347 | 356 |
| Audrey Caling | shARF6 | shARF6 | Service_Clair | Collagen alph | CO5A2_HUM | Collagen alph | 100.00% | 0.21% | 144,891.60 | 5 | 5 | 5 | index=4765 | VGDRGPAGP | R | G | 99.70% | | 3,630.68 | 4 | 0.0057 | 1.6 | 953 | 992 |
| Audrey Caling | shARF6 | shARF6 | Service_Clair | Collagen alph | CO5A2_HUM | Collagen alph | 100.00% | 0.21% | 144,891.60 | 5 | 5 | 5 | index=8623 | VGPPGPAGA | R | V | 98.60% | | 3,248.66 | 5 | 0.011 | 3.4 | 917 | 952 |

| Experiment na | Biological sar | Biological sar | MS/MS sampl | Protein group | Protein acces | Protein name | Protein identi | Protein perce | Protein molec | Number of un | Number of un | Number of tot | Spectrum nar | Peptide seque | Previous amir | Next amino ac | Peptide identi | Modifications | Actual peptide | Spectrum cha | Actual minus | Actual minus | Peptide start i | Peptide stop index |
|---|---|---|---|---|---|---|---|---|---|---|---|---|---|---|---|---|---|---|---|---|---|---|---|---|
| Audrey Caling | SCR | SCR | Service_Clair | Collagen alph | CO6A1_HUM | Collagen alph | 100.00% | 0.22% | 108,512.90 | 5 | 6 | 6 | index=1123 | FYREASSGAA | R | L | 99.70% | | 1,469.78 | 3 | 0.0028 | 1.9 | 926 | 938 |
| Audrey Caling | SCR | SCR | Service_Clair | Collagen alph | CO6A1_HUM | Collagen alph | 100.00% | 0.22% | 108,512.90 | 5 | 6 | 6 | index=4209 | GVFHQTVSR | R | K | 99.40% | | 1,029.54 | 3 | 0.0036 | 3.5 | 1015 | 1023 |
| Audrey Caling | SCR | SCR | Service_Clair | Collagen alph | CO6A1_HUM | Collagen alph | 100.00% | 0.22% | 108,512.90 | 5 | 6 | 6 | index=4214 | GVFHQTVSR | R | K | 99.70% | | 1,029.54 | 2 | 0.0031 | 3 | 1015 | 1023 |
| Audrey Caling | SCR | SCR | Service_Clair | Collagen alph | CO6A1_HUM | Collagen alph | 100.00% | 0.22% | 108,512.90 | 5 | 6 | 6 | index=8475 | LAERFLTAGR | R | V | 99.70% | | 2,024.06 | 4 | 0.0049 | 2.4 | 855 | 872 |
| Audrey Caling | SCR | SCR | Service_Clair | Collagen alph | CO6A1_HUM | Collagen alph | 100.00% | 0.22% | 108,512.90 | 5 | 6 | 6 | index=16081 | LLLFSDGNSC | R | A | 99.70% | | 1,931.01 | 2 | 0.0054 | 2.8 | 939 | 957 |
| Audrey Caling | SCR | SCR | Service_Clair | Collagen alph | CO6A1_HUM | Collagen alph | 100.00% | 0.22% | 108,512.90 | 5 | 6 | 6 | index=13638 | VFSVAITPDHI | K | L | 99.70% | | 1,579.84 | 3 | 0.0048 | 3 | 186 | 199 |
| Audrey Caling | shARF6 | shARF6 | Service_Clair | Collagen alph | CO6A1_HUM | Collagen alph | 100.00% | 0.21% | 108,512.90 | 4 | 5 | 5 | index=3723 | GVFHQTVSR | R | K | 98.70% | | 1,029.54 | 3 | 0.002 | 1.9 | 1015 | 1023 |
| Audrey Caling | shARF6 | shARF6 | Service_Clair | Collagen alph | CO6A1_HUM | Collagen alph | 100.00% | 0.21% | 108,512.90 | 4 | 5 | 5 | index=3730 | GVFHQTVSR | R | K | 99.70% | | 1,029.54 | 2 | 0.0031 | 3 | 1015 | 1023 |
| Audrey Caling | shARF6 | shARF6 | Service_Clair | Collagen alph | CO6A1_HUM | Collagen alph | 100.00% | 0.21% | 108,512.90 | 4 | 5 | 5 | index=14645 | LLLFSDGNSC | R | A | 99.70% | | 1,931.00 | 2 | 0.0047 | 2.4 | 939 | 957 |
| Audrey Caling | shARF6 | shARF6 | Service_Clair | Collagen alph | CO6A1_HUM | Collagen alph | 100.00% | 0.21% | 108,512.90 | 4 | 5 | 5 | index=4627 | VAVVQYSGTG | R | A | 99.70% | | 1,773.92 | 2 | 0.0088 | 5 | 873 | 888 |
| Audrey Caling | shARF6 | shARF6 | Service_Clair | Collagen alph | CO6A1_HUM | Collagen alph | 100.00% | 0.21% | 108,512.90 | 4 | 5 | 5 | index=12375 | VFSVAITPDHI | K | L | 99.70% | | 1,579.84 | 3 | 0.0052 | 3.3 | 186 | 199 |

| Experiment na | Biological sar | Biological sar | MS/MS sampl | Protein group | Protein acces | Protein name | Protein identi | Protein perce | Protein molec | Number of un | Number of un | Number of to | Spectrum nar | Peptide seque | Previous amir | Next amino ac | Peptide identi | Modifications | Actual peptid | Spectrum cha | Actual minus | Actual minus | Peptide start i | Peptide stop index |
|---|---|---|---|---|---|---|---|---|---|---|---|---|---|---|---|---|---|---|---|---|---|---|---|---|
| Audrey Caling | SCR | SCR | Service_Clair | Collagen alph | CO6A3_HUM | Collagen alph | 100.00% | 0.29% | 343,649.40 | 7 | 8 | 8 | index=18381 | AAPLQGMLP( | R | T | 99.00% | Oxidation (+1 | 1,632.94 | 2 | 0.0048 | 2.9 | 610 | 625 |
| Audrey Caling | SCR | SCR | Service_Clair | Collagen alph | CO6A3_HUM | Collagen alph | 100.00% | 0.29% | 343,649.40 | 7 | 8 | 8 | index=10733 | EFRELPNIEEF | R | I | 99.70% | | 1,430.72 | 3 | 0.0042 | 2.9 | 1598 | 1608 |
| Audrey Caling | SCR | SCR | Service_Clair | Collagen alph | CO6A3_HUM | Collagen alph | 100.00% | 0.29% | 343,649.40 | 7 | 8 | 8 | index=4603 | GMTQLQGTR | R | A | 99.70% | | 990.49 | 2 | 0.0024 | 2.5 | 2705 | 2713 |
| Audrey Caling | SCR | SCR | Service_Clair | Collagen alph | CO6A3_HUM | Collagen alph | 100.00% | 0.29% | 343,649.40 | 7 | 8 | 8 | index=16310 | IGDLHPQIVN | K | S | 99.70% | | 1,458.86 | 3 | 0.0031 | 2.1 | 1002 | 1014 |
| Audrey Caling | SCR | SCR | Service_Clair | Collagen alph | CO6A3_HUM | Collagen alph | 100.00% | 0.29% | 343,649.40 | 7 | 8 | 8 | index=13353 | NIDSEEVGKI/ | R | V | 99.70% | | 2,007.99 | 3 | 0.0059 | 2.9 | 1780 | 1798 |
| Audrey Caling | SCR | SCR | Service_Clair | Collagen alph | CO6A3_HUM | Collagen alph | 100.00% | 0.29% | 343,649.40 | 7 | 8 | 8 | index=1536 | SQPPPPQPAI | K | S | 99.70% | | 1,073.56 | 2 | 0.0027 | 2.5 | 3079 | 3088 |
| Audrey Caling | SCR | SCR | Service_Clair | Collagen alph | CO6A3_HUM | Collagen alph | 100.00% | 0.29% | 343,649.40 | 7 | 8 | 8 | index=5147 | VNHFVPEAGS | R | L | 99.70% | | 1,211.61 | 3 | 0.0014 | 1.1 | 1728 | 1738 |
| Audrey Caling | SCR | SCR | Service_Clair | Collagen alph | CO6A3_HUM | Collagen alph | 100.00% | 0.29% | 343,649.40 | 7 | 8 | 8 | index=5158 | VNHFVPEAGS | R | L | 99.70% | | 1,211.61 | 2 | 0.0026 | 2.1 | 1728 | 1738 |
| Audrey Caling | shARF6 | shARF6 | Service_Clair | Collagen alph | CO6A3_HUM | Collagen alph | 100.00% | 0.09% | 343,649.40 | 2 | 2 | 2 | index=4108 | GMTQLQGTR | R | A | 99.70% | | 990.49 | 2 | 0.0026 | 2.6 | 2705 | 2713 |
| Audrey Caling | shARF6 | shARF6 | Service_Clair | Collagen alph | CO6A3_HUM | Collagen alph | 100.00% | 0.09% | 343,649.40 | 2 | 2 | 2 | index=4157 | NADQEELVK | R | I | 99.70% | | 1,044.51 | 2 | 0.0032 | 3 | 1372 | 1380 |

Audrey Caling SCR_ARF6 010319 (7), Publication report created on 07/14/2025
Experiment: Audrey Caling SCR_ARF6 010319 (7)
- Peak List Generator: unknown
  - Version: unknown
  - Charge States Calculated: unknown
  - Deisotoped: unknown
  - Textual Annotation: unknown
- Database Set: 1 Database
  - Database Name: the Human 2018 database
    - Version: unknown
    - Taxonomy: All Entries
    - Number of Proteins: 20349
  - Does database contain common contaminants?: unknown
- Search Engine Set: 1 Search Engine
  - Search Engine: PEAKS Studio
    - Version: 10.0 (2018-11-21)
    - Samples: All Samples
    - Fragment Tolerance: 0.0100 Da (Monoisotopic)
    - Parent Tolerance: 10.0 PPM (Monoisotopic)
    - Fixed Modifications: +57 on C (Carbamidomethyl)
    - Variable Modifications: +1 on NQ (Deamidated), +16 on M (Oxidation), +42 on Peptide N-Terminal (Acetyl), +80 on STY (Phospho)
    - Database: the Human 2018 database (unknown version, 20349 entries)
    - Digestion Enzyme: Trypsin
    - Max Missed Cleavages: 3
    - Probability Model:
      - Service_Claing_010319_1 (peptides_1_1_0.mzid): Peptide Prophet with Delta Mass Correction (No decoys found) [all charge states]
      - Service_Claing_010319_2 (peptides_1_1_0.mzid): Peptide Prophet with Delta Mass Correction (No decoys found) [all charge states]
- Scaffold: Version: Scaffold_4.8.9
  - Modification Metadata Set: 2334 modifications
    - Source: C:\Program Files\Scaffold 4\parameters\unimod.xml
    - Comment:
  - Protein Grouping Strategy: Experiment-wide grouping with binary peptide-protein weights
  - Peptide Thresholds: 95.0% minimum
  - Protein Thresholds: 99.0% minimum and 1 peptide minimum
  - Peptide FDR: 0.3% (Prophet)
  - Protein FDR: 0.0% (Prophet)
  - GO Annotation Source(s):
  - Alternate ID Source(s): FASTA:UniProt/Swiss-Prot (UniProtKB)

DATABASE SEARCHING-- Tandem mass spectra were extracted by [unknown] version [unknown]. Charge state deconvolution and deisotoping were not performed. All MS/MS samples were analyzed using PEAKS Studio (Bioinformatics Solutions, Waterloo, ON Canada; version 10.0 (2018-11-21)). PEAKS Studio was set up to search the Human 2018 database (unknown version, 20349 entries) (only samples Service_Claing_010319_1 (peptides_1_1_0.mzid), "Service_Claing_010319_1 (peptides_1_1_0.mzid)", "Service_Claing_010319_1 (peptides_1_1_0.mzid)", Service_Claing_010319_2 (peptides_1_1_0.mzid), "Service_Claing_010319_2 (peptides_1_1_0.mzid)" and "Service_Claing_010319_2 (peptides_1_1_0.mzid)") assuming the digestion enzyme trypsin. PEAKS Studio was searched with a fragment ion mass tolerance of 0.0100 Da and a parent ion tolerance of 10.0 PPM. Carbamidomethyl of cysteine (only samples "Service_Claing_010319_1 (peptides_1_1_0.mzid)", "Service_Claing_010319_1 (peptides_1_1_0.mzid)", Service_Claing_010319_1 (peptides_1_1_0.mzid), "Service_Claing_010319_2 (peptides_1_1_0.mzid)", "Service_Claing_010319_2 (peptides_1_1_0.mzid)" and Service_Claing_010319_2 (peptides_1_1_0.mzid)) was specified in PEAKS Studio as a fixed modification. Deamidated of asparagine and glutamine (only samples Service_Claing_010319_1 (peptides_1_1_0.mzid), "Service_Claing_010319_1 (peptides_1_1_0.mzid)", "Service_Claing_010319_1 (peptides_1_1_0.mzid)", Service_Claing_010319_2 (peptides_1_1_0.mzid), "Service_Claing_010319_2 (peptides_1_1_0.mzid)" and "Service_Claing_010319_2 (peptides_1_1_0.mzid)"), oxidation of methionine (only samples "Service_Claing_010319_1 (peptides_1_1_0.mzid)", "Service_Claing_010319_1 (peptides_1_1_0.mzid)", Service_Claing_010319_1 (peptides_1_1_0.mzid), "Service_Claing_010319_2 (peptides_1_1_0.mzid)", "Service_Claing_010319_2 (peptides_1_1_0.mzid)" and Service_Claing_010319_2 (peptides_1_1_0.mzid)), acetyl of the n-terminus (only samples "Service_Claing_010319_1 (peptides_1_1_0.mzid)", Service_Claing_010319_1 (peptides_1_1_0.mzid), "Service_Claing_010319_1 (peptides_1_1_0.mzid)", "Service_Claing_010319_2 (peptides_1_1_0.mzid)", Service_Claing_010319_2 (peptides_1_1_0.mzid) and "Service_Claing_010319_2 (peptides_1_1_0.mzid)") and phospho of serine, threonine and tyrosine (only samples "Service_Claing_010319_1 (peptides_1_1_0.mzid)", Service_Claing_010319_1 (peptides_1_1_0.mzid), "Service_Claing_010319_1 (peptides_1_1_0.mzid)", "Service_Claing_010319_2 (peptides_1_1_0.mzid)", Service_Claing_010319_2 (peptides_1_1_0.mzid) and "Service_Claing_010319_2 (peptides_1_1_0.mzid)") were specified in PEAKS Studio as variable modifications.

CRITERIA FOR PROTEIN IDENTIFICATION-- Scaffold (version Scaffold_4.8.9, Proteome Software Inc., Portland, OR) was used to validate MS/MS based peptide and protein identifications. Peptide identifications were accepted if they could be established at greater than 95.0% probability by the Peptide Prophet algorithm (Keller, A et al Anal. Chem. 2002;74(20):5383-92) with Scaffold delta-mass correction. Protein identifications were accepted if they could be established at greater than 99.0% probability and contained at least 1 identified peptide. Protein probabilities were assigned by the Protein Prophet algorithm (Nesvizhskii, Al et al Anal. Chem. 2003;75(17):4646-58). Proteins that contained similar peptides and could not be differentiated based on MS/MS analysis alone were grouped to satisfy the principles of parsimony.
